# Development and characterisation of a promoter library for *Sulfolobus acidocaldarius*

**DOI:** 10.1101/2025.11.20.689296

**Authors:** Andries I. Peeters, Yifei Xu, Pauline Pijpstra, Peter J.T. Verheijen, Eveline Peeters, Brecht De Paepe, Marjan De Mey

**Affiliations:** Centre for Synthetic Biology, Ghent University, Ghent, Belgium; Research Group of Microbiology, Vrije Universiteit Brussel, Brussels, Belgium; Department of Biotechnology, Delft University of Technology, Delft, The Netherlands

## Abstract

The hyperthermoacidophilic archaeon *Sulfolobus acidocaldarius* is a promising production host for industrial biotechnology applications due to its ability to thrive in extreme conditions. However, the lack of well-characterised genetic parts and tools, particularly promoters, limits its potential for metabolic engineering. In this study, we developed the first promoter library for *S. acidocaldarius* by randomising specific regions of the core promoter sequence of P_Saci 2137_, a promoter known to function in both *S. acidocaldarius* and *Escherichia coli*. The library was initially screened in *E. coli* using mKate2, a red fluorescent reporter protein, and seven promoters were selected for characterisation in *S. acidocaldarius* using the thermostable β-galactosidase reporter LacS. The resulting promoter collection exhibited a 5-fold range of expression levels in *S. acidocaldarius*, spanning from low to high constitutive expression when compared to *S. acidocaldarius* promoters P_sac7d_ and P_malE_. This study demonstrates a successful workflow for generating and characterising *S. acidocaldarius* promoters, providing a valuable toolkit for fine-tuning gene expression and optimising metabolic pathways in this extremophilic archaeon. The design principles established here can be extended to other archaeal systems with similar promoter architectures.

## Introduction

The advancement of synthetic biology has revolutionised industrial biotechnology by enabling the rational design and engineering of microorganisms for diverse biotechnological applications [1, 2]. Central to synthetic biology is the development of robust molecular toolkits, which include essential genetic components such as promoters, ribosome binding site (RBS), and terminators [3–7]. Among these, promoters play a crucial role in controlling gene expression, allowing precise modulation of metabolic pathways in microbial cell factories [8]. The development of well-characterised promoter libraries is therefore critical for optimising gene expression and fine-tuning cellular functions to enhance bioproduction yields and efficiencies.

Promoters are fundamental DNA elements that regulate transcription by facilitating the recruitment of RNA polymerase (RNAP) and associated transcription factor (TF) [8]. In microbial synthetic biology, they serve as key regulatory elements to modulate gene expression levels, thereby influencing cellular metabolism, stress responses, and overall growth dynamics. The ability to engineer and characterise promoters with varying strengths enables researchers to construct optimised metabolic pathways for biotechnological applications, such as biofuel production, bioremediation, and pharmaceutical synthesis [9–13]. While extensive promoter libraries have been developed for model organisms such as *Escherichia coli* [14] and *Saccharomyces cerevisiae* [15], the availability of well-characterised promoters remains limited for non-model (extremophilic) microorganisms.

In recent years, there has been growing interest in exploring non-conventional microorganisms as production hosts, particularly those with unique properties that can offer advantages in industrial processes [16–19]. One such promising microorganism is *Sulfolobus acidocaldarius*, a member of the Thermoproteota. This hyperthermoacidophilic archaeon thrives in high-temperature (70-80 °C) and acidic (pH 2–3) environments, making it a highly attractive host for industrial biotechnology applications requiring robust metabolic performance under extreme conditions, as it can potentially withstand harsh process environments, reduce the risk of contamination and cut down cooling costs. Additionally, *S. acidocaldarius* offers other unique advantages, including thermostable enzymes and the ability to utilise various carbon sources, positioning it as a novel promising production host for biocatalysis and biomolecule synthesis [17, 19]. Despite its potential, the genetic toolbox for *S. acidocaldarius* remains underdeveloped, particularly in terms of well-characterised epxression elements such as promoters.

For the reasons stated above, *S. acidocaldarius* has been the subject of increasing research efforts to develop genetic tools and establish it as a viable production host. A set of genetic tools has been created for this organism, including methods for creating markerless deletion mutants, genomic tagging of genes, and ectopic integration of foreign DNA. These tools enable the construction of single, double, and triple deletion strains that can be complemented with pRN1-based expression vectors [20].

Currently, only a few promoters have been thoroughly characterised and utilised in *Sulfolobus acidocaldarius* (*S. acidocaldarius*), such as a maltose- (P_malE_ or P_Saci_1165_) [21], a copper- (PcopM or PSaci_0873), a xylose- (Pxyl or PSaci_1938) and an arabinose-inducible promoter (P_ara_ or P_Saci_2122_) [20, 22], and the strong constitutive promoters P_gdhA_ (or P_Saci_0155_) and P_sac7d_ (or P_Saci_0064_) [21]. While some promoters have been adapted for controlled gene expression, the lack of a comprehensive promoter library for *S. acidocaldarius* hinders the optimisation of gene expression and metabolic pathways in this extremophilic microorganism, limiting its potential as a production host for industrial biotechnology. To address this gap, the purpose of this study is to unlock the potential of this microorganism through the development of a promoter library, expanding the toolkit available for metabolic engineering and synthetic biology applications in this promising archaeal host.

## Results & discussion

Transcription is the first step in the expression of genes, and therefore the first level for tuning this expression. Promoters are the key regulatory DNA sequences that control transcription initiation by interacting with the RNAP and associated TFs. Transcription initiation frequencies of promoters are known to vary considerably, spanning over a 10,000-fold range, making them interesting targets for tuning gene expression levels [8].

Various strategies have been developed to modify promoter strengths. Some are based on combining existing promoter elements to form hybrid promoters, while others involve altering the actual promoter DNA sequence. These last type of strategies target different functional regions of the promoter by: i) mutating the entire sequence, ii) modifying the sequence or length of spacers, or iii) altering the promoter conserved boxes, either individually or in combination. From these three approaches, the second approach has been proven to have the highest success rate in bacteria as the others carry the risk of disrupting functionality since essential elements for recruiting RNAP are also targeted for mutagenesis [8, 14, 25, 26].

Although the promoter structure of Archaea differs significantly from that of bacteria, reflecting their evolutionary divergence and unique transcriptional machinery, we hypothesise that a similar engineering strategy can still be applied. However, high-throughput screening of large promoter libraries in *S. acidocaldarius* is currently not feasible. While a thermostable fluorescent reporter (TGP) has been developed for live imaging in *S. acidocaldarius* [27], it has not been shown to be functional for gene expression quantification. Instead, expression levels can only be assessed using the established thermostable β-galactosidase reporter (LacS). To overcome this limitation and enable the screening of larger *S. acidocaldarius* promoter libraries, we propose to first screen the promoter library in *E. coli* using a fluorescent reporter protein, and then evaluate a selected subset in *S. acidocaldarius* using the β-galactosidase reporter (Figure 1). This approach requires starting with a promoter sequence that functions in both *S. acidocaldarius* and *E. coli*. To this end we selected the promoter region, P_Saci_2137_, of a β-alanine aminotransferase gene Saci_2137 of *S. acidocaldarius* that was shown to exhibit transcription initiation in *E. coli* [23, 28, 29].

**Figure 1.**
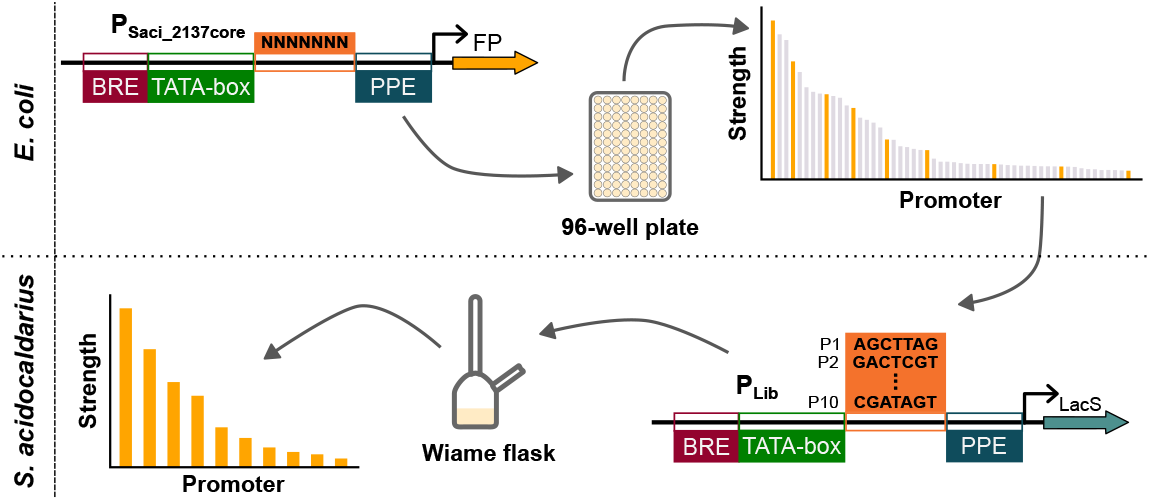
Overview of the workflow for the development of a promoter library for *S. acidocaldarius*. After randomisation of the P_Saci_2137_ promoter [23], the library is screened in *E. coli* using a fluorescent reporter protein. A subset of the library is then characterised in *S. acidocaldarius* using the LacS reporter, a thermostable β-galactosidase. BRE: B recognition element, PPE: proximal promoter element, FP: fluorescent protein. Genetic constructs are visualised following the Synthetic Biology Open Language standard [24].

First, the *S. acidocaldarius* P_Saci_2137_ was studied, focusing on identifying the core promoter sequence and eliminating any potential bacterial promoter elements. Following this, the consensus sequence for *S. acidocaldarius* promoters was determined using transcriptomics data to select the optimal region for variation in the promoter library design. Once designed, the promoter library was constructed and initially evaluated in *E. coli* using a red fluorescent reporter protein. A subset of this library was then tested in *S. acidocaldarius* using the thermostable β-galactosidase reporter LacS. Finally, these promoters were benchmarked against previously characterised *S. acidocaldarius* promoters (Figure 1).

### The *S. acidocaldarius* core promoter region of P_Saci_2137_ is still functional in *E. coli*

It is generally accepted that the core promoter region of Archaea is made up of a six-base pairs purine rich B recognition element (BRE), the TATA-box, which is an eight-base pair conserved region with consensus sequence TTTAWATR (W= A/T and R = A/G), and the proximal promoter element (PPE), an A/T-rich sequence located at - 12 to -1 from the transcription start site (TSS) [30]. The transcripts that these promoters produce are in most cases leaderless, meaning that there is no 5’ untranslated region (5’UTR), the TSS and the translation start site therefore coincide.

The Saci_2136-2137 region forms a divergent operon, where Saci_2136 encodes a TF that regulates the expression of both genes. Two promoters can be found in the intergenic region, one for each gene (Figure 2) [23]. Putative bacterial promoter -35 and -10 boxes were predicted by BPROM in this intergenic region, upstream of the BRE and TATA-box of the promoter region of Saci_2137 (P_Saci_2137_), possibly explaining the transciption promoting activity in *E. coli* (Figure 2) [31]. As we hypothesise that successful screening of a *S. acidocaldarius* promoter library in *E. coli* requires the transcription to be initiated by the same promoter elements, it was necessary to first assess whether transcription could still be initiated by the core P_Saci_2137_ region (P_Saci_2137core_) in *E. coli*. Therefore, P_Saci_2137core_ was defined starting from 20 base pairs upstream of the BRE (light blue arrow in Figure 2), not including the putative bacterial promoter boxes.

**Figure 2.**
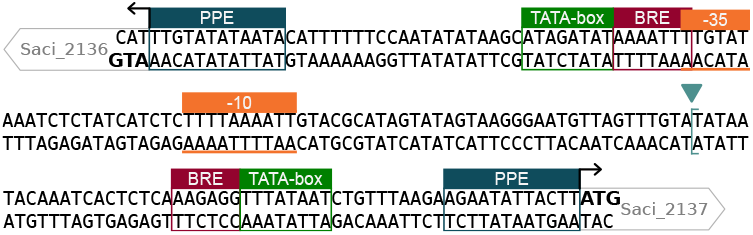
DNA sequence of the region between the Saci_2136 and Saci_2137 genes. B recognition element (BRE), proximal promoter element (PPE) and TATA-box are highlighted in red, green and blue respectively. The putative bacterial promoter -35 and -10 boxes are highlighted in between orange lines. The start of the core P_Saci_2137_ region is indicated with a light blue arrow and opening square bracket. The start codon is indicated in bold. The transcription start sites are indicated with an arrow showing the direction of transcription. The genes downstream of the promoter region are indicated in grey.

P_Saci_2137_ and P_Saci_2137core_ were cloned upstream of *mKate2*, coding for a red fluorescent protein,and their strengths in *E. coli* were characterised by measuring the fluorescence over ± 23 hours of growth. In *S. acidocaldarius*, as most promoters, P_Saci_2137_ lacks a 5’UTR and promotes the production of a leaderless transcript. To analyse these promoters as close to their natural function as possible, no 5’UTR, and thus no RBS, was added in the *E. coli* constructs. The strengths of these two promoters were compared to that of the *E. coli* ProB promoter, which is characterised as a promoter with medium strength (Figure 3) [32]. It is important to note that the ProB promoter was characterised in the presence of a 5’UTR upstream of the *mKate2* gene.

**Figure 3.**
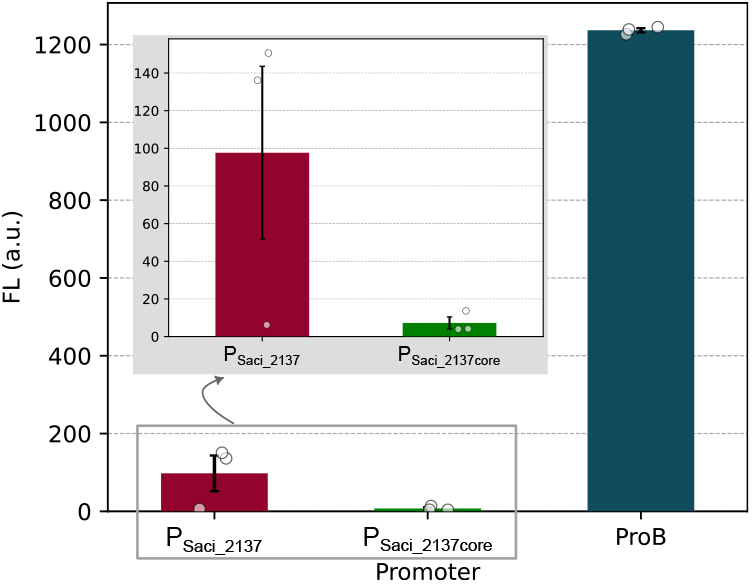
Mean mKate2 expression values of the *Sulfolobus acidocaldarius* promoters P_Saci_2137_ and P_Saci_2137core_, and the *E. coli* ProB promoter in *E. coli*. Fluorescent values were obtained in the stationary phase and normalised for cell growth determined by optical density at 600 nm (OD_600_). The error bars represent the standard error of the mean of three replicates. The dots represent the expression values of the individual replicates. FL: Fluorescence (arbitrary units (a.u.)).

From Figure 3 it is clear that the strengths of both P_Saci_2137_ and P_Saci_2137core_ are lower than that of the ProB promoter. This is not surprising, especially taking into account that the ProB promoter sequence includes an RBS, which is not the case for the P_Saci_2137_ promoter variants. Contrary to *S. acidocaldarius*, where leaderless genes are the norm, in *E. coli* most promoters are indeed followed by a 5’UTR. It is therefore not surprising that the absence thereof results in promoters with lower expression strengths. When comparing P_Saci_2137_ and P_Saci_2137core_, deleting the putative -35 and -10 boxes from the sequence had a great impact on the promoter strength. So much so, that the fluorescence expression from P_Saci_2137core_ in *E. coli* was not significantly different from the wild-type *E. coli* strain, which was used to correct the data (see Equation 1) (one tailed one-sample t-test, α = 0.05, p-value = 0.076), albeit only by a small margin. For the screening strategy (Figure 1) to work, it is important that the transcription in *E. coli* is driven from the same promoter elements as in *S. acidocaldarius*. Therefore, designing the promoter library starting from P_Saci_2137_ would not make sense and P_Saci_2137core_ was selected for developing the *S. acidocaldarius* promoter library.

### Construction of a promoter library for *S. acidocaldarius* by randomising the core promoter region of P_Saci 2137_

To design the promoter library, first a sequence logo [33] was generated from 2305 *S. acidocaldarius* promoter regions, spanning from -80 to +20 relative to the TSSs, which were determined from transcriptomic data [34]. The resulting logo (Figure 4) reveals that the conserved regions in *S. acidocaldarius* promoters align with the three archaeal core promoter regions (BRE, TATA-box and PPE). The region between the TATA-box and PPE is relatively unconserved in *S. acidocaldarius*, making it an ideal target for randomisation.

**Figure 4.**
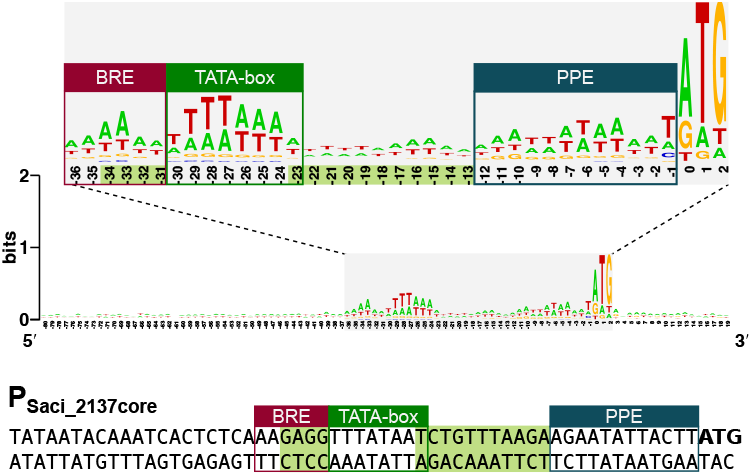
**TOP:** Sequence logo of promoter regions in *S. acidocaldarius*. B recognition element (BRE) in red, TATA-box in green and proximal promoter element (PPE) in blue. In light green the positions are highlighted that are not conserved and therefore ideal targets for randomisation in promoter engineering. **BOTTOM:** The P_Saci_2137core_ promoter sequence. BRE in red, TATA-box in green and PPE in blue. In light green the positions are highlighted that were selected for randomisation in the promoter library. The start codon is indicated in bold.

Additionally, it has been demonstrated that modifying the last four base pairs of the BRE can alter the promoter’s expression level without disrupting its function [35]. Therefore, the regions selected for randomisation include the last four base pairs of the BRE, as well as the region between the TATA-box and PPE, including the final base pair of the TATA-box, which also shows low conservation in *S. acidocaldarius* (see Figure 4 - positions highlighted in green).

The promoter library design was applied to the P_Saci_2137core_ promoter as shown in Figure 4. Degenerated oligonucleotides were used to randomise the selected promoter regions in P_Saci_2137core_, upstream of *mKate2. E. coli* was then transformed with the resulting promoter library and 188 randomly selected transformants were screened for promoter strength.

Figure 5 shows the fluorescence expression of the *S. acidocaldarius* promoter library in *E. coli*. While many promoters in the library exhibited no mKate2 expression, 57 variants displayed mKate2 expression levels higher than the background fluorescence. This supports the assumption that randomising P_Saci_2137core_, using the promoter library design for *S. acidocaldarius*, can generate functional promoters with a broad range of strengths. Among these 57 promoter variants, the promoter strengths spanned a 7169-fold range.

**Figure 5.**
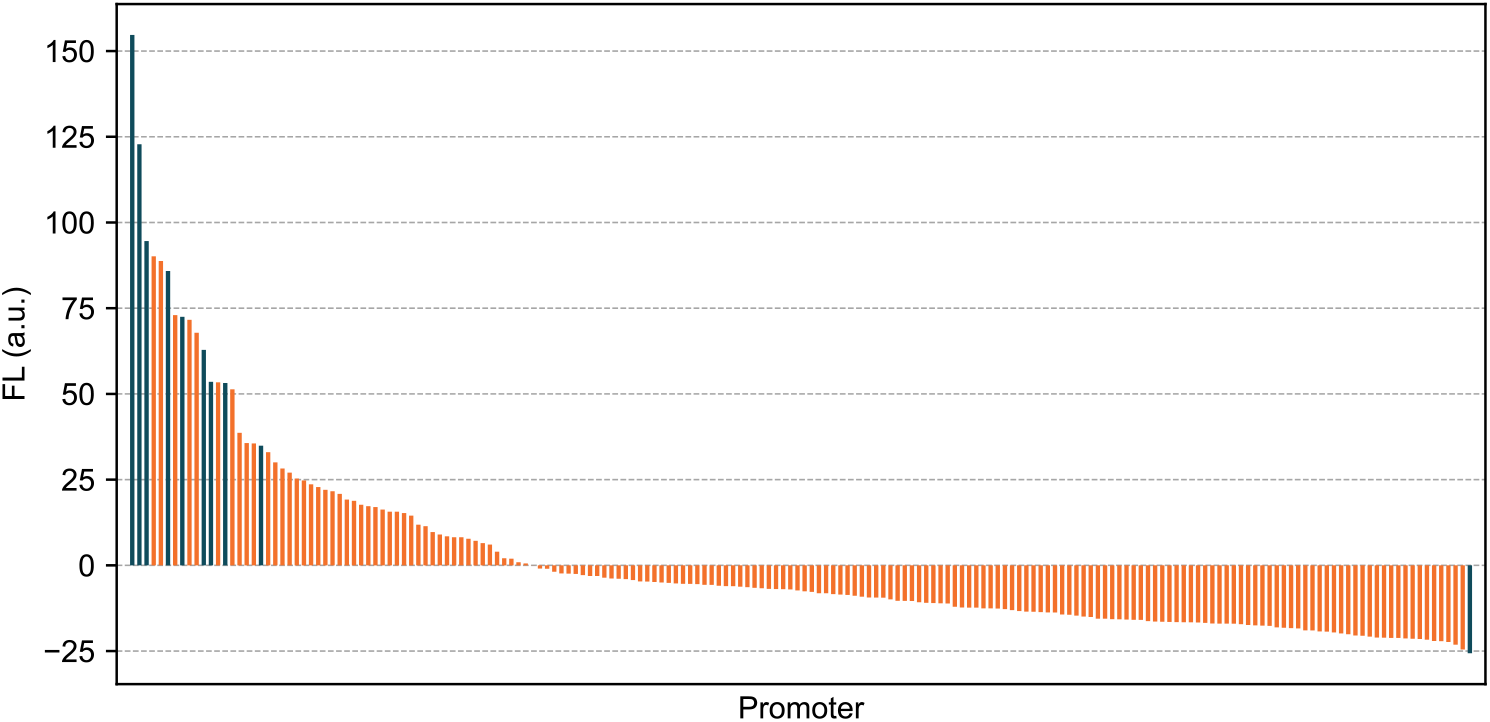
mKate2 expression values of the *S. acidocaldarius* P_Saci_2137core_ promoter library in *E. coli*. Blue bars represent the promoter variants selected for further characterisation. Fluorescence values were obtained in the stationary phase. Negative fluorescence values can be explained by the correction of the normalised fluorescence values with the mean normalised fluorescence of the reference strain (see Equation 1). FL: Fluorescence (arbitrary units (a.u.)).

Next, ten promoter variants were selected (Figure 5 in blue) for further characterisation. Variants were selected based on their fluorescence expression levels, aiming to cover a broad range of strengths, while focussing on promoters with higher expression levels. The promoter with the lowest expression level was also selected because the hypothesis that these promoters will exhibit comparable strengths in *S. acidocaldarius* is not guaranteed.

### Evaluation of the *S. acidocaldarius* promoter library

The ten selected promoters were characterised in depth, first in *E. coli* and next in *S. acidocaldarius*. We refer to these promoters by ‘P_Ec_’, followed by the number of the ranked promoter according to its strength in *E. coli*. When tested in *S. acidocaldarius*, this name is extended with ‘Sa’, followed by the number of the ranked promoter according to its strength in *S. acidocaldarius*.

The promoter collection was characterised by analysing mKate2 expression in *E. coli*. The resulting fluorescence values showing the promoter strengths are depicted in Figure 6, together with those of *S. acidocaldarius* promoters P_Saci_2137_ and P_Saci_2137core_. In *E. coli*, the promoter collection spans an 18-fold range of fluorescence expression. When compared to the reference promoters P_Saci_2137_ and P_Saci_2137core_, most promoters from the collection have a strength between those of the *S. acidocaldarius* promoters. Only P_Ec1_ seems to outperform the original P_Saci_2137_ promoter region.

As there are no thermostable fluorescent proteins for the analysis of promoter strengths in *S. acidocaldarius*, the promoter strengths were analysed using the thermostable β-galactosidase LacS reporter from *Saccharolobus solfataricus*.Each of the ten selected promoter variants was cloned upstream of the *lacS* gene in a pRN1-based vector with the *pyrEF* genes for uracil auxotrophy (based on pSVA1450 [36], without the *malR* transcription unit). *S. acidocaldarius* was transformed with these constructs and grown in triplicate for ± 24 hours, until an OD_600_ was reached of around 0.4. The cultures were then sampled and a β-galactosidase activity assay was performed on the samples, in triplicate. The P_Saci_2137_ and P_Saci_2137core_ promoters were also analysed in this way. To assess the applicability of this new promoter collection for *S. acidocaldarius*, it was benchmarked to two well-characterised *S. acidocaldarius* promoters from literature. The promoter of the *sac7d* gene (Saci_0064), P_sac7d_, encoding an abundant DNA-binding protein, was chosen as it has shown to be a very strong constitutive promoter in *S. acidocaldarius* [21]. The malE (Saci_1165) promoter from *S. acidocaldarius*, P_malE_, is a maltose-inducible promoter that has been thoroughly characterised and used for the overexpression of genes in *S. acidocaldarius* [20–22, 36]. In induced state, the native P_malE_ is 5-fold weaker, compared to P_sac7d_ [21]. In this study we used pSVA1450 as a reference, which harbours a mutated version of P_malE_ developed by Wagner *et al*. (P_malE pSVA1450_), increasing its strength 4-fold compared to pCmalLacS which contains the native P_malE_ [21, 36]. pSVA1450 harbours a copy of the *malR* transcription unit coding for the activator of P_malE_. As an extra control, we also included pJL1601, a variant of pSVA1450, where the copy of *malR* is removed from the vector (P_malE pJL1601_). The malE promoters were characterised in their leaky, uninduced state and are therefore considered weak promoters.

**Figure 6.**
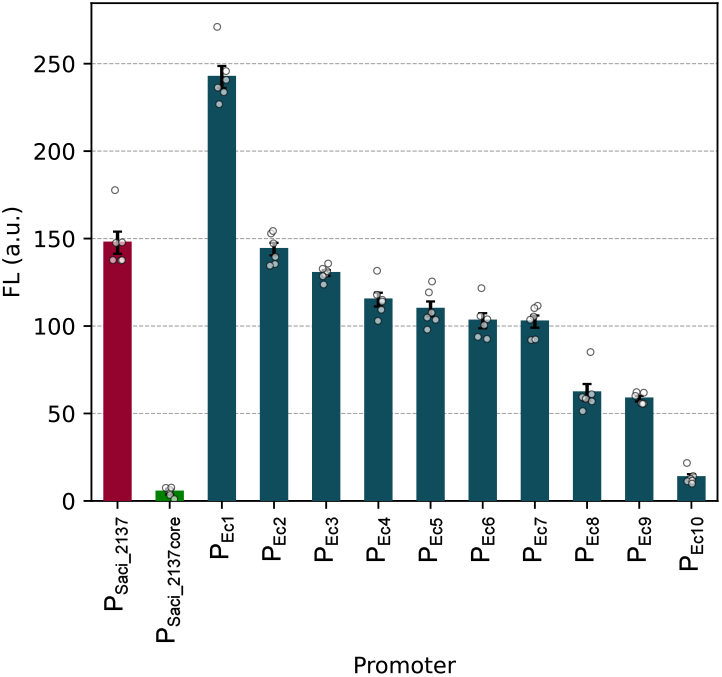
Mean mKate2 expression values of the ten selected promoter variants of the P_Saci_2137core_ promoter library and, P_Saci_2137_ and P_Saci_2137core_ as references, in *Escherichia coli*. The error bars represent the standard error of the mean of six biological replicates. The dots represent the expression values of the individual replicates. FL: Fluorescence (arbitrary units (a.u.)).

The results of the *S. acidocaldarius* promoter library characterisation are shown in Figure 7. We were not able to get any stable transformants for three of the ten selected promoters (P_Ec1_, P_Ec2_ and P_Ec5_), so only seven promoters were characterised in *S. acidocaldarius*. The promoter collection shows a 5-fold range of expression levels, which is in the same range as the reference promoters. P_Ec10 Sa1_ is the strongest promoter, with an expression level comparable to that of P_sac7d_ and P_Saci_2137core_. It is interesting that P_Saci_2137core_ has such a high expression level and that this level is ±2.5-fold higher than the full Saci_2137 promoter region (P_Saci_2137_). Especially, considering that previous research showed that a differently truncated P_Saci_2137_ promoter (106 base pairs from the TSS, versus 56 base pairs in P_Saci_2137core_) had lower expression compared to P_Saci_2137_ [28]. The promoter with the lowest expression level, P_Ec3 Sa7_, has a level similar to that of P_malE pJL1601_. Interestingly, from the promoters tested in *S. acidocaldarius*, P_Ec3 Sa7_ showed the highest expression level in *E. coli*. In contrast, P_Ec10 Sa1_, which had the lowest expression level in *E. coli*, is the strongest performing one in *S. acidocaldarius*.

**Figure 7.**
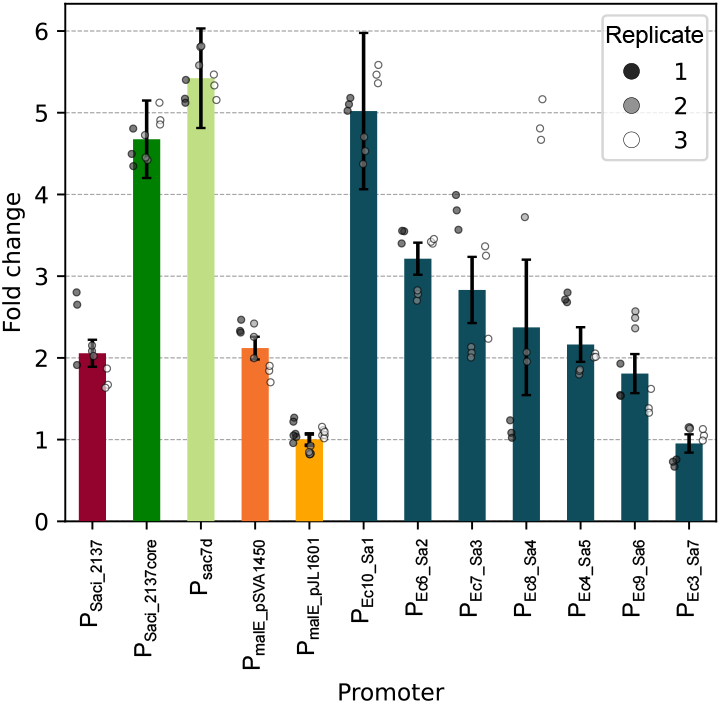
Fold changes, compared to P_malE_pJL1601_, of the mean LacS expression values of the selected promoter variants of the PS_aci_2137core_ promoter library, and P_Saci_2137_, P_Saci_2137core_, P_sac7d_, P_malE-pSVA1450_ and P_malE_pJL1601_ as references, in *S. acidocaldarius*. The error bars represent the standard error of the mean LacS expression values. The dots represent the expression values of individual technical replicates, while the colour of the dots indicates the biological replicate.

The sequences of the promoter library (Figure 8) are all distinct from each other. Looking at the randomised regions, we can see that there is only conservation in the last base pair of the TATA box. On this position, an adenine is favoured, while a thymine is also an option. There is also not a clear trend in the last four base pairs in the BRE. The BRE is a purine-rich six base pair DNA-sequence, and it has been shown to have a great influence on the promoter strength [30, 37, 38]. Table 1 contains the purine contents of the BRE of the library promoters. All promoters have a BRE purine content of three to five, which is lower than the six purine bases in the P_Saci_2137core_ promoter. However, in this collection higher purine content of the BRE does not result in higher promoter strength. P_Ec9 Sa6_ and P_Ec3 Sa7_, which are the promoters with the lowest expression levels have a purine content of five, compared to three or four for the rest of the collection, which are stronger promoters. The spacer region between the TATA-box and PPE is also very random. Here, the purine contents (Table 1) in the library promoters range from three to nine, while the P_Saci_2137core_ promoter spacer has a purine content of five, and do not seem to correlate with the promoter strength either. The BRE and spacer were randomised simultaneously, it is therefore not possible to determine the influence of the BRE and spacer on the promoter strength separately. However, the results show that the randomisation of these regions resulted in a library with a 5-fold range in promoter strengths. This collection contains promoters with a range of strengths, which can be used to fine-tune gene expression levels in *S. acidocaldarius*.

**Table 1.** Purine content of the B recognition element (BRE) and spacer between the TATA-box and proximal promoter element of the promoter collection selected from the P_Saci_2137core_ promoter library.

|  | BRE (n/6) | Spacer (n/10) |
| --- | --- | --- |
| $P_{Saci\_2137core}$ | 6 | 5 |
| $P_{Ec10\_Sa1}$ | 4 | 8 |
| $P_{Ec6\_Sa2}$ | 3 | 6 |
| $P_{Ec7\_Sa3}$ | 3 | 8 |
| $P_{Ec8\_Sa4}$ | 4 | 7 |
| $P_{Ec4\_Sa5}$ | 3 | 6 |
| $P_{Ec9\_Sa6}$ | 5 | 9 |
| $P_{Ec3\_Sa7}$ | 5 | 3 |

**Figure 8.**
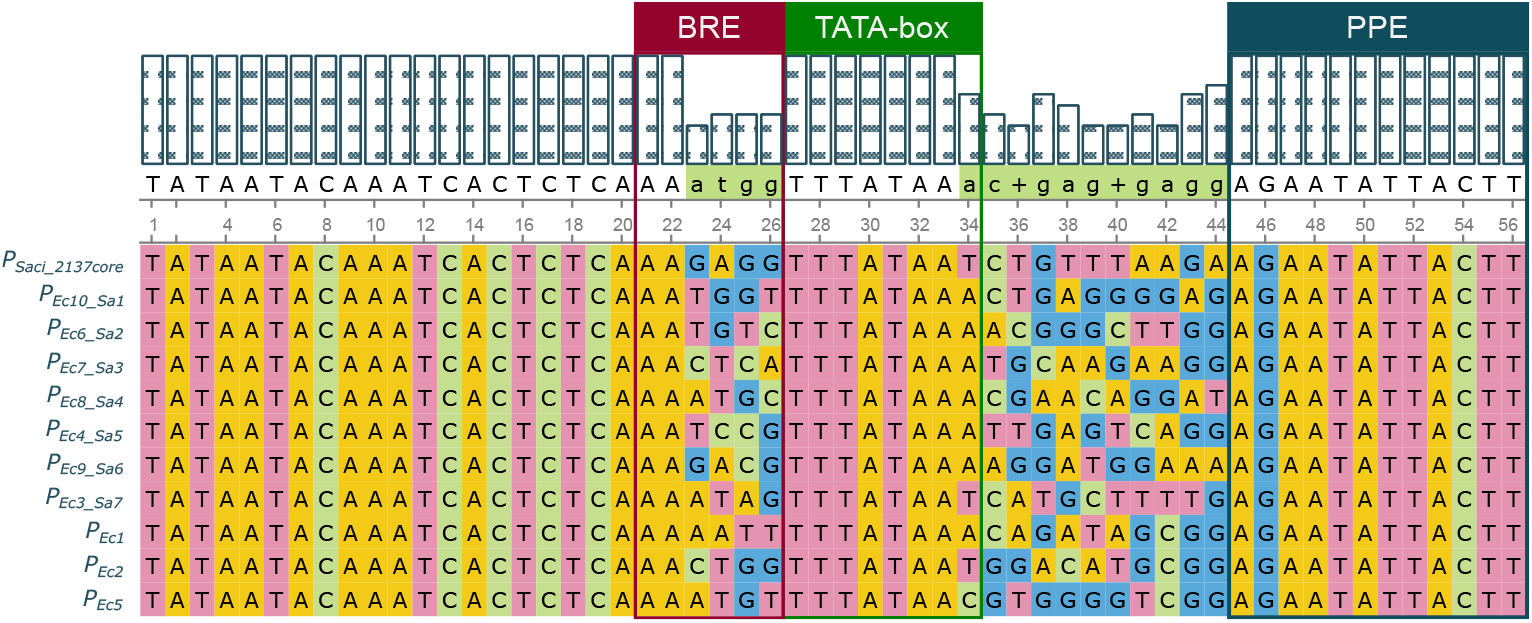
Clustal Omega alignment of the promoter sequences of P_Saci_2137core_ and the promoter collection selected from the P_Saci_2137core_ promoter library. B recognition element (BRE) in red, TATA-box in green and proximal promoter element (PPE) in blue. In light green the positions are highlighted that were selected for randomisation in the promoter library.

## Conclusions

We created the first-ever promoter library for *S. acidocaldarius*. The library was designed based on the P_Saci_2137core_ promoter and specific regions were selected for randomisation. To evaluate the library, the promoters were first screened in *E. coli* using a red fluorescent protein. A subset of the library was then characterised in *S. acidocaldarius* using the thermostable β-galactosidase reporter LacS. The promoter strengths span a 5-fold range, corresponding to very high to low constitutive expression when compared to known *S. acidocaldarius* promoters (P_sac7d_ and P_malE pJL1601_). With this promoter library, we are now able to fine-tune gene expression levels in *S. acidocaldarius*, which is essential for optimising metabolic pathways and enhancing bioproduction yields in industrial biotechnology applications.

The workflow developed in this study, based on starting from a *S. acidocaldarius* promoter with activity in *E. coli* to screen a promoter library in a high-throughput way, using a fluorescent reporter protein, and then characterising a subset of the library in *S. acidocaldarius* using LacS as reporter, is not flawless. It is clear from the results that the promoter expression levels in *E. coli* do not correlate with the levels in *S. acidocaldarius*. Even stronger, for the strongest and weakest promoter an inverse correlation could be observed. Although this is not surprising, as the transcription machinery in bacteria and archaea differ significantly, it is interesting to note that all promoters that were selected for further characterisation, and successfully tested in *S. acidocaldarius*, resulted in expression of the reporter protein. This shows that the randomisation of the selected regions in the P_Saci_2137core_ promoter resulted in functional promoters.

The sequence logo from transcriptomic data showed that the *S. acidocaldarius* promoter regions closely match the archaeal core promoter architecture. Randomising the less conserved region between the TATA-box and PPE together with the last four base pairs of the BRE resulted in a library with a 5-fold range in promoter strengths. This design for promoter libraries in *S. acidocaldarius* was very successful, resulting in promoters with both high, medium and low expression levels, while only characterising seven promoter variants. The design principles used in this study can be applied to other *S. acidocaldarius* promoters, and potentially to other archaea with a similar core promoter architecture.

## Materials & methods

All materials were obtained from Sigma-Aldrich (Overijse, Belgium), unless stated otherwise. All media were sterilised by autoclaving at 121°C for 30 minutes at 1 bar, unless specified differently. Standard molecular biology methods were applied as described by Green [39].

### Strains, media and growth conditions

Cloning was carried out using *Escherichia coli* (*E. coli*) TOP10 cells (Invitrogen, Carlsbad, USA). Lysogeny broth (LB) medium, containing 10 g/L bacto-tryptone, 5 g/L yeast extract (both Difco, Erembodegem, Belgium), and 5 g/L NaCl, was used for culturing *E. coli*. To prepare LB agar plates, 12 g/L agar (Difco, Erembodegem, Belgium) was added to the medium. When necessary, growth media were supplemented with antibiotics at the following concentrations: ampicillin (100 µg/mL), kanamycin (50 µg/mL), or chloramphenicol (25 µg/mL). *E. coli* strains were incubated at 30 °C with shaking at 200 rotations per minute (rpm) on a shaker with a 5 cm amplitude (LS-X AppliTek, Nazareth, Belgium).

*S. acidocaldarius* SK-1 [40] was grown in Brock basal salts medium [41] with addition of 0.2% sucrose and 0.1% NZ-amine, and adjusted to pH 3 with sulfuric acid. Uracil (20 µg/mL) was added when cells were grown under non-selective conditions. For solid culture, a 2x Brock basal salts solution containing 0.2% NZ-amine and 0.4% sucrose was used, supplemented with 6 mM CaCl_2_ and 20 mM MgCl_2_. After sterilisation, this solution was mixed with the same volume of 1.2% Gelrite solution (Labconsult, Schaerbeek, Belgium) at 70° C and poured quickly to prevent premature solidification. *S. acidocaldarius* was cultured at 75 °C and 180 rpm in a New Brunswick Innova 44 incubator with a 5 cm orbit (Eppendorf, Aarschot, Belgium).

### Transformation protocols

*E. coli* Top10 cells were made electrocompetent using the glycerol/mannitol density gradient centrifugation method [42]. 50 µL aliquots were stored at -80 °C for later use. 2 µL of circular polymerase extension cloning (CPEC) mix was added to the electrocompetent cells, unless stated differently. Electroporating was carried out at 2.5 kV, 200 Ω and 25 µF in a pre-chilled electroporation cuvette (Bio-Rad, Lokeren, Belgium) with a gap width of 2 mm. Immediately after electroporation, 950 µL of LB was added and the cells were incubated for 2 hours at 30 °C before 100 L of the cell mixture was plated onto selective LB-agar. *S. acidocaldarius* cells were transformed as described by Schleper *et al*. [43] with 200 ng of plasmid DNA. The entire transformation mixture was plated onto selective solid medium. Plates were incubated for 5 days at 75 °C. The plates were then sprayed with a 5 mg/mL X-gal in NN– dimethylformamide solution and incubated at 75°C for 30 minutes. Colonies with high β-galactosidase activity turned blue.

### Plasmid construction

All plasmids in this chapter were constructed using CPEC [44] with Q5^®^ High-Fidelity DNA polymerase (New Englands Biolabs, Ipswich, MA, USA). Oligonucleotides and double-stranded DNA fragments were ordered from Integrated DNA Technologies (Leuven, Belgium). The DNA sequence of all constructed plasmids was verified by Sanger sequencing (Macrogen, Amsterdam, The Netherlands). The plasmids used in this study are listed in Supplementary Table 1 and the plasmid maps can be found in Supplementary Tables 2-6.The *E. coli* plasmids were all based on the pBR322 backbone with a kanamycin antibiotic resistance gene. The in-house plasmid pBR322-ProB was used for mKate2 expression driven by the ProB promoter [32]. The same plasmid was also used as template to amplify the pBR322 backbone and *mKate2* gene. The P_Saci_2137_ and P_Saci_2137core_ promoters were amplified from the pUC18-V1 plasmid developed by Liu [28] and combined with the pBR322 backbone and *mKate2* for the construction of pBR322-P_Saci_2137_ and pBR322-P_Saci_2137core_. The *E. coli* promoter library (pBR322-P_EcLib_) was constructed by CPEC with a degenerated primer 5’-TAATACAAATCACTCTCAAANNNNTTTATANNNNNNNNNNNAGAATATTACTTATGGTTAGCG-3’ and the plasmid backbone of pBR322PSaci_2137core. PEc1-10 plasmids (pBR322-PEc1-10) were harvested from the corresponding *E. coli* library strains.

All *S. acidocaldarius* plasmids were constructed with the pRN1 backbone and *pyrEF* genes for uracil auxotrophy based selection. pSV1450 [36] and pJL1601, harbouring P_malE pSVA1450_ and P_malE pJL1601_ respectively, were kindly provided by the MICR research group (Vrije Universiteit Brussel, Brussels, Belgium). To construct the *S. acidocaldarius* plasmids, the pRN1 backbone and *LacS* gene were amplified from pJL1601. Next, primers, containing the sequences of the promoter collection variants (Supplementary Table 6), were ordered and the *S. acidocaldarius* promoter collection plasmids (pRN1-P_Ec10 Sa1-Ec3 Sa7_) were constructed by CPEC with the pRN1 backbone and *LacS*. The P_sac7d_ sequence was ordered as a double-stranded DNA fragment with overhangs and combined with the pRN1 backbone and *LacS* gene by CPEC to construct pRN1-P_sac7d_.

### Sequence logo generation

To generate the sequence logo of *S. acidocaldarius* promoter regions, the TSSs were determined from transcriptomic data [34]. Next the promoter regions, spanning from -80 to +20 relative to the TSSs, were extracted from the *S. acidocaldarius* genome, taking into account the orientation of the transcript. With these sequences the sequence logo was generated using the WebLogo tool [33].

### *In vivo* fluorescence measurements

*E. coli* transformants were inoculated in 150 L of LB medium with kanamycin in a transparent flat-bottomed 96-well plate (Greiner Bio-One, Vilvoorde, Belgium). After growth overnight on a Compact Digital Microplate shaker (Thermo Fisher Scientific, Massachusetts, USA) at 30 °C and 800 rpm, the cultures were diluted 300x in an assay plate (a black flat-bottomed 96-well plate (Greiner Bio-One, Vilvoorde, Belgium)) to a volume of 150 L. All plates were sealed with a Breathe-Easy^®^ sealing membrane. For the promoter library screening, 188 randomly selected transformants were selected and the assay plate was incubated on a Compact Digital Microplate shaker (Thermo Fisher Scientific, Massachusetts, USA) at 30 °C and 800 rpm. The OD_600_ and mKate2 expression (fluorescence emission at 635 nm after excitation at 588 nm) were measured at the end of the 24 hours in a Tecan M200 Pro plate reader (Tecan, Benelux, Mechelen, Belgium). For the P_Saci_2137_/P_Saci_2137core_ characterisation in *E. coli*, three replicates were tested per strain, while six replicates were tested for the promoter collection characterisation in *E. coli*. The assay plate of the two characterisation experiments was incubated in a Tecan M200 Pro plate reader (Tecan, Benelux, Mechelen, Belgium) at 30 °C while shaking at an orbit of 2 mm. The OD_600_ and mKate2 expression (fluorescence emission at 635 nm after excitation at 588 nm) were measured every 15 minutes for 24 hours.

### β-galactosidase activity assay

*S. acidocaldarius* strains were cultured in 20 mL of Brock basal salts medium supplemented with 0.2% sucrose and 0.1% NZ-amine in 100 mL Wiame flasks in triplicate for ± 24 hours until the OD_600_ reached around 0.4. 2 mL of culture was sampled for the β-galactosidase activity assay, which was conducted following van der Kolk *et al*. [22], with modifications. The conversion rate of o-nitrophenol-β-D-galactopyranoside (ONPG) was monitored at 410 nm using a microplate reader (Infinite M Nano+, TECAN, Männedorf, Switzerland) every 5 minutes while incubated at 42 °C for 4 hours. The β-galactosidase activity, serving as a proxy for LacS expression, was determined from the slope of the conversion curves. Each sample was analysed in triplicate.

### Promoter sequence alignment

The promoter sequences of P_Saci_2137core_ and the promoter collection were aligned using Clustal Omega [45]. The alignment was visualised using Unipro UGENE [46].

### Data processing and statistical testing

Data analysis was performed in Python using the pandas package (https://pandas.pydata.org/) unless stated differently. Generative artificial intelligence was used to generate and debug python code.

#### Fluorescence measurements

The fluorescence measurements (FP) were normalised for growth by dividing by the OD_600_ values. At each time point, the normalised fluorescence was corrected by the mean normalised fluorescence of the wild type *E. coli* strain with no plasmid (see Equation 1). The corrected fluorescence values were then averaged over the replicates. Standard errors (SEs) were determined through proper error propagation.

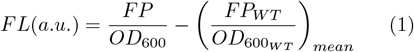

#### β-galactosidase activity assay

To determine the slopes, a Monod-type model with smoothing was fitted to the conversion data:

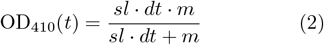

where

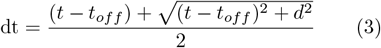

Here, *sl* represents the slope, *m* the maximum value, *t* the time, *t*_*off*_ the offset, and *d* the period around the offset in which the transition takes place. The slope was obtained by fitting the model to data points ranging from t_0_-t_max_ to t_0_-t_5_ Local minima of the fit error of these fits were then searched and from those fits, the slope of the fit resulting in the slope with the lowest standard error was selected. If the resulting fit was inadequate, the average fitted d value of the other technical and/or biological replicates was used as a fixed value for subsequent fitting. The geometric mean of the slopes was then calculated, along with the geometric standard error. If the fit error or technical replicate error exceeded the geometric standard error, the larger value was used instead. The fold change was calculated by dividing the geometric mean of the slopes of the promoter variants by the geometric mean of the slopes of the reference promoter P_malE pJL1601_. Errors were scaled accordingly. The parameters determined by the model fitting can be found in Supplementary Table 7. The graphs showing the estimated slopes and fit errors can be found in Supplementary Figures 1-39.

#### Statistical testing

A one tailed one-sample t-test was performed to test whether the mean fluorescence of P_Saci_2137core_ was significantly different from the background. This statistical test was performed with a significance level of α= 0.05.

## Acknowledgements

The authors would like to thank Stijn Bovijn, Siel Goethals and Glen Herremans for their contributions to the *Escherichia coli* screening experiments. Additionally, we would like to thank David Sybers for his help and training in working with *Sulfolobus acidocaldarius*. Generative artificial intelligence was used for improving the structure and flow of the written text.

## Author contributions

Peeters I. A., Peeters E., De Paepe B. and De Mey M. were involved in the conceptualisation of the research. The experimental work was performed by Peeters A., Pijpstra P. and Xu Y. Verheijen P. established the models and script for the β-galactosidase assay data processing. Peeters A. wrote the manuscript that was critically reviewed by De Mey M. and De Paepe B.

## Funding

The research presented in this article was supported by the FWO research project Sulfofact (FWOOPR2018007301), the Moonshot cSBO project TACBIO (HBC.2020.2618), the BOF iBOF project POSSIBL (BOF iBOF BOF20/IBF/131) and the Vrije Universiteit Brussel (Strategic Research Program SRP91). Xu Y. was supported by a PhD scholarship from the Chinese Scholarship Council. We would also like to thank the UGent Core Facility HTS for SynBio for training, support and access to the instrument park. This Core Facility is supported by UGent-BOF through grants BOF/COR/2022/002, BAS018-18, BAS020-131 and BOF/BAS/2022/114 as well as FWO grants I011118N and I000925N.

## Data availability

All data presented in this study can be found in either the Supplementary material or the online dataset 10.5281/zenodo.17651622.

## Notes

https://doi.org/10.5281/zenodo.17651622

### Competing Interest Statement

The authors have declared no competing interest.

## References

1. Zhang, W. & Nielsen, D. R. Synthetic Biology Applications in Industrial Microbiology. Frontiers in Microbiology 5, 451 (Aug. 26, 2014).

2. Clarke, L. & Kitney, R. Developing Synthetic Biology for Industrial Biotechnology Applications. Biochemical Society Transactions 48, 113–122 (Feb. 28, 2020).

3. Weber, E., Engler, C., Gruetzner, R., Werner, S. & Marillonnet, S. A Modular Cloning System for Standardized Assembly of Multigene Constructs. PLOS ONE 6, e16765 (Feb. 18, 2011).

4. Engler, C. et al. A Golden Gate Modular Cloning Toolbox for Plants. ACS Synthetic Biology 3, 839–843 (Nov. 21, 2014).

5. Lee, M. E., DeLoache, W. C., Cervantes, B. & Dueber, J. E. A Highly Characterized Yeast Toolkit for Modular, Multipart Assembly. ACS Synthetic Biology 4, 975–986 (Sept. 18, 2015).

6. Rajkumar, A. S., Varela, J. A., Juergens, H., Daran, J. M. G. & Morrissey, J. P. Biological Parts for Kluyveromyces Marxianus Synthetic Biology. Frontiers in Bioengineering and Biotechnology 7, 97 (MAY Jan. 1, 2019).

7. Stukenberg, D. et al. The Marburg Collection: A Golden Gate DNA Assembly Framework for Synthetic Biology Applications in Vibrio Natriegens. ACS Synthetic Biology 10, 1904– 1919 (Aug. 20, 2021).

8. Deal, C., De Wannemaeker, L. & De Mey, M. Towards a Rational Approach to Promoter Engineering: Understanding the Complexity of Transcription Initiation in Prokaryotes. FEMS Microbiology Reviews 48, fuae004 (Feb. 21, 2024).

9. Jin, L.-Q. et al. Promoter Engineering Strategies for the Overproduction of Valuable Metabolites in Microbes. Applied Microbiology and Biotechnology 103, 8725–8736 (Nov. 1, 2019).

10. Xu, N., Wei, L. & Liu, J. Recent Advances in the Applications of Promoter Engineering for the Optimization of Metabolite Biosynthesis. World Journal of Microbiology and Biotechnology 35, 33 (Jan. 31, 2019).

11. Van Brempt, M. et al. Biosensor-Driven, Model-Based Optimization of the Orthogonally Expressed Naringenin Biosynthesis Pathway. Microbial Cell Factories 21, 49 (Mar. 27, 2022).

12. Khamwachirapithak, P. et al. Optimizing Ethanol Production in Saccharomyces Cerevisiae at Ambient and Elevated Temperatures through Machine Learning-Guided Combinatorial Promoter Modifications. ACS Synthetic Biology 12, 2897–2908 (Sept. 8, 2023).

13. Cautereels, C. et al. Combinatorial Optimization of Gene Expression through Recombinase-Mediated Promoter and Terminator Shuffing in Yeast. Nature Communications 15, 1112 (Feb. 7, 2024).

14. De Mey, M., Maertens, J., Lequeux, G. J., Soetaert, W. K. & Vandamme, E. J. Construction and Model-Based Analysis of a Promoter Library for E. Coli: An Indispensable Tool for Metabolic Engineering. BMC biotechnology 7, 34 (June. 18, 2007).

15. Decoene, T., Maeseneire, S. L. D. & Mey, M. D. Modulating Transcription through Development of Semi-Synthetic Yeast Core Promoters. PLOS ONE 14, e0224476 (Nov. 5, 2019).

16. De Wannemaeker, L., Bervoets, I. & De Mey, M. Unlocking the Bacterial Domain for Industrial Biotechnology Applications Using Universal Parts and Tools. Biotechnology Advances 60, 108028 (Nov. 1, 2022).

17. Quehenberger, J., Shen, L., Albers, S.-V., Siebers, B. & Spadiut, O. Sulfolobus – A Potential Key Organism in Future Biotechnology. Frontiers in Microbiology 8, 2474 (Dec. 12, 2017).

18. Yin, J., Chen, J.-C., Wu, Q. & Chen, G.-Q. Halophiles, Coming Stars for Industrial Biotechnology. Biotechnology Advances 33, 1433–1442 (Nov. 15, 2015).

19. Schocke, L., Bräsen, C. & Siebers, B. Thermoacidophilic Sulfolobus Species as Source for Extremozymes and as Novel Archaeal Platform Organisms. Current Opinion in Biotechnology 59, 71–77 (Oct. 1, 2019).

20. Wagner, M. et al. Versatile Genetic Tool Box for the Crenarchaeote Sulfolobus Acidocaldarius. Frontiers in Microbiology 3, 214 (June. 13, 2012).

21. Berkner, S., Wlodkowski, A.Albers, S.-V. & Lipps, G. Inducible and Constitutive Promoters for Genetic Systems in Sulfolobus Acidocaldarius. Extremophiles 14, 249–259 (May. 1, 2010).

22. Van der Kolk, N. et al. Identification of XylR, the Activator of Arabinose/Xylose Inducible Regulon in Sulfolobus Acidocaldarius and Its Application for Homologous Protein Expression. Frontiers in Microbiology 11 (May. 26, 2020).

23. Liu, H. et al. BarR, an Lrp-type Transcription Factor in Sulfolobus Acidocaldarius, Regulates an Aminotransferase Gene in a β-Alanine Responsive Manner. Molecular Microbiology 92, 625–639 (2014).

24. Galdzicki, M. et al. The Synthetic Biology Open Language (SBOL) Provides a Community Standard for Communicating Designs in Synthetic Biology. Nature Biotechnology 32, 545–550 (June 2014).

25. Jensen, P. R. & Hammer, K. The Sequence of Spacers between the Consensus Sequences Modulates the Strength of Prokaryotic Promoters. Applied and Environmental Microbiology 64, 82–87 (Jan. 1998).

26. Han, L. et al. Development of a Novel Strategy for Robust Synthetic Bacterial Promoters Based on a Stepwise Evolution Targeting the Spacer Region of the Core Promoter in Bacillus Subtilis. Microbial Cell Factories 18, 96 (May. 29, 2019).

27. Recalde, A. et al. The Use of Thermostable Fluorescent Proteins for Live Imaging in Sulfolobus Acidocaldarius. Frontiers in Microbiology 15 (Sept. 9, 2024).

28. Liu, H. Characterization of a Transcription Factor Involved in the Regulation of !-Alanine Metabolism in the Hyperthermoacidophilic Archaeon Sulfolobus Acidocaldarius (Vrije Universiteit Brussel, 2015). BioRXiv Preprint - 11

29. Liu, H., Wang, K., Lindås, A.-C. & Peeters, E. The Genome-Scale DNA-binding Profile of BarR, a β-Alanine Responsive Transcription Factor in the Archaeon Sulfolobus Acidocaldarius. BMC Genomics 17, 569 (Aug. 8, 2016).

30. Ao, X. et al. The Sulfolobus Initiator Element Is an Important Contributor to Promoter Strength. Journal of Bacteriology 195, 5216– 5222 (Oct. 24, 2013).

31. Solovyev, V. & Salamov, A. Automatic Annotation of Microbial Genomes and Metagenomic Sequences. Nova Science Publishers Inc Metagenomics and its applications in agriculture, biomedicine and environmental studies. p 61–78 (In Li RW. (ed) 2011).

32. Davis, J. H., Rubin, A. J. & Sauer, R. T. Design, Construction and Characterization of a Set of Insulated Bacterial Promoters. Nucleic Acids Research 39, 1131 (Sept. 15, 2010).

33. Crooks, G. E., Hon, G.Chandonia, J.-M. & Brenner, S. E. WebLogo: A Sequence Logo Generator. Genome Research 14, 1188–1190 (June 2004).

34. Cohen, O. et al. Comparative Transcriptomics across the Prokaryotic Tree of Life. Nucleic Acids Research 44, W46 (Web Server issue May 6, 2016).

35. Peng, N., Xia, Q., Chen, Z., Liang, Y. X. & She, Q. An Upstream Activation Element Exerting Differential Transcriptional Activation on an Archaeal Promoter. Molecular Microbiology 74, 928–939 (2009).

36. Wagner, M. et al. Investigation of the malE Promoter and MalR, a Positive Regulator of the Maltose Regulon, for an Improved Expression System in Sulfolobus Acidocaldarius. Applied and Environmental Microbiology 80, 1072–1081 (Feb. 2014).

37. Hain, J., Reiter, W. D., Hüdepohl, U. & Zillig, W. Elements of an Archaeal Promoter Defined by Mutational Analysis. Nucleic Acids Research 20, 5423–5428 (Oct. 25, 1992).

38. Qureshi, S. A. & Jackson, S. P. Sequence-Specific DNA Binding by the S. Shibatae TFIIB Homolog, TFB, and Its Effect on Promoter Strength. Molecular Cell 1, 389–400 (Feb. 1, 1998).

39. Green, M. R. Molecular Cloning: A Laboratory Manual 4th ed. xxxiii+1890+46 (Cold Spring Harbor Laboratory Press, Cold Spring Harbor, N.Y, 2012).

40. Suzuki, S. & Kurosawa, N. Disruption of the Gene Encoding Restriction Endonuclease SuaI and Development of a Host-Vector System for the Thermoacidophilic Archaeon Sulfolobus Acidocaldarius. Extremophiles: life under extreme conditions 20, 139–148 (Mar. 1, 2016).

41. Brock, T. D., Brock, K. M., Belly, R. T. & Weiss, R. L. Sulfolobus: A New Genus of Sulfur-Oxidizing Bacteria Living at Low pH and High Temperature. Archiv fur Mikrobiologie 84, 54–68 (Mar. 1972).

42. Warren, D. J. Preparation of Highly Effcient Electrocompetent Escherichia Coli Using Glycerol/Mannitol Density Step Centrifugation. Analytical Biochemistry 413, 206–207 (June. 15, 2011).

43. Schleper, C., Kubo, K. & Zillig, W. The Particle SSV1 from the Extremely Thermophilic Archaeon Sulfolobus Is a Virus: Demonstration of Infectivity and of Transfection with Viral DNA. Proceedings of the National Academy of Sciences of the United States of America 89, 7645 (Aug. 8, 1992).

44. Quan, J. & Tian, J. Circular Polymerase Extension Cloning of Complex Gene Libraries and Pathways. PLOS ONE 4, e6441 (July. 30, 2009).

45. Sievers, F. et al. Fast, Scalable Generation of High-Quality Protein Multiple Sequence Alignments Using Clustal Omega. Molecular Systems Biology 7, 539 (Oct. 11, 2011).

46. Okonechnikov, K., Golosova, O., Fursov, M. & the UGENE team. Unipro UGENE: A Unified Bioinformatics Toolkit. Bioinformatics 28, 1166–1167 (Apr. 15, 2012).

