## Supplementary Materials for "Development and characterisation of a promoter library for *Sulfolobus acidocaldarius*"

| Plasmid | Organism | Promoter | Reporter | Backbone | Selection marker | Source |
|---|---|---|---|---|---|---|
| pBR322-ProB | *E. coli* | ProB | mKate2 | pBR322 | $Kan^{R}$ | In-house |
| pBR322-$P_{Saci\_2137}$ | *E. coli* | $P_{Saci\_2137}$ | mKate2 | pBR322 | $Kan^{R}$ | This study |
| pBR322-$P_{Saci\_2137core}$ | *E. coli* | $P_{Saci\_2137core}$ | mKate2 | pBR322 | $Kan^{R}$ | This study |
| pBR322-$P_{EcLib}$ | *E. coli* | $P_{EcLib}$ | mKate2 | pBR322 | $Kan^{R}$ | This study |
| pBR322-$P_{Ec1}$ | *E. coli* | $P_{Ec1}$ | mKate2 | pBR322 | $Kan^{R}$ | This study |
| pBR322-$P_{Ec2}$ | *E. coli* | $P_{Ec2}$ | mKate2 | pBR322 | $Kan^{R}$ | This study |
| pBR322-$P_{Ec3}$ | *E. coli* | $P_{Ec3}$ | mKate2 | pBR322 | $Kan^{R}$ | This study |
| pBR322-$P_{Ec4}$ | *E. coli* | $P_{Ec4}$ | mKate2 | pBR322 | $Kan^{R}$ | This study |
| pBR322-$P_{Ec5}$ | *E. coli* | $P_{Ec5}$ | mKate2 | pBR322 | $Kan^{R}$ | This study |
| pBR322-$P_{Ec6}$ | *E. coli* | $P_{Ec6}$ | mKate2 | pBR322 | $Kan^{R}$ | This study |
| pBR322-$P_{Ec7}$ | *E. coli* | $P_{Ec7}$ | mKate2 | pBR322 | $Kan^{R}$ | This study |
| pBR322-$P_{Ec8}$ | *E. coli* | $P_{Ec8}$ | mKate2 | pBR322 | $Kan^{R}$ | This study |
| pBR322-$P_{Ec9}$ | *E. coli* | $P_{Ec9}$ | mKate2 | pBR322 | $Kan^{R}$ | This study |
| pBR322-$P_{Ec10}$ | *E. coli* | $P_{Ec10}$ | mKate2 | pBR322 | $Kan^{R}$ | This study |
| pSVA1450 | *S. acidocaldarius* | $P_{malE\_pSVA1450}$ | LacS | pRN1 | Ura | [36] |
| pJL1601 | *S. acidocaldarius* | $P_{malE\_pJL1601}$ | LacS | pRN1 | Ura | MICR |
| pRN1-$P_{Saci\_2137}$ | *S. acidocaldarius* | $P_{Saci\_2137}$ | LacS | pRN1 | Ura | This study |
| pRN1-$P_{Saci\_2137core}$ | *S. acidocaldarius* | $P_{Saci\_2137core}$ | LacS | pRN1 | Ura | This study |
| pRN1-$P_{sac7d}$ | *S. acidocaldarius* | $P_{sac7d}$ | LacS | pRN1 | Ura | This study |
| pRN1-$P_{Ec10\_Sa1}$ | *S. acidocaldarius* | $P_{Ec10\_Sa1}$ | LacS | pRN1 | Ura | This study |
| pRN1-$P_{Ec6\_Sa2}$ | *S. acidocaldarius* | $P_{Ec6\_Sa2}$ | LacS | pRN1 | Ura | This study |
| pRN1-$P_{Ec7\_Sa3}$ | *S. acidocaldarius* | $P_{Ec7\_Sa3}$ | LacS | pRN1 | Ura | This study |
| pRN1-$P_{Ec8\_Sa4}$ | *S. acidocaldarius* | $P_{Ec8\_Sa4}$ | LacS | pRN1 | Ura | This study |
| pRN1-$P_{Ec4\_Sa5}$ | *S. acidocaldarius* | $P_{Ec4\_Sa5}$ | LacS | pRN1 | Ura | This study |
| pRN1-$P_{Ec9\_Sa6}$ | *S. acidocaldarius* | $P_{Ec9\_Sa6}$ | LacS | pRN1 | Ura | This study |
| pRN1-$P_{Ec3\_Sa7}$ | *S. acidocaldarius* | $P_{Ec3\_Sa7}$ | LacS | pRN1 | Ura | This study |

**Supplementary Table 1:** Plasmid constructs used in this chapter and their properties. *E. coli*: *Escherichia coli*, *S. acidocaldarius*: *Sulfolobus acidocaldarius*, $Kan^{R}$: kanamycin, Ura: uracil, MICR: MICR research group (Vrije Universiteit Brussel, Brussels, Belgium.)

**Supplementary Table 2:** Plasmid sequence map of pBR322-ProB. CDS: coding sequence, sRNA: small RNA, Ori: origin of replication.

| **Plasmid** | **pBR322-ProB** | **6403 bp DNA circular** |
|---|---|---|
| **Features** | **Qualifiers** | **Location** |
| Promoter | ProB | 0..144 |
| CDS | mKate2 | 171..870 |
| Terminator | Biofab terminator_FAB391 | 902..984 |
| CDS | rop | complement(1208..1400) |
| CDS | $Kan^R$ | complement(1521..2313) |
| Terminator | TT5-T7term | complement(2549..2637) |
| CDS | sfGFP | complement(2658..3375) |
| Promoter | promotor 14 | complement(3414..3465) |
| Terminator | TT4-trp_att | complement(3750..3837) |
| sRNA | cpxQ | complement(3857..3915) |
| Promoter | pBAD promoter | complement(3857..3966) |
| CDS | araC | 4225..5104 |
| Ori | pMB1 | 5611..6200 |

```
   1 CACAGCTAAC ACCACGTCGT CCCTATCTGC TGCCCTAGGT CTATGAGTGG TTGCTGGATA ACTTTACGGG
  71 CATGCATAAG GCTCGTAATA TATATTCAGG GAGACCACAA CGGTTTCCCT CTACAAATAA TTTTGTTTAA
 141 CTTTTACTAG AGTCACACAG GAAAGTACTA GATGGTTAGC GAGCTGATCA AAGAAAACAT GCACATGAAA
 211 CTGTATATGG AAGGCACCGT GAATAACCAC CACTTTAAAT GTACCAGCGA AGGTGAAGGT AAACCGTATG
 281 AAGGCACCCA GACCATGCGT ATTAAAGCAG TTGAAGGTGG TCCGCTGCCG TTTGCATTTG ATATTCTGGC
 351 AACCAGCTTT ATGTATGGCA GCAAAACCTT TATTAACCAT ACCCAGGGTA TCCCGGATTT TTTCAAACAG
 421 AGCTTTCCGG AAGGTTTTAC CTGGGAACGT GTTACCACCT ATGAAGATGG TGGTGTTCTG ACCGCAACCC
 491 AGGATACCAG TCTGCAGGAT GGTTGTCTGA TTTATAATGT GAAAATTCGC GGTGTGAACT TTCCGAGCAA
 561 TGGTCCGGTT ATGCAGAAAA AAACCCTGGG TTGGGAAGCA AGCACCGAAA CCCTGTATCC GGCAGATGGT
 631 GGTCTGGAAG GTCGTGCAGA TATGGCACTG AAACTGGTTG GTGGTGGTCA TCTGATTTGC AATCTGAAAA
 701 CCACCTATCG TAGCAAAAAA CCGGCAAAAA ATCTGAAAAT GCCTGGCGTG TATTATGTTG ATCGTCGTCT
 771 GGAACGTATT AAAGAGGCAG ATAAAGAAAC CTATGTGGAA CAGCATGAAG TTGCAGTTGC ACGTTATTGT
 841 GATCTGCCGA GCAAACTGGG TCACCGCTGA TAACCATGGG CTAGCGGTTT GAAGGGTATT GGTCGGTCAG
 911 TTTCACCTGA TTTACGTAAA AACCCGCTTC GGCGGGTTTT TGCTTTTGGA GGGGCAGAAA GATGAATGAC
 981 TGTCCCTTAC GCATCTGTGC GGTATTTCAC ACCGCATATG GTGCACTCTC AGTACAATCT GCTCTGATGC
1051 CGCATAGTTA AGCCAGTATA CACTCCGCTA TCGCTACGTG ACTGGGTCAT GGCTGCGCCC CGACACCCGC
1121 CAACACCCGC TGACGCGCCC TGACGGGCTT GTCTGCTCCC GGCATCCGCT TACAGACAAG CTGTGACCGT
1191 CTCCGGGAGC TGCATGTGTC AGAGGTTTTC ACCGTCATCA CCGAAACGCG CGAGGCAGCT GCGGTAAAGC
1261 TCATCAGCGT GGTCGTGAAG CGATTCACAG ATGTCTGCCT GTTCATCCGC GTCCAGCTCG TTGAGTTTCT
1331 CCAGAAGCGT TAATGTCTGG CTTCTGATAA AGCGGGCCAT GTTAAGGGCG GTTTTTTCCT GTTTGGTCAC
1401 TGATGCCTCC GTGTAAGGGG GATTTCTGTT CATGGGGGGA AGGCGGCGGT GGAATCGAAA TCTCGTGATG
1471 GCAGGTTGGG CGTCGCTTGG TCGGTCATTT CGAACCCCAG AGTCCCGCTC AGAAGAACTC GTCAAGAAGG
1541 CGATAGAAGG CGATGCGCTG CGAATCGGGA GCGGCGATAC CGTAAAGCAC GAGGAAGCGG TCAGCCCATT
1611 CGCCGCCAAG CTCTTCAGCA ATATCACGGG TAGCCAACGC TATGTCCTGA TAGCGGTCCG CCACACCCAG
1681 CCGGCCACAG TCGATGAATC CAGAAAAGCG GCCATTTTCC ACCATGATAT TCGGCAAGCA GGCATCGCCA
1751 TGGGTCACGA CGAGATCCTC GCCGTCGGGC ATGCGCGCCT TGAGCCTGGC GAACAGTTCG GCTGGCGCGA
1821 GCCCCTGATG CTCTTCGTCC AGATCATCCT GATCGACAAG ACCGGCTTCC ATCCGAGTAC GTGCTCGCTC
1891 GATGCGATGT TTCGCTTGGT GGTCGAATGG GCAGGTAGCC GGATCAAGCG TATGCAGCCG CCGCATTGCA
1961 TCAGCCATGA TGGATACTTT CTCGGCAGGA GCAAGGTGAG ATGACAGGAG ATCCTGCCCC GGCACTTCGC
2031 CCAATAGCAG CCAGTCCCTT CCCGCTTCAG TGACAACGTC GAGCACAGCT GCGCAAGGAA CGCCCGTCGT
2101 GGCCAGCCAC GATAGCCGCG CTGCCTCGTC CTGCAGTTCA TTCAGGGCAC CGGACAGGTC GGTCTTGACA
2171 AAAAGAACCG GGCGCCCCTG CGCTGACAGC CGGAACACGG CGGCATCAGA GCAGCCGATT GTCTGTTGTG
2241 CCCAGTCATA GCCGAATAGC CTCTCCACCC AAGCGGCCGG AGAACCTGCG TGCAATCCAT CTTGTTCAAT
2311 CATGCGAAAC GATCCTCATC CTGTCTCTTG ATCAGATCTT GATCCCCTGC GCCATCAGAT CCTTGGCGGC
2381 AAGAAAGCCA TCCAGTTTAC TTTGCAGGGC TTCCCAACCT TACCAGAGGG CGCCCCAGCT GGCAATTCCG
2451 GTTCGCTTGC TGTCCATAAA ACCGCCCAGT CTAGCTATCG CCATGTAAGC CCACATAGCT AGCATCCTAG
```

```
2521 GACTTGGAGC CGTGCGATAA TATCATGCAC TTCTTAAACA TAAAGTGTCT CCTTATAAAC GCAGAAAGGC
2591 CCACCCGAAG GTGAGCCAGT GTGATTACAT TTTCTCTTGA GGGTTGTGGT TAAGGCCGTT TGGCATTGTT
2661 ACTTATAGAG TTCATCCATG CCATGAGTAA TCCCCGCTGC CGTGACGAAT TCCAACAGGA CCATATGGTC
2731 ACGCTTCTCG TTTGGGTCTT TACTAAGAAC GCTTTGTGTA GACAGGTAAT GATTATCCGG GAGCAGAACG
2801 GGGCCATCGC CAATCGGAGT ATTCTGCTGG TAATGATCAG CCAGCTGCAC GGAACCGTCC TCTACGTTAT
2871 GACGGATTTT AAAGTTGGCT TTGATGCCAT TTTTCTGTTT ATCCGCTGTA ATGTATACGT TGTGCGAATT
2941 AAAGTTGTAT TCCAATTTGT GCCCCAGGAT ATTCCCATCC TCTTTGAAAT CGATACCTTT TAATTCAATG
3011 CGGTTAACTA AGGTATCGCC TTCAAATTTC ACTTCCGCGC GGGTCTTATA CGTCCCATCG TCTTTGAAGC
3081 TAATAGTCCG TTCCTGCACA TAACCTTCAG GCATTGCGCT TTTAAAAAAG TCGTGGCGTT TCATGTGATC
3151 TGGATAGCGT GAAAAGCATT GGACGCCATA CGTCAACGTT GTGACCAGCG TCGGCCAAGG CACGGGCAGC
3221 TTACCGGTGG TACAGATGAA CTTCAGGGTC AGCTTACCAT TGGTGGCATC TCCCTCTCCT TCGCCACGAA
3291 CAGAGAACTT ATGACCGTTC ACGTCTCCGT CCAGTTCTAC TAAAATCGGC ACAACACCGG TAAAAAGCTC
3361 TTCGCCCTTG CCCATGAAAT TTTTCCTTGC CTAGAAAAAT TATTTCTAGA GGGGCCCTAT TATACCATAT
3431 GCCGGCCAAG ATGTCAAGAA ACTTATAGAA TGAAGCTAGC CCATGGTTCC GATGGCGTGC ATCAGATGGG
3501 TACAAGCACT GAAGCGATGT AAAGCGGGCG CACCCTCTGA GAATTAACTC ACGGTTAAAC TCTCTCTGCA
3571 ACTTTGTCAT TTAGTAAGTA TCTTAGCGAG CAGTTCTTCA ATAAAGCGCT GCGGCAATCT TATGGTACTC
3641 GACGCCTGAC TAATTATTTA ATGGTAGTAA TTGCAATTAA TGACGTCAGG TGGCACTTTT CGTCATATGG
3711 TACTCGACGC CTGACTAATT ATTTAATGGT AGTAATTGCA TGTTTGCATT GTTATTCTCT AATTTTGTTC
3781 AAAAAAAAGC CCGCTTATTA GGCGGGCTGG GTATCTGATT GCTTTACGCA TGGTGAAGCT TTGTCCCTTT
3851 GCTGGCCAAA AAAACCCCCA CAGCATGTGG GGGAAGACAG GGATGGTGTC TATGGCAAGG AAAAATGGAG
3921 AAACAGTAGA GAGTTGCGAT AAAAAGCGTC AGGTAGGATC CGCTAATCTT ATGGATAAAA ATGCTATGGC
3991 ATAGCAAAGT GTGACGCCGT GCAAATAATC AATGTGGACT TTTCTGCCGT GATTATAGAC ACTTTTGTTA
4061 CGCGTTTTTG TCATGGCTTT GGTCCCGCTT TGTTACAGAA TGCTTTTAAT AAGCGGGGTT ACCGGTTTGG
4131 TTAGCGAGAA GAGCCAGTAA AAGACGCAGT GACGGCAATG TCTGATGCAA TATGGACAAT TGGTTTCTTC
4201 TCTGAATGGC GGGAGTATGA AAAGTATGGC TGAAGCGCAA AATGATCCCC TGCTGCCGGG ATACTCGTTT
4271 AATGCCCATC TGGTGGCGGG TTTAACGCCG ATTGAGGCCA ACGGTTATCT CGATTTTTTT ATCGACCGAC
4341 CGCTGGGAAT GAAAGGTTAT ATTCTCAATC TCACCATTCG CGGTCAGGGG GTGGTGAAAA ATCAGGGACG
4411 AGAATTTGTT TGCCGACCGG GTGATATTTT GCTGTTCCCG CCAGGAGAGA TTCATCACTA CGGTCGTCAT
4481 CCGGAGGCTC GCGAATGGTA TCACCAGTGG GTTTACTTTC GTCCGCGCGC CTACTGGCAT GAATGGCTTA
4551 ACTGGCCGTC AATATTTGCC AATACGGGGT TCTTTCGCCC GGATGAAGCG CACCAGCCGC ATTTCAGCGA
4621 CCTGTTTGGG CAAATCATTA ACGCCGGGCA AGGGGAAGGG CGCTATTCGG AGCTGCTGGC GATAAATCTG
4691 CTTGAGCAAT TGTTACTGCG GCGCATGGAA GCGATTAACG AGTCGCTCCA TCCACCGATG GATAATCGGG
4761 TACGCGAGGC TTGTCAGTAC ATCAGCGATC ACCTGGCAGA CAGCAATTTT GATATCGCCA GCGTCGCACA
4831 GCATGTTTGC TTGTCGCCGT CGCGTCTGTC ACATCTTTTC CGCCAGCAGT TAGGGATTAG CGTCTTAAGC
4901 TGGCGCGAGG ACCAACGTAT CAGCCAGGCG AAGCTGCTTT TGAGCACCAC CCGGATGCCT ATCGCCACCG
4971 TCGGTCGCAA TGTTGGTTTT GACGATCAAC TCTATTTCTC GCGGGTATTT AAAAAATGCA CCGGGGCCAG
5041 CCCGAGCGAG TTCCGTGCCG GTTGTGAAGA AAAAGTGAAT GATGTAGCCG TCAAGTTGTC ATAATAAATC
5111 GATGCAGGTG GCACTTTTCG GGGAAATGTG CGCGGAACCC CGCTAGCCCA TGCTGAAGGC GCTCCTCATA
5181 CTGGCTGAAT ACATGACCCG ACACGTTATC AGAATTTATC ATGTCATGAT TGCTTACAAT ACTCACATAA
5251 TTGTGAAAAT TGTAGAAGCC GCAACATTCA ATTATCAATC AAAATCATGT CCGCGCCAGT TTCGTTCCTC
5321 GCCGTTGCAC TAATACAGGG TAATTCCCTC ACCACATAAG GAATTAATGA CGTCAGGTGG CACTTTTCCA
5391 CACAAGCTGA GGCGACGTCT ATTTCCAGTC GGGAAACCTG TCGTGCCAGC TCTGTCAGAC CAAGTTTACT
5461 CATATATACT TTAGATTGAT TTAAAACTTC ATTTTTAATT TAAAAGGATC TAGGTGAAGA TCCTTTTTGA
5531 TAATCTCATG ACCAAAATCC CTTAACGTGA GTTTTCGTTC CACTGAGCGT CAGACCCCGT AGAAAAGATC
5601 AAAGGATCTT CTTGAGATCC TTTTTTTCTG CGCGTAATCT GCTGCTTGCA AACAAAAAAA CCACCGCTAC
5671 CAGCGGTGGT TTGTTTGCCG GATCAAGAGC TACCAACTCT TTTTCCGAAG GTAACTGGCT TCAGCAGAGC
5741 GCAGATACCA AATACTGTCC TTCTAGTGTA GCCGTAGTTA GGCCACCACT TCAAGAACTC TGTAGCACCG
5811 CCTACATACC TCGCTCTGCT AATCCTGTTA CCAGTGGCTG CTGCCAGTGG CGATAAGTCG TGTCTTACCG
5881 GGTTGGACTC AAGACGATAG TTACCGGATA AGGCGCAGCG GTCGGGCTGA ACGGGGGGTT CGTGCACACA
5951 GCCCAGCTTG GAGCGAACGA CCTACACCGA ACTGAGATAC CTACAGCGTG AGCTATGAGA AAGCGCCACG
6021 CTTCCCGAAG GGAGAAAGGC GGACAGGTAT CCGGTAAGCG GCAGGGTCGG AACAGGAGAG CGCACGAGGG
6091 AGCTTCCAGG GGGAAACGCC TGGTATCTTT ATAGTCCTGT CGGGTTTCGC CACCTCTGAC TTGAGCGTCG
6161 ATTTTTGTGA TGCTCGTCAG GGGGGCGGAG CCTATGGAAA AACGCCAGCA ACGCGGCCTT TTTACGGTTC
6231 CTGGCCTTTT GCTGGCCTTT TGCTCACATG TTCTTTCCTG CGTTATCCCC TGATTCTGTG GATAACCGTA
6301 TTACCGCCTT TGAGTGAGCT GATACCGCTC GCCGCAGCCG AACGACCGAG CGCAGCGAGT CAGTGAGCGA
6371 GGAAGCGGAA GAGCGCCTGA TGCGGTATTT TCT
```

**Supplementary Table 3:** Plasmid sequence map of pBR322-P$_{\mathrm{Saci_2137}}$. The P$_{\mathrm{Saci_2137}}$ promoter sequence is highlighted in yellow and can be substituted by the promoter sequences in Supplementary Table 6 to result in the plasmid maps of pBR322-P$_{\mathrm{Saci_2137core}}$ and pBR322-P$_{\mathrm{Ec1\text{-}10}}$. CDS: coding sequence, Ori: origin of replication.

| **Plasmid** | **pBR322-P$_{\mathrm{Saci_2137}}$** | **3766 bp DNA circular** |
|---|---|---|
| **Features** | **Qualifiers** | **Location** |
| Promoter | P$_{\mathrm{Saci_2137}}$ | 0..173 |
| CDS | mKate2 | 173..872 |
| Terminator | BioFab terminator FAB391 | 904..986 |
| CDS | rop | complement(1210..1402) |
| CDS | Kan$^{R}$ | complement(1523..2315) |
| Ori | pMB1 | 2974..3563 |

```
   1 CATTTGTATA TAATACATTT TTTCCAATAT ATAAGCATAG ATATAAAATT TTGTATAAAT CTCTATCATC
  71 TCTTTTAAAA TTGTACGCAT AGTATAGTAA GGGAATGTTA GTTTGTATAT AATACAAATC ACTCTCAAAG
 141 AGGTTTATAA TCTGTTTAAG AAGAATATTA CTTATGGTTA GCGAGCTGAT CAAAGAAAAC ATGCACATGA
 211 AACTGTATAT GGAAGGCACC GTGAATAACC ACCACTTTAA ATGTACCAGC GAAGGTGAAG GTAAACCGTA
 281 TGAAGGCACC CAGACCATGC GTATTAAAGC AGTTGAAGGT GGTCCGCTGC CGTTTGCATT TGATATTCTG
 351 GCAACCAGCT TTATGTATGG CAGCAAAACC TTTATTAACC ATACCCAGGG TATCCCGGAT TTTTTCAAAC
 421 AGAGCTTTCC GGAAGGTTTT ACCTGGGAAC GTGTTACCAC CTATGAAGAT GGTGGTGTTC TGACCGCAAC
 491 CCAGGATACC AGTCTGCAGG ATGGTTGTCT GATTTATAAT GTGAAAATTC GCGGTGTGAA CTTTCCGAGC
 561 AATGGTCCGG TTATGCAGAA AAAAACCCTG GGTTGGGAAG CAAGCACCGA AACCCTGTAT CCGGCAGATG
 631 GTGGTCTGGA AGGTCGTGCA GATATGGCAC TGAAACTGGT TGGTGGTGGT CATCTGATTT GCAATCTGAA
 701 AACCACCTAT CGTAGCAAAA AACCGGCAAA AAATCTGAAA ATGCCTGGCG TGTATTATGT TGATCGTCGT
 771 CTGGAACGTA TTAAAGAGGC AGATAAAGAA ACCTATGTGG AACAGCATGA AGTTGCAGTT GCACGTTATT
 841 GTGATCTGCC GAGCAAACTG GGTCACCGCT GATAACCATG GGCTAGCGGT TTGAAGGGTA TTGGTCGGTC
 911 AGTTTCACCT GATTTACGTA AAAACCCGCT TCGGCGGGTT TTTGCTTTTG GAGGGGCAGA AAGATGAATG
 981 ACTGTCCCTT ACGCATCTGT GCGGTATTTC ACACCGCATA TGGTGCACTC TCAGTACAAT CTGCTCTGAT
1051 GCCGCATAGT TAAGCCAGTA TACACTCCGC TATCGCTACG TGACTGGGTC ATGGCTGCGC CCCGACACCC
1121 GCCAACACCC GCTGACGCGC CCTGACGGGC TTGTCTGCTC CCGGCATCCG CTTACAGACA AGCTGTGACC
1191 GTCTCCGGGA GCTGCATGTG TCAGAGGTTT TCACCGTCAT CACCGAAACG CGCGAGGCAG CTGCGGTAAA
1261 GCTCATCAGC GTGGTCGTGA AGCGATTCAC AGATGTCTGC CTGTTCATCC GCGTCCAGCT CGTTGAGTTT
1331 CTCCAGAAGC GTTAATGTCT GGCTTCTGAT AAAGCGGGCC ATGTTAAGGG CGGTTTTTTC CTGTTTGGTC
1401 ACTGATGCCT CCGTGTAAGG GGGATTTCTG TTCATGGGGG GAAGGCGGCG GTGGAATCGA AATCTCGTGA
1471 TGGCAGGTTG GGCGTCGCTT GGTCGGTCAT TTCGAACCCC AGAGTCCCGC TCAGAAGAAC TCGTCAAGAA
1541 GGCGATAGAA GGCGATGCGC TGCGAATCGG GAGCGGCGAT ACCGTAAAGC ACGAGGAAGC GGTCAGCCCA
1611 TTCGCCGCCA AGCTCTTCAG CAATATCACG GGTAGCCAAC GCTATGTCCT GATAGCGGTC CGCCACACCC
1681 AGCCGGCCAC AGTCGATGAA TCCAGAAAAG CGGCCATTTT CCACCATGAT ATTCGGCAAG CAGGCATCGC
1751 CATGGGTCAC GACGAGATCC TCGCCGTCGG GCATGCGCGC CTTGAGCCTG GCGAACAGTT CGGCTGGCGC
1821 GAGCCCCTGA TGCTCTTCGT CCAGATCATC CTGATCGACA AGACCGGCTT CCATCCGAGT ACGTGCTCGC
1891 TCGATGCGAT GTTTCGCTTG GTGGTCGAAT GGGCAGGTAG CCGGATCAAG CGTATGCAGC CGCCGCATTG
1961 CATCAGCCAT GATGGATACT TTCTCGGCAG GAGCAAGGTG AGATGACAGG AGATCCTGCC CCGGCACTTC
2031 GCCCAATAGC AGCCAGTCCC TTCCCGCTTC AGTGACAACG TCGAGCACAG CTGCGCAAGG AACGCCCGTC
2101 GTGGCCAGCC ACGATAGCCG CGCTGCCTCG TCCTGCAGTT CATTCAGGGC ACCGGACAGG TCGGTCTTGA
2171 CAAAAAGAAC CGGGCGCCCC TGCGCTGACA GCCGGAACAC GGCGGCATCA GAGCAGCCGA TTGTCTGTTG
2241 TGCCCAGTCA TAGCCGAATA GCCTCTCCAC CCAAGCGGCC GGAGAACCTG CGTGCAATCC ATCTTGTTCA
2311 ATCATGCGAA ACGATCCTCA TCCTGTCTCT TGATCAGATC TTGATCCCCT GCGCCATCAG ATCCTTGGCG
2381 GCAAGAAAGC CATCCAGTTT ACTTTGCAGG GCTTCCCAAC CTTACCAGAG GGCGCCCCAG CTGGCAATTC
2451 CGGTTCGCTT GCTGTCCATA AAACCGCCCA GTCTAGCTAT CGCCATGTAA GCCCACATAG CTAGCATCCT
2521 AGGACTTGGA GCCGTGCGAT AAACTGGCTG AATACATGAC CCGACACGTT ATCAGAATTT ATCATGTCAT
2591 GATTGCTTAC AATACTCACA TAATTGTGAA AATTGTAGAA GCCGCAACAT TCAATTATCA ATCAAAATCA
2661 TGTCCGCGCC AGTTTCGTTC CTCGCCGTTG CACTAATACA GGGTAATTCC CTCACCACAT AAGGAATTAA
2731 TGACGTCAGG TGGCACTTTT CCACACAAGC TGAGGCGACG TCTATTTCCA GTCGGGAAAC CTGTCGTGCC
2801 AGCTCTGTCA GACCAAGTTT ACTCATATAT ACTTTAGATT GATTTAAAAC TTCATTTTTA ATTTAAAAGG
2871 ATCTAGGTGA AGATCCTTTT TGATAATCTC ATGACCAAAA TCCCTTAACG TGAGTTTTCG TTCCACTGAG
2941 CGTCAGACCC CGTAGAAAAG ATCAAAGGAT CTTCTTGAGA TCCTTTTTTT CTGCGCGTAA TCTGCTGCTT
```

```
3011 GCAAACAAAA AAACCACCGC TACCAGCGGT GGTTTGTTTG CCGGATCAAG AGCTACCAAC TCTTTTTCCG
3081 AAGGTAACTG GCTTCAGCAG AGCGCAGATA CCAAATACTG TCCTTCTAGT GTAGCCGTAG TTAGGCCACC
3151 ACTTCAAGAA CTCTGTAGCA CCGCCTACAT ACCTCGCTCT GCTAATCCTG TTACCAGTGG CTGCTGCCAG
3221 TGGCGATAAG TCGTGTCTTA CCGGGTTGGA CTCAAGACGA TAGTTACCGG ATAAGGCGCA GCGGTCGGGC
3291 TGAACGGGGG GTTCGTGCAC ACAGCCCAGC TTGGAGCGAA CGACCTACAC CGAACTGAGA TACCTACAGC
3361 GTGAGCTATG AGAAAGCGCC ACGCTTCCCG AAGGGAGAAA GGCGGACAGG TATCCGGTAA GCGGCAGGGT
3431 CGGAACAGGA GAGCGCACGA GGGAGCTTCC AGGGGGAAAC GCCTGGTATC TTTATAGTCC TGTCGGGTTT
3501 CGCCACCTCT GACTTGAGCG TCGATTTTTG TGATGCTCGT CAGGGGGGCG GAGCCTATGG AAAAACGCCA
3571 GCAACGCGGC CTTTTTACGG TTCCTGGCCT TTTGCTGGCC TTTTGCTCAC ATGTTCTTTC CTGCGTTATC
3641 CCCTGATTCT GTGGATAACC GTATTACCGC CTTTGAGTGA GCTGATACCG CTCGCCGCAG CCGAACGACC
3711 GAGCGCAGCG AGTCAGTGAG CGAGGAAGCG GAAGAGCGCC TGATGCGGTA TTTTCT
```

**Supplementary Table 4:** Plasmid sequence map of pSVA1450 [36]. CDS: coding sequence, Orf: open reading frame, Ori: origin of replication.

| **Plasmid** | **pSVA1450** | **12014 bp DNA circular** |
|---|---|---|
| **Features** | **Qualifiers** | **Location** |
| Promoter | PmalE | 0..264 |
| CDS | LacS | 264..1734 |
| Ori | pRN1 | 1779..7138 |
| Orf | orf90b | 1779..1892 |
| Orf | orf56 | 2165..2336 |
| Orf | orf904 | 2316..5031 |
| Orf | orf80 | 5872..6115 |
| Orf | orf90a | 6328..6601 |
| Orf | orf72 | 6708..6927 |
| Orf | orf90b | 6969..7138 |
| CDS | PyrE | 7437..8025 |
| CDS | PyrF | 8011..8680 |
| CDS | MalR (Saci_1161) | complement(8805..9855) |
| Promoter | $P_{malR}$ | complement(9855..10046) |
| CDS | $Amp^R$ | 10209..11070 |
| Ori | pMB1 | 11240..11829 |

```
   1 CCAGATATCT GATAGTTGGA GAAATGCTGA TGATTGCACG CTTATCTTTT TTTACCTACT GCTTGTGGTT
  71 TAAAAATTTT ACTGGAAACT AGAAATAAAT TTAAGTAAAT TAAATAAAGG CTAATTAATA ATACTAATAT
 141 CATTTCGTAA ACTACATCTT ATAATACTTA AGTTGACATG TTCAACGGAG GTGTCCTTAA GTTTAGACCT
 211 TAAATATTTT ATATATATAT TAAGTTTATA AATAATTACG TGATTAAGTT AACCATGGAC TCATTTCCAA
 281 ATAGCTTTAG GTTTGGTTGG TCCCAGGCCG GATTTCAATC AGAAATGGGA ACACCAGGGT CAGAAGATCC
 351 AAATACTGAC TGGTATAAAT GGGTTCATGA TCCAGAAAAC ATGGCAGCGG GATTAGTAAG TGGAGATCTA
 421 CCAGAAAATG GGCCAGGCTA CTGGGGAAAC TATAAGACAT TTCACGATAA TGCACAAAAA ATGGGATTAA
 491 AAATAGCTAG ACTAAATGTG GAATGGTCTA GGATATTTCC TAATCCATTA CCAAGGCCAC AAAACTTTGA
 561 TGAATCAAAA CAAGATGTGA CAGAGGTTGA GATAAACGAA AACGAGTTAA AGAGACTTGA CGAGTACGCT
 631 AATAAAGACG CATTAAACCA TTACAGGGAA ATATTCAAGG ATCTTAAAAG TAGAGGACTT TACTTTATAC
 701 TAAACATGTA TCATTGGCCA TTACCTCTAT GGTTACACGA CCCAATAAGA GTAAGAAGAG GAGATTTTAC
 771 TGGACCAAGT GGTTGGCTAA GTACTAGAAC AGTTTACGAA TTCGCTAGAT TCTCAGCTTA TATAGCTTGG
 841 AAATTCGATG ATCTAGTGGA TGAGTACTCA ACAATGAATG AACCTAACGT TGTTGGAGGT TTAGGATACG
 911 TTGGTGTTAA GTCCGGTTTT CCCCCAGGAT ACCTAAGCTT TGAACTTTCC CGTAGGGCAA TGTATAACAT
 981 CATTCAAGCT CACGCAAGAG CGTATGATGG GATAAAGAGT GTTTCTAAAA AACCAGTTGG AATTATTTAC
1051 GCTAATAGCT CATTCCAGCC GTTAACGGAT AAAGATATGG AAGCGGTAGA GATGGCTGAA AATGATAATA
1121 GATGGTGGTT CTTTGATGCT ATAATAAGAG GTGAGATCAC CAGAGGAAAC GAGAAGATTG TAAGAGATGA
1191 CCTAAAGGGT AGATTGGATT GGATTGGAGT TAATTATTAC ACTAGGACTG TTGTGAAGAG GACTGAAAAG
1261 GGATACGTTA GCTTAGGAGG TTACGGTCAC GGATGTGAGA GGAATTCTGT AAGTTTAGCG GGATTACCAA
1331 CCAGCGACTT CGGCTGGGAG TTCTTCCCAG AAGGTTTATA TGACGTTTTG ACGAAATACT GGAATAGATA
1401 TCATCTCTAT ATGTACGTTA CTGAAAATGG TATTGCGGAT GATGCCGATT ATCAAAGGCC CTATTATTTA
1471 GTATCTCACG TTTATCAAGT TCATAGAGCA ATAAATAGTG GTGCAGATGT TAGAGGGTAT TTACATTGGT
1541 CTCTAGCTGA TAATTACGAA TGGGCTTCAG GATTCTCTAT GAGGTTTGGT CTGTTAAAGG TCGATTACAA
1611 CACTAAGAGA CTATACTGGA GACCCTCAGC ACTAGTATAT AGGGAAATCG CCACAAATGG CGCAATAACT
1681 GATGAAATAG AGCACTTAAA TAGCGTACCT CCAGTAAAGC CATTAAGGCA CTAAGCGGCC GCTCTAGAAC
1751 TAGTGGATCC AGATGTGTAT AAGAGACAGA CTTTATCACG CCCGCCTTTG CTTCTGTTTC TTACTCTCAA
1821 CTGTTTCTAA CTCTGCTAAC TCACAATCCC GCTAATCACA CTTGTGACTT TACTAACTCT GCAGCTTTCT
1891 AACTAAGCCC GCCCTGTCTA ACTCTGCTAA CTATCCCCCG CCTTAATCAC ACCCGTGATT TTGTTTTTCT
1961 AACTCTGCAG CTTTGCTTCT TACTTACAGC TTTCTACCTA ACTCTTCTAA TCTCTCTACT TTCCTAACTC
2031 TCTGCCCCCC CCCCCCCCCC CGAACCCGAC GAAATAGAAA GTTAATTGCA GGTATTAGCT TTTCACACGG
2101 GTAATACTGA AAAATCTACC GTTGTCAATT TTAAGGATAA TTGCGGATAC AATTTTGATC CACAAATGGG
2171 TAGACCATAC AAACTATTAA ATGGAATAAA ATTAGGAGTC TATATTCCAC AGGAATGGCA TGACAGATTA
2241 ATGGAAATCG CTAAGGAAAA GAATCTTACG TTAAGTGATG TGTGCAGGCT GGCAATTAAA GAGTATTTAG
```

```
2311 ACAATCATGA TAAACAAAAG AAGTAAGGTA ATTCTTCATG GAAATGTGAA AAAAACAAGA AGAACGGGGG
2381 TGTACATGAT TAGTTTAGAT AATTCAGGTA ATAAAGATTT CTCTTCGAAT TTCTCAAGTG AGAGAATTAG
2451 ATATGCTAAG TGGTTTCTAG AACACGGCTT TAATATTATC CCAATCGACC CTGAGAGCAA AAAACCTGTA
2521 CTGAAGGAGT GGCAAAAGTA TAGTCATGAA ATGCCTTCTG ATGAGGAGAA ACAGAGATTT CTAAAGATGA
2591 TTGAAGAGGG GTATAATTAC GCTATTCCAG GGGGACAAAA AGGACTAGTG ATTCTCGATT TTGAGAGTAA
2661 GGAGAAGTTG AAAGCATGGA TAGGAGAATC AGCGTTAGAA GAGCTCTGTA GAAAAACATT ATGCACAAAC
2731 ACGGTTCATG GCGGAATCCA CATCTATGTT TTAAGTAACG ACATTCCGCC ACACAAAATC AATCCTCTCT
2801 TTGAAGAGAA TGGTAAGGGC ATAATTGATT TACAAAGTTA TAACAGTTAC GTTTTAGGAC TTGGCTCTTG
2871 TGTGAACCAT TTACACTGTA CTACTGATAA ATGCCCGTGG AAAGAACAGA ACTACACTAC TTGTTATACA
2941 TTATATAATG AATTAAAGGA AATTAGTAAA GTTGACTTAA AGAGTCTCCT AAGATTCCTG GCGGAAAAGG
3011 GCAAACGTTT AGGCATAACA TTAAGCAAAA CGGCAAAGGA GTGGTTAGAA GGAAAGAAGG AAGAAGAGGA
3081 CACGGTGGTG GAGTTTGAGG AGTTGAGAAA GGAATTAGTC AAACGTGACA GTGGGAAACC AGTTGAGAAA
3151 ATTAAAGAGG AGATATGTAC TAAGAGCCCG CCTAAACTGA TTAAGGAAAT AATATGTGAA AACAAGACTT
3221 ACGCTGATGT AAATATTGAT AGGAGTAGAG GGGATTGGCA TGTTATACTC TACTTAATGA AACACGGGGT
3291 TACGGATCCA GACAAGATAT TGGAATTATT ACCGCGTGAT TCAAAAGCAA AAGAAAATGA AAAGTGGAAT
3361 ACACAGAAAT ATTTTGTTAT AACTCTTTCG AAAGCCTGGA GTGTAGTAAA GAAGTACTTA GAAGCTAAAA
3431 GGAAAGCACA AAAAGACAAA AGCACGGCTA AGGCTCTGTT AATAGAGGCG ATTGCTGAAG AGGTGTTACA
3501 TGAACATTTT CTCGTTACCT TTATACAGAC TGATCAATTG AAGGAGTCAA AAATTGGGCT TTTCAGATTC
3571 AACAAGAAGA AGGGGATTTT CGAACCATTT GATGAAAGAA TTGAGAAGAT AATTAATGTA AAACTTGAGG
3641 AGTATAAGGA ATTCCCCCTG GGTTCTGATA AATCACGTGT GATACGTAAT ATTAAAGAAG AGATTATGAG
3711 GAGAACACAG AGACTGTTAT TAGAAGAGTC ACTTAGAATT GCGTTTCGTA ATGGGACACT GGAATGGGAT
3781 AGCAAAGGAG TAACATGGTA TGATGTTAAA GAAAGAACTC CTAAGGTGTA TTCATTTAAT TATGTGGATT
3851 GGAACTTAAA GATTGAGGAG ATTGAGAAGT TTAACATGAA GGAAATAACA GTTGAAGATA TAGAGAATTT
3921 AGCCCGGCGT GTTTGCCCTA GATCACTTGA GACTTTTAAG CAGTGGGTTG ATGATAAATG GGTGTTATTG
3991 TTTGAAGTCA TAGGCTATAC ATTCTATCCA AAGTACATCT TTAATAAGGC TATACTTCTC ACGGGGGCTG
4061 GGGCAAACGG GAAATCGACT TTCCTTAATT TGCTGTTAAA GATTCTAGGA CAGAAGAACG TTTCCGCAAT
4131 GCCTTTGAAG AGGATCATGG AGAGTGATAG ATTTGCTTCA ATTGAACTAT TCCATAAACT AGCAAATGTA
4201 AGCAGTGAGT TGTTTGCCTT TAAGATCACA AACACAGACC TGTTTAAAAA ACTCACGGGG GAAGACTACA
4271 TAGAGGGACA GAAGAAGTTC AGAGACCCCA TATACTTCAT TAATTACGCA AAACTATTCA ACGCCACTAA
4341 CGAATTGCCC GTGGTTTCTG ATCAAAGTTA TGGATTCTGG AGAAGATGGA TTGTGATAGA ATTTCCACAC
4411 CAATTCCCGC CGGATCCTAA CTTTTTCGAT AAGACATTCA CGGTGGAGGA GGTTGAAGGA GTAATCACGG
4481 TAGCTGTTAT CGCATTCGCC CGTGTACTTC AACAGAAGAA ATTCGACTTT GAGGATAGCA GTGCTAATGT
4551 CAAAGAATTA TGGGAAAGAA AAACAGACAG TGTTTACGCA TTCGTGAAAG AGTTACTAGA AACCGAAAGG
4621 GCTGAATATG ATCCGGCAAA CGGCGATTTA TTCATGCCCA CTGAGGACTT CTACCAGGCT TATTTAGAAT
4691 GGAGCGAAGA GAACGACACA AAAGCCGAAA GCAAAGCCGT GGTAACCCAA AGATTGCAAA GCAAATTTAG
4761 AATAACAAAA GACAAGAAGA AGATTAATGG AAAAAGAGTT TGGTGTTATG TGGGAATAAG ATTGAAGAAT
4831 AACAATATAA GTACGGGGGG CGGACAGGAC GGCGATTCTC CTAATTCTTT ATTAGAATTA TATAAGGAAT
4901 TCCAGGGCAA AGTGGTAAGT AGGAAAGACT TATATGATAT GCTAAAGCTA AGAGCGTTTG AGTTGTTGGA
4971 GTTGTGCGAG AAGAGAAACA TATGTCACTG GATTGATGAA GAGCACGTGC GGTTTGATTG AGTCCTTCAA
5041 GTTTTCAATT TTTTAAATTG AATTTTTCAT CTGTAATGAC CAATTTATGT CCATAGTGTC CAACTTTTTT
5111 TCTATTATTT TGATACACGG TGGGACAATA ATATAATAAA ATAATGCCTT TTTAGTTGGA CACTATGGAC
5181 ATAAGTAGTC ACACCCGTGA TAATATTTGT ATAGTAATGG CGTTTTTCAA ATATTCTTAC AACAAAAATT
5251 TGAATTTATC TAACGTCATT CTCTCTATAG AACCATTCTC TTTATAGAGC TATACTTTAT TCTTATTAAA
5321 AATTTAGAAA TGGGGAATTT CTAAAAACTG GAATTCCCCA TTTCCTCTAG ACTCTTCTTT TTCTTCTCTT
5391 CTCTCTCCTG TCTTGAGTCT AGAGTCCATA AACACTGCTG CATAACAGCA TGACAACCGT GATAAAGTCT
5461 GAGTTCAAGA TACAAGATAT TTATAAGGAG TAGGCATTTT TTTCTCTCTT CTTAAAATAT TTTTCCAATC
5531 GGTTATCGAT ATTCTGGCAC TAGTGCCAGA AATTTACTCA AAAGAAGTAC AAAAATTCTC GCCACTTGGC
5601 GAGAAATTTG CTCAAAGTAG TGAAACAATA TGAAAAAGAA AGAGAACAAT CACAGATATG ATTAAACTGC
5671 TGCATGCAGC CACGAAATCA CAACTATGAT TATGCAGTCC ATATGTTAAT CCTGGTCGGA TCCGCAAAAT
5741 TTTAGTTATA AGAGTTAGCT AACACGATAA GGCAAACAGT ATTAATAAAG CGTTAATCCT ACCTCCACCG
5811 TGTTATTTAG CTAACTTTTT GCACGCCAAA AGATATTTAA CAGTCTGTTA ATCCTACTTT ACATGGGATC
5881 CCATATGAGT GATCTGAAGG AAAAGCTAAC TCTAACTCAA CTAATCCTGA TTCGGCTATC AAAATCTTGT
5951 CAAACCCTGG AAGAGTTAGA ACGATATACA GGTGCAAACA GAAATGTACT TCTCGTTACC TTGACACGAC
6021 TCCATAAAAA AGGCATAATC TACAGGAAAT GGCGTAGGTT TGGCGGTAGG AAGTATAGAG AATATTGTTT
6091 GAAAAGCCGT GACGAACTCC TGTGAAACTC CCCAGTTTAC CGTTATTATT GATACATATC GATACATAAT
6161 GATACATATG CACACATAAT GATACAGAAT AAGCGTAGAA TAATAGTATA AGCATAGTAT AAGCACATGA
6231 TAAGAGTATA AAATAGGCGT AAAAGCGTAC GGCTTTCAGC TATATATCCC CAAACCTACT ATTTTTCCAC
6301 GCTCCCTTAA ACAAAATTCT AAGATAATTT GTTTTACTTT ACTGTATATT ATTCAATGCA TCTAAAATTT
```

```
 6371 TACTGTATAC GTCTATCGTT TTACCTGATG CGTATACCAT TTTCCCCCAT GCACCGCATT TCGTCTATAT
 6441 ACTCCCATTC TTCTATTTTT TGCACATTTC ACGCAAAAAA CTTCCCATAC CCAATCCCGC CGTGCTTTTT
 6511 TTCCTTTTAC TGCTTACTCT TTCACATAGC CCCCTCTGCT TCTCTAACTC TGCAGTTTTG CTCAGGCTTT
 6581 TGCTTAATCT CTCTCACATA GCTCTTCTTC TTTCTGCTTT TCTAACTCTA CTTTCTTTCT TTTCCTCTAC
 6651 TTAACTACTC TTTCACTTCT ACTTTTCTAA CTCTTCTTTT GCTCTTCTCT TCTTAGCTTT GCTAACTATG
 6721 TTTCCTTTCT ACTTTCATTA TCTCTTCTTA GCTTTTGCTC TCTTTCTTTA CTCTTTATCT ACAGCTTCAC
 6791 TCTTCTTCTT CTTTTTCCAA TCTCTGATTT TTCTCACGAT TTCTAGACAA ATTCCCGTAA CCCCCAATCC
 6861 CCCCGTGCTA ACCCAATCTT TGATTATCTT AGACAATCTT TCCCTATGTT TGACTTTCTT AGACTAACTT
 6931 AGACAGTTTC AAACAAACTA TTCCTACTTA TGCCGACTTA TGCCGACTTC TAACAACATC TTACGACATC
 7001 TTATGACATC TTACGACATC TTACAAATTT TAGGACTTTC ATTTCTATAC CCAAACTCTC TGTTTTTTTG
 7071 CACTTTCGTG CAAACAATTC TATATTCCAA TTCCAGACTC CTATACCCCC CATGCTTCTA CTTTATCACT
 7141 GTCTCTTATA CACATCTTCG AGGGGGGGCC GCTCTAGAAC TAGTGGATCC CCCGGGCTGC AGTATGAAAG
 7211 CTTTGAGCAG TTCTAGTACT TGGCTCAAAG AATGCTAATG AAACTACTTT CCCTGATAGA TAATTTAGTT
 7281 TTTTTACATC ATAGAACTTG TCTGCAAGTT CGAATATTCT AAAGTAGTCA TCTCTGGTCA AGTCAAGCGA
 7351 CGAAACTACG TGTCTTAATC TCACAAAGCC CTTATAACTG GTTATGATAG AAGTATATTT AAATTCTTTT
 7421 TCACAGAGTC TCTACGTATG AATTTCGCAG AAGTCTTACT CGAAAGGAAA TTATTATTAA TAGGAAGTTT
 7491 CGTTTTAACA TCAGGTAAGG TTAGTCCATA TTACTTAGAC TTAAGACCTT TACCAAATTA TCCAGAATTT
 7561 TACGATATAG TTAATCAAGC TATAAAGAAA GCAAAAGATA TACCCCATGA TATAATAGTA GGAATAGCCA
 7631 CTGGAGGAGT TCCCTTATCG GCATTCATAG CTTGTAACCT TAAAGAGCCT ATGGGATATA TTAGAATAGA
 7701 AAAGAAAGGT CATGGAACTA ATCGTACATT AGAACTCGAT GTAAAAGGAA AAAGAGTATT GTTAGTAGAT
 7771 GACGTTGCAA CTACAGGAGT ATCCATAGAG AAAGCAACAT TGGAGATTCT TAACGGTGGA GGTAAAGTTT
 7841 CAGACGCACT AGTAATCATA GATAGACAAG AAGGGGCTTC ACAAAGATTG GAAAAACTAG GAGTCAAATT
 7911 ACACTCTCTA TTTAAAATTT CAGAAATTCT AGATGAATTG TTGAAGAGTG ATAAACTTAA GGATAATGAA
 7981 AAGAAGTCAA TTCTAGATTA TTTGGTGAAG AATGTTGAAA AGTAGAGTAA TATTAGCAAT GGATAAACCT
 8051 CTCTCATATC AAGTTCTTAA AGAGATGGAA AATGAGTTAT ATGGGATAAA AGTTGGTTTA CCTTTAGTTT
 8121 TAGATCTAGG AGTGGATAAA ACTAGAGAGC TCTTAATTGG TTTAGACGTG GAGGAAATTA TTGTTGATTT
 8191 TAAGCTTGCA GATATCGGAT ACATAATGAA AAGCATAGTT GAAAGATTAT CTTTCGCCAA CTCGTTCATA
 8261 GCACATTCCT TTATAGGCGT TAAGGGATCT CTAGATGAAT TAAAAAGATA TCTTGATGCA AACTCTAAAA
 8331 ATTTATACTT AGTTGCCGTA ATGTCACATG AAGGATGGAG TACGTTATTC GCAGACTATA TTAAAAACGT
 8401 TATAAGAGAG ATAAGCCCAA AAGGAATAGT AGTTGGAGGG ACTAAATTAG ATCATATAAC GCAGTATAGG
 8471 AGAGACTTCG AAAAAATGAC CATAGTCTCT CCGGGTATGG GTAGTCAAGG TGGAAGTTAT GGCGATGCAG
 8541 TATGTGCTGG AGCGGATTAT GAAATCATTG GAAGAAGTAT TTATAATGCA GGGAATCCAT TAACAGCATT
 8611 AAGAACTATA AATAAGATTA TCGAGGATAA GGTGATGAAA TGTAAAGGAG CAATTTTTAG GAAAAAATGA
 8681 AATTAGTGAA AAAATAGCCG GTCTTTATAA ACTAACATTA ATAGGGGAAA ATCTACACGA GGTACCGTAA
 8751 GGAGGTTCCT GAAATAGCAT AATTTATAAC TAAAATATAA GGTAAAAAAT CTTTTTTATT GAGTATAAAC
 8821 TTCTATTTTT TCAGCTGAGA CGTCCTCCAC GAGTGCATTA TCTCCTCCTA TGAGGTAACT CCTACCACTC
 8891 CTTTCCTCTA CTAATTCGAA ATTAATCACT TCACTACTGA TATTAACGTT ATTGGGATAA CCAAATAACT
 8961 CAACATTTTC TCCTGTCCTG GTTATTACAC CATCCACCTT TACCTTTAAT CTTAATCCCT GCTCTTTAGC
 9031 CTTCATGATC TCCCATGTAG CAAATCTATG GGCACTAAAT ATCATAGGGA AATCACTTTG TGTCACAGCT
 9101 TTCCTGTAAA CAGTCCTAGA TGTCCTCCAT CCTTCGAAAT AGTAGTGAAT GAAGAAAAGA CTCATTGTCG
 9171 AGTCTCTAAA GACATAACTG TAAATGTTAG GATTCGTCGA GAGGCTCATG ACTCTCGGCA TAAACACGCT
 9241 TTCCTCTGCA TCAGCAGATA CCAAGAAAAA GTGTCCTAAA GAATGTACCT TTACTTCCAT CACTTTACTT
 9311 TCCAATAACT TGTTCACTAG AAACTCATCC ACATTAGGAT AAATATTTAG ATAAGTTTTC ACGCCTGACT
 9381 CTGCAACCTT AATAATGTAG CTGATTAGCT TCTGAAGGTA GTTATTAGGA ATTTCAACGT ATAACTCATT
 9451 TTTAGCACTC GTAATAACAG ATATACAGTT GTTCAATACG TTTTCAGTGC TCTTTGTCAT CCACATCATT
 9521 CCCTGAGATT TCCTGTGATT AGTTTCAAGC TTTTTGATCT GTTCCAGAAT AACATGTTTT CTCTTTGTCT
 9591 GCTCTTGGTC TATCTTTTCC AATACTATGT TTAACCTTGC TGGGATGTAG AATTTCGGCT TACTGTTCTG
 9661 AACATATATT AGACCTTTCC TCTCTAGGGA TAAAAGTATA TTGTAAAGCT GAGGCTGATG TATGTAAAGG
 9731 TCAGATATCA TCTCTTTAGC AGTAGCTCCT CCTTTATAAG TTAGATAAGC AAAAACTAAA GCTTCTTTCT
 9801 TCGATATTCC CAATACTGTA AGGGCATCTA TAAACTCATC ATTGACTTCA AACATCTATA TATAAAAGCT
 9871 CTGTTAGAAT TTTTAAATCT TTTTTTCTCT CCAAATCTTA ATTAATAATG AATTGTAGTA AAAATTTTGA
 9941 TATAGTGATA ACTATTTAAA TTAGAGAATC AATTAGCATA ATGTGGAGAT TAAAGCAGAG AGAAAAGGAA
10011 AGGTAGTAAG GGTACTCATA AACAACCCTA ATCCAGCTCG AGGGGGGGCC CGATTATTGA AGCATTTATC
10081 AGGGTTATTG TCTCATGAGC GGATACATAT TTGAATGTAT TTAGAAAAAT AAACAAATAG GGGTTCCGCG
10151 CACATTTCCC CGAAAAGTGC CACCTAAATT GTAAGCGTTA ATATTGAAAA AGGAAGAGTA TGAGTATTCA
10221 ACATTTCCGT GTCGCCCTTA TTCCCTTTTT TGCGGCATTT TGCCTTCCTG TTTTTGCTCA CCCAGAAACG
10291 CTGGTGAAAG TAAAAGATGC TGAAGATCAG TTGGGTGCAC GAGTGGGTTA CATCGAACTG GATCTCAACA
10361 GCGGTAAGAT CCTTGAGAGT TTCGCCCCG AAGAACGTTT TCCAATGATG AGCACTTTTA AAGTTCTGCT
```

```
10431 ATGTGGCGCG GTATTATCCC GTATTGACGC CGGGCAAGAG CAACTCGGTC GCCGCATACA CTATTCTCAG
10501 AATGACTTGG TTGAGTACTC ACCAGTCACA GAAAAGCATC TTACGGATGG CATGACAGTA AGAGAATTAT
10571 GCAGTGCTGC CATAACCATG AGTGATAACA CTGCGGCCAA CTTACTTCTG ACAACGATCG GAGGACCGAA
10641 GGAGCTAACC GCTTTTTTGC ACAACATGGG GGATCATGTA ACTCGCCTTG ATCGTTGGGA ACCGGAGCTG
10711 AATGAAGCCA TACCAAACGA CGAGCGTGAC ACCACGATGC CTGTAGCAAT GGCAACAACG TTGCGCAAAC
10781 TATTAACTGG CGAACTACTT ACTCTAGCTT CCCGGCAACA ATTAATAGAC TGGATGGAGG CGGATAAAGT
10851 TGCAGGACCA CTTCTGCGCT CGGCCCTTCC GGCTGGCTGG TTTATTGCTG ATAAATCTGG AGCCGGTGAG
10921 CGTGGGTCTC GCGGTATCAT TGCAGCACTG GGGCCAGATG GTAAGCCCTC CCGTATCGTA GTTATCTACA
10991 CGACGGGGAG TCAGGCAACT ATGGATGAAC GAAATAGACA GATCGCTGAG ATAGGTGCCT CACTGATTAA
11061 GCATTGGTAA CTGTCAGACC AAGTTTACTC ATATATACTT TAGATTGATT TAAAACTTCA TTTTTAATTT
11131 AAAAGGATCT AGGTGAAGAT CCTTTTTGAT AATCTCATGA CCAAAATCCC TTAACGTGAG TTTTCGTTCC
11201 ACTGAGCGTC AGACCCCGTA GAAAAGATCA AAGGATCTTC TTGAGATCCT TTTTTTCTGC GCGTAATCTG
11271 CTGCTTGCAA ACAAAAAAAC CACCGCTACC AGCGGTGGTT TGTTTGCCGG ATCAAGAGCT ACCAACTCTT
11341 TTTCCGAAGG TAACTGGCTT CAGCAGAGCG CAGATACCAA ATACTGTCCT TCTAGTGTAG CCGTAGTTAG
11411 GCCACCACTT CAAGAACTCT GTAGCACCGC CTACATACCT CGCTCTGCTA ATCCTGTTAC CAGTGGCTGC
11481 TGCCAGTGGC GATAAGTCGT GTCTTACCGG GTTGGACTCA AGACGATAGT TACCGGATAA GGCGCAGCGG
11551 TCGGGCTGAA CGGGGGGTTC GTGCACACAG CCCAGCTTGG AGCGAACGAC CTACACCGAA CTGAGATACC
11621 TACAGCGTGA GCTATGAGAA AGCGCCACGC TTCCCGAAGG GAGAAAGGCG GACAGGTATC CGGTAAGCGG
11691 CAGGGTCGGA ACAGGAGAGC GCACGAGGGA GCTTCCAGGG GGAAACGCCT GGTATCTTTA TAGTCCTGTC
11761 GGGTTTCGCC ACCTCTGACT TGAGCGTCGA TTTTTGTGAT GCTCGTCAGG GGGGCGGAGC CTATGGAAAA
11831 ACGCCAGCAA CGCGGCCTTT TTACGGTTCC TGGCCTTTTG CTGGCCTTTT GCTCACATGT TCTTTCCTGC
11901 GTTATCCCCT GATTCTGTGG ATAACCGTAT TACCGCCTTT GAGTGAGCTG ATACCGCTCG CCGCAGCCGA
11971 ACGACCGAGC GCAGCGAGTC AGTGAGCGAG GAAGCCCACC GCGG
```

**Supplementary Table 5:** Plasmid sequence map of pJL1601. The $P_{malE\_pJL1601}$ promoter sequence is highlighted in yellow and can be substituted by the promoter sequences in Supplementary Table 6 to result in the plasmid maps of pRN1-$P_{Saci\_2137}$, pRN1-$P_{Saci\_2137core}$, pRN1-$P_{sac7d}$ and pRN1-$P_{Ec10\_Sa1\text{-}Ec3\_Sa7}$. CDS: coding sequence, Orf: open reading frame, Ori: origin of replication.

| **Plasmid** | **pJL1601** | **10735 bp DNA circular** |
|---|---|---|
| **Features** | **Qualifiers** | **Location** |
| Promoter | PmalE | 0..264 |
| CDS | LacS | 264..1734 |
| Ori | pRN1 | 1779..7138 |
| Orf | orf90b | 1779..1892 |
| Orf | orf56 | 2165..2336 |
| Orf | orf904 | 2316..5031 |
| Orf | orf80 | 5872..6115 |
| Orf | orf90a | 6328..6601 |
| Orf | orf72 | 6708..6927 |
| Orf | orf90b | 6969..7138 |
| misc_feature | End of pyrB | 7195..7372 |
| CDS | PyrE | 7437..8025 |
| CDS | PyrF | 8011..8680 |
| CDS | $Amp^R$ | 8930..9791 |
| Ori | pMB1 | 9961..10550 |

```
   1 CCAGATATCT GATAGTTGGA GAAATGCTGA TGATTGCACG CTTATCTTTT TTTACCTACT GCTTGTGGTT
  71 TAAAAATTTT ACTGGAAACT AGAAATAAAT TTAAGTAAAT TAAATAAAGG CTAATTAATA ATACTAATAT
 141 CATTTCGTAA ACTACATCTT ATAATACTTA AGTTGACATG TTCAACGGAG GTGTCCTTAA GTTTAGACCT
 211 TAAATATTTT ATATATATAT TAAGTTTATA AATAATTACG TGATTAAGTT AACCATGGAC TCATTTCCAA
 281 ATAGCTTTAG GTTTGGTTGG TCCCAGGCCG GATTTCAATC AGAAATGGGA ACACCAGGGT CAGAAGATCC
 351 AAATACTGAC TGGTATAAAT GGGTTCATGA TCCAGAAAAC ATGGCAGCGG GATTAGTAAG TGGAGATCTA
 421 CCAGAAAATG GGCCAGGCTA CTGGGGAAAC TATAAGACAT TTCACGATAA TGCACAAAAA ATGGGATTAA
 491 AAATAGCTAG ACTAAATGTG GAATGGTCTA GGATATTTCC TAATCCATTA CCAAGGCCAC AAAACTTTGA
 561 TGAATCAAAA CAAGATGTGA CAGAGGTTGA GATAAACGAA AACGAGTTAA AGAGACTTGA CGAGTACGCT
 631 AATAAAGACG CATTAAACCA TTACAGGGAA ATATTCAAGG ATCTTAAAAG TAGAGGACTT TACTTTATAC
 701 TAAACATGTA TCATTGGCCA TTACCTCTAT GGTTACACGA CCCAATAAGA GTAAGAAGAG GAGATTTTAC
 771 TGGACCAAGT GGTTGGCTAA GTACTAGAAC AGTTTACGAA TTCGCTAGAT TCTCAGCTTA TATAGCTTGG
 841 AAATTCGATG ATCTAGTGGA TGAGTACTCA ACAATGAATG AACCTAACGT TGTTGGAGGT TTAGGATACG
 911 TTGGTGTTAA GTCCGGTTTT CCCCCAGGAT ACCTAAGCTT TGAACTTTCC CGTAGGGCAA TGTATAACAT
 981 CATTCAAGCT CACGCAAGAG CGTATGATGG GATAAAGAGT GTTTCTAAAA AACCAGTTGG AATTATTTAC
1051 GCTAATAGCT CATTCCAGCC GTTAACGGAT AAAGATATGG AAGCGGTAGA GATGGCTGAA AATGATAATA
1121 GATGGTGGTT CTTTGATGCT ATAATAAGAG GTGAGATCAC CAGAGGAAAC GAGAAGATTG TAAGAGATGA
1191 CCTAAAGGGT AGATTGGATT GGATTGGAGT TAATTATTAC ACTAGGACTG TTGTGAAGAG GACTGAAAAG
1261 GGATACGTTA GCTTAGGAGG TTACGGTCAC GGATGTGAGA GGAATTCTGT AAGTTTAGCG GGATTACCAA
1331 CCAGCGACTT CGGCTGGGAG TTCTTCCCAG AAGGTTTATA TGACGTTTTG ACGAAATACT GGAATAGATA
1401 TCATCTCTAT ATGTACGTTA CTGAAAATGG TATTGCGGAT GATGCCGATT ATCAAAGGCC CTATTATTTA
1471 GTATCTCACG TTTATCAAGT TCATAGAGCA ATAAATAGTG GTGCAGATGT TAGAGGGTAT TTACATTGGT
1541 CTCTAGCTGA TAATTACGAA TGGGCTTCAG GATTCTCTAT GAGGTTTGGT CTGTTAAAGG TCGATTACAA
1611 CACTAAGAGA CTATACTGGA GACCCTCAGC ACTAGTATAT AGGGAAATCG CCACAAATGG CGCAATAACT
1681 GATGAAATAG AGCACTTAAA TAGCGTACCT CCAGTAAAGC CATTAAGGCA CTAAGCGGCC GCTCTAGAAC
1751 TAGTGGATCC AGATGTGTAT AAGAGACAGA CTTTATCACG CCCGCCTTTG CTTCTGTTTC TTACTCTCAA
1821 CTGTTTCTAA CTCTGCTAAC TCACAATCCC GCTAATCACA CTTGTGACTT TACTAACTCT GCAGCTTTCT
1891 AACTAAGCCC GCCCTGTCTA ACTCTGCTAA CTATCCCCCG CCTTAATCAC ACCCGTGATT TTGTTTTTCT
1961 AACTCTGCAG CTTTGCTTCT TACTTACAGC TTTCTACCTA ACTCTTCTAA TCTCTCTACT TTCCTAACTC
2031 TCTGCCCCCC CCCCCCCCCC CGAACCCGAC GAAATAGAAA GTTAATTGCA GGTATTAGCT TTTCACACGG
2101 GTAATACTGA AAAATCTACC GTTGTCAATT TTAAGGATAA TTGCGGATAC AATTTTGATC CACAAATGGG
2171 TAGACCATAC AAACTATTAA ATGGAATAAA ATTAGGAGTC TATATTCCAC AGGAATGGCA TGACAGATTA
```

2241 ATGGAAATCG CTAAGGAAAA GAATCTTACG TTAAGTGATG TGTGCAGGCT GGCAATTAAA GAGTATTTAG
2311 ACAATCATGA TAAACAAAAG AAGTAAGGTA ATTCTTCATG GAAATGTGAA AAAAACAAGA AGAACGGGGG
2381 TGTACATGAT TAGTTTAGAT AATTCAGGTA ATAAAGATTT CTCTTCGAAT TTCTCAAGTG AGAGAATTAG
2451 ATATGCTAAG TGGTTTCTAG AACACGGCTT TAATATTATC CCAATCGACC CTGAGAGCAA AAAACCTGTA
2521 CTGAAGGAGT GGCAAAAGTA TAGTCATGAA ATGCCTTCTG ATGAGGAGAA ACAGAGATTT CTAAAGATGA
2591 TTGAAGAGGG GTATAATTAC GCTATTCCAG GGGGACAAAA AGGACTAGTG ATTCTCGATT TTGAGAGTAA
2661 GGAGAAGTTG AAAGCATGGA TAGGAGAATC AGCGTTAGAA GAGCTCTGTA GAAAAACATT ATGCACAAAC
2731 ACGGTTCATG GCGGAATCCA CATCTATGTT TTAAGTAACG ACATTCCGCC ACACAAAATC AATCCTCTCT
2801 TTGAAGAGAA TGGTAAGGGC ATAATTGATT TACAAAGTTA TAACAGTTAC GTTTTAGGAC TTGGCTCTTG
2871 TGTGAACCAT TTACACTGTA CTACTGATAA ATGCCCGTGG AAAGAACAGA ACTACACTAC TTGTTATACA
2941 TTATATAATG AATTAAAGGA AATTAGTAAA GTTGACTTAA AGAGTCTCCT AAGATTCCTG GCGGAAAAGG
3011 GCAAACGTTT AGGCATAACA TTAAGCAAAA CGGCAAAGGA GTGGTTAGAA GGAAAGAAGG AAGAAGAGGA
3081 CACGGTGGTG GAGTTTGAGG AGTTGAGAAA GGAATTAGTC AAACGTGACA GTGGGAAACC AGTTGAGAAA
3151 ATTAAAGAGG AGATATGTAC TAAGAGCCCG CCTAAACTGA TTAAGGAAAT AATATGTGAA AACAAGACTT
3221 ACGCTGATGT AAATATTGAT AGGAGTAGAG GGGATTGGCA TGTTATACTC TACTTAATGA AACACGGGGT
3291 TACGGATCCA GACAAGATAT TGGAATTATT ACCGCGTGAT TCAAAAGCAA AAGAAAATGA AAAGTGGAAT
3361 ACACAGAAAT ATTTTGTTAT AACTCTTTCG AAAGCCTGGA GTGTAGTAAA GAAGTACTTA GAAGCTAAAA
3431 GGAAAGCACA AAAAGACAAA AGCACGGCTA AGGCTCTGTT AATAGAGGCG ATTGCTGAAG AGGTGTTACA
3501 TGAACATTTT CTCGTTACCT TTATACAGAC TGATCAATTG AAGGAGTCAA AAATTGGGCT TTTCAGATTC
3571 AACAAGAAGA AGGGGATTTT CGAACCATTT GATGAAAGAA TTGAGAAGAT AATTAATGTA AAACTTGAGG
3641 AGTATAAGGA ATTCCCCCTG GGTTCTGATA AATCACGTGT GATACGTAAT ATTAAAGAAG AGATTATGAG
3711 GAGAACACAG AGACTGTTAT TAGAAGAGTC ACTTAGAATT GCGTTTCGTA ATGGGACACT GGAATGGGAT
3781 AGCAAAGGAG TAACATGGTA TGATGTTAAA GAAAGAACTC CTAAGGTGTA TTCATTTAAT TATGTGGATT
3851 GGAACTTAAA GATTGAGGAG ATTGAGAAGT TTAACATGAA GGAAATAACA GTTGAAGATA TAGAGAATTT
3921 AGCCCGGCGT GTTTGCCCTA GATCACTTGA GACTTTTAAG CAGTGGGTTG ATGATAAATG GGTGTTATTG
3991 TTTGAAGTCA TAGGCTATAC ATTCTATCCA AAGTACATCT TTAATAAGGC TATACTTCTC ACGGGGGCTG
4061 GGGCAAACGG GAAATCGACT TTCCTTAATT TGCTGTTAAA GATTCTAGGA CAGAAGAACG TTTCCGCAAT
4131 GCCTTTGAAG AGGATCATGG AGAGTGATAG ATTTGCTTCA ATTGAACTAT TCCATAAACT AGCAAATGTA
4201 AGCAGTGAGT TGTTTGCCTT TAAGATCACA AACACAGACC TGTTTAAAAA ACTCACGGGG GAAGACTACA
4271 TAGAGGGACA GAAGAAGTTC AGAGACCCCA TATACTTCAT TAATTACGCA AAACTATTCA ACGCCACTAA
4341 CGAATTGCCC GTGGTTTCTG ATCAAAGTTA TGGATTCTGG AGAAGATGGA TTGTGATAGA ATTTCCACAC
4411 CAATTCCCGC CGGATCCTAA CTTTTTCGAT AAGACATTCA CGGTGGAGGA GGTTGAAGGA GTAATCACGG
4481 TAGCTGTTAT CGCATTCGCC CGTGTACTTC AACAGAAGAA ATTCGACTTT GAGGATAGCA GTGCTAATGT
4551 CAAAGAATTA TGGGAAAGAA AAACAGACAG TGTTTACGCA TTCGTGAAAG AGTTACTAGA AACCGAAAGG
4621 GCTGAATATG ATCCGGCAAA CGGCGATTTA TTCATGCCCA CTGAGGACTT CTACCAGGCT TATTTAGAAT
4691 GGAGCGAAGA GAACGACACA AAAGCCGAAA GCAAAGCCGT GGTAACCCAA AGATTGCAAA GCAAATTTAG
4761 AATAACAAAA GACAAGAAGA AGATTAATGG AAAAAGAGTT TGGTGTTATG TGGGAATAAG ATTGAAGAAT
4831 AACAATATAA GTACGGGGGG CGGACAGGAC GGCGATTCTC CTAATTCTTT ATTAGAATTA TATAAGGAAT
4901 TCCAGGGCAA AGTGGTAAGT AGGAAAGACT TATATGATAT GCTAAAGCTA AGAGCGTTTG AGTTGTTGGA
4971 GTTGTGCGAG AAGAGAAACA TATGTCACTG GATTGATGAA GAGCACGTGC GGTTTGATTG AGTCCTTCAA
5041 GTTTTCAATT TTTTAAATTG AATTTTTCAT CTGTAATGAC CAATTTATGT CCATAGTGTC CAACTTTTTT
5111 TCTATTATTT TGATACACGG TGGGACAATA ATATAATAAA ATAATGCCTT TTTAGTTGGA CACTATGGAC
5181 ATAAGTAGTC ACACCCGTGA TAATATTTGT ATAGTAATGG CGTTTTTCAA ATATTCTTAC AACAAAAATT
5251 TGAATTTATC TAACGTCATT CTCTCTATAG AACCATTCTC TTTATAGAGC TATACTTTAT TCTTATTAAA
5321 AATTTAGAAA TGGGGAATTT CTAAAAACTG GAATTCCCCA TTTCCTCTAG ACTCTTCTTT TTCTTCTCTT
5391 CTCTCTCCTG TCTTGAGTCT AGAGTCCATA AACACTGCTG CATAACAGCA TGACAACCGT GATAAAGTCT
5461 GAGTTCAAGA TACAAGATAT TTATAAGGAG TAGGCATTTT TTTCTCTCTT CTTAAAATAT TTTTCCAATC
5531 GGTTATCGAT ATTCTGGCAC TAGTGCCAGA AATTTACTCA AAAGAAGTAC AAAAATTCTC GCCACTTGGC
5601 GAGAAATTTG CTCAAAGTAG TGAAACAATA TGAAAAAGAA AGAGAACAAT CACAGATATG ATTAAACTGC
5671 TGCATGCAGC CACGAAATCA CAACTATGAT TATGCAGTCC ATATGTTAAT CCTGGTCGGA TCCGCAAAAT
5741 TTTAGTTATA AGAGTTAGCT AACACGATAA GGCAAACAGT ATTAATAAAG CGTTAATCCT ACCTCCACCG
5811 TGTTATTTAG CTAACTTTTT GCACGCCAAA AGATATTTAA CAGTCTGTTA ATCCTACTTT ACATGGGATC
5881 CCATATGAGT GATCTGAAGG AAAAGCTAAC TCTAACTCAA CTAATCCTGA TTCGGCTATC AAAATCTTGT
5951 CAAACCCTGG AAGAGTTAGA ACGATATACA GGTGCAAACA GAAATGTACT TCTCGTTACC TTGACACGAC
6021 TCCATAAAAA AGGCATAATC TACAGGAAAT GGCGTAGGTT TGGCGGTAGG AAGTATAGAG AATATTGTTT
6091 GAAAAGCCGT GACGAACTCC TGTGAAACTC CCCAGTTTAC CGTTATTATT GATACATATC GATACATAAT
6161 GATACATATG CACACATAAT GATACAGAAT AAGCGTAGAA TAATAGTATA AGCATAGTAT AAGCACATGA
6231 TAAGAGTATA AAATAGGCGT AAAAGCGTAC GGCTTTCAGC TATATATCCC CAAACCTACT ATTTTTCCAC

```
6301 GCTCCCTTAA ACAAAATTCT AAGATAATTT GTTTTACTTT ACTGTATATT ATTCAATGCA TCTAAAATTT
6371 TACTGTATAC GTCTATCGTT TTACCTGATG CGTATACCAT TTTCCCCCAT GCACCGCATT TCGTCTATAT
6441 ACTCCCATTC TTCTATTTTT TGCACATTTC ACGCAAAAAA CTTCCCATAC CCAATCCCGC CGTGCTTTTT
6511 TTCCTTTTAC TGCTTACTCT TTCACATAGC CCCCTCTGCT TCTCTAACTC TGCAGTTTTG CTCAGGCTTT
6581 TGCTTAATCT CTCTCACATA GCTCTTCTTC TTTCTGCTTT TCTAACTCTA CTTTCTTTCT TTTCCTCTAC
6651 TTAACTACTC TTTCACTTCT ACTTTTCTAA CTCTTCTTTT GCTCTTCTCT TCTTAGCTTT GCTAACTATG
6721 TTTCCTTTCT ACTTTCATTA TCTCTTCTTA GCTTTTGCTC TCTTTCTTTA CTCTTTATCT ACAGCTTCAC
6791 TCTTCTTCTT CTTTTTCCAA TCTCTGATTT TTCTCACGAT TTCTAGACAA ATTCCCGTAA CCCCCAATCC
6861 CCCCGTGCTA ACCCAATCTT TGATTATCTT AGACAATCTT TCCCTATGTT TGACTTTCTT AGACTAACTT
6931 AGACAGTTTC AAACAAACTA TTCCTACTTA TGCCGACTTA TGCCGACTTC TAACAACATC TTACGACATC
7001 TTATGACATC TTACGACATC TTACAAATTT TAGGACTTTC ATTTCTATAC CCAAACTCTC TGTTTTTTTG
7071 CACTTTCGTG CAAACAATTC TATATTCCAA TTCCAGACTC CTATACCCCC CATGCTTCTA CTTTATCACT
7141 GTCTCTTATA CACATCTTCG AGGGGGGGCC GCTCTAGAAC TAGTGGATCC CCCGGGCTGC AGTATGAAAG
7211 CTTTGAGCAG TTCTAGTACT TGGCTCAAAG AATGCTAATG AAACTACTTT CCCTGATAGA TAATTTAGTT
7281 TTTTTACATC ATAGAACTTG TCTGCAAGTT CGAATATTCT AAAGTAGTCA TCTCTGGTCA AGTCAAGCGA
7351 CGAAACTACG TGTCTTAATC TCACAAAGCC CTTATAACTG GTTATGATAG AAGTATATTT AAATTCTTTT
7421 TCACAGAGTC TCTACGTATG AATTTCGCAG AAGTCTTACT CGAAAGGAAA TTATTATTAA TAGGAAGTTT
7491 CGTTTTAACA TCAGGTAAGG TTAGTCCATA TTACTTAGAC TTAAGACCTT TACCAAATTA TCCAGAATTT
7561 TACGATATAG TTAATCAAGC TATAAAGAAA GCAAAAGATA TACCCCATGA TATAATAGTA GGAATAGCCA
7631 CTGGAGGAGT TCCCTTATCG GCATTCATAG CTTGTAACCT TAAAGAGCCT ATGGGATATA TTAGAATAGA
7701 AAAGAAAGGT CATGGAACTA ATCGTACATT AGAACTCGAT GTAAAAGGAA AAAGAGTATT GTTAGTAGAT
7771 GACGTTGCAA CTACAGGAGT ATCCATAGAG AAAGCAACAT TGGAGATTCT TAACGGTGGA GGTAAAGTTT
7841 CAGACGCACT AGTAATCATA GATAGACAAG AAGGGGCTTC ACAAAGATTG GAAAAACTAG GAGTCAAATT
7911 ACACTCTCTA TTTAAAATTT CAGAAATTCT AGATGAATTG TTGAAGAGTG ATAAACTTAA GGATAATGAA
7981 AAGAAGTCAA TTCTAGATTA TTTGGTGAAG AATGTTGAAA AGTAGAGTAA TATTAGCAAT GGATAAACCT
8051 CTCTCATATC AAGTTCTTAA AGAGATGGAA AATGAGTTAT ATGGGATAAA AGTTGGTTTA CCTTTAGTTT
8121 TAGATCTAGG AGTGGATAAA ACTAGAGAGC TCTTAATTGG TTTAGACGTG GAGGAAATTA TTGTTGATTT
8191 TAAGCTTGCA GATATCGGAT ACATAATGAA AAGCATAGTT GAAAGATTAT CTTTCGCCAA CTCGTTCATA
8261 GCACATTCCT TTATAGGCGT TAAGGGATCT CTAGATGAAT TAAAAAGATA TCTTGATGCA AACTCTAAAA
8331 ATTTATACTT AGTTGCCGTA ATGTCACATG AAGGATGGAG TACGTTATTC GCAGACTATA TTAAAAACGT
8401 TATAAGAGAG ATAAGCCCAA AAGGAATAGT AGTTGGAGGG ACTAAATTAG ATCATATAAC GCAGTATAGG
8471 AGAGACTTCG AAAAAATGAC CATAGTCTCT CCGGGTATGG GTAGTCAAGG TGGAAGTTAT GGCGATGCAG
8541 TATGTGCTGG AGCGGATTAT GAAATCATTG GAAGAAGTAT TTATAATGCA GGGAATCCAT TAACAGCATT
8611 AAGAACTATA AATAAGATTA TCGAGGATAA GGTGATGAAA TGTAAAGGAG CAATTTTTAG GAAAAAATGA
8681 AATTAGTGAA AAAATAGCCG GTCTTTATAA ACTAACATTA ATAGGGGAAA ATCTACACGA GGTACCCTCT
8751 GTTAGAATTT TTAAATCCTC GAGGGGGGGC CCGATTATTG AAGCATTTAT CAGGGTTATT GTCTCATGAG
8821 CGGATACATA TTTGAATGTA TTTAGAAAAA TAAACAAATA GGGGTTCCGC GCACATTTCC CCGAAAAGTG
8891 CCACCTAAAT TGTAAGCGTT AATATTGAAA AAGGAAGAGT ATGAGTATTC AACATTTCCG TGTCGCCCTT
8961 ATTCCCTTTT TTGCGGCATT TTGCCTTCCT GTTTTTGCTC ACCCAGAAAC GCTGGTGAAA GTAAAAGATG
9031 CTGAAGATCA GTTGGGTGCA CGAGTGGGTT ACATCGAACT GGATCTCAAC AGCGGTAAGA TCCTTGAGAG
9101 TTTTCGCCCC GAAGAACGTT TTCCAATGAT GAGCACTTTT AAAGTTCTGC TATGTGGCGC GGTATTATCC
9171 CGTATTGACG CCGGGCAAGA GCAACTCGGT CGCCGCATAC ACTATTCTCA GAATGACTTG GTTGAGTACT
9241 CACCAGTCAC AGAAAAGCAT CTTACGGATG GCATGACAGT AAGAGAATTA TGCAGTGCTG CCATAACCAT
9311 GAGTGATAAC ACTGCGGCCA ACTTACTTCT GACAACGATC GGAGGACCGA AGGAGCTAAC CGCTTTTTTG
9381 CACAACATGG GGGATCATGT AACTCGCCTT GATCGTTGGG AACCGGAGCT GAATGAAGCC ATACCAAACG
9451 ACGAGCGTGA CACCACGATG CCTGTAGCAA TGGCAACAAC GTTGCGCAAA CTATTAACTG GCGAACTACT
9521 TACTCTAGCT TCCCGGCAAC AATTAATAGA CTGGATGGAG GCGGATAAAG TTGCAGGACC ACTTCTGCGC
9591 TCGGCCCTTC CGGCTGGCTG GTTTATTGCT GATAAATCTG GAGCCGGTGA GCGTGGGTCT CGCGGTATCA
9661 TTGCAGCACT GGGGCCAGAT GGTAAGCCCT CCCGTATCGT AGTTATCTAC ACGACGGGGA GTCAGGCAAC
9731 TATGGATGAA CGAAATAGAC AGATCGCTGA GATAGGTGCC TCACTGATTA AGCATTGGTA ACTGTCAGAC
9801 CAAGTTTACT CATATATACT TTAGATTGAT TTAAAACTTC ATTTTTAATT TAAAAGGATC TAGGTGAAGA
9871 TCCTTTTTGA TAATCTCATG ACCAAAATCC CTTAACGTGA GTTTTCGTTC CACTGAGCGT CAGACCCCGT
9941 AGAAAAGATC AAAGGATCTT CTTGAGATCC TTTTTTTCTG CGCGTAATCT GCTGCTTGCA AACAAAAAAA
10011 CCACCGCTAC CAGCGGTGGT TTGTTTGCCG GATCAAGAGC TACCAACTCT TTTTCCGAAG GTAACTGGCT
10081 TCAGCAGAGC GCAGATACCA AATACTGTCC TTCTAGTGTA GCCGTAGTTA GGCCACCACT TCAAGAACTC
10151 TGTAGCACCG CCTACATACC TCGCTCTGCT AATCCTGTTA CCAGTGGCTG CTGCCAGTGG CGATAAGTCG
10221 TGTCTTACCG GGTTGGACTC AAGACGATAG TTACCGGATA AGGCGCAGCG GTCGGGCTGA ACGGGGGGTT
10291 CGTGCACACA GCCCAGCTTG GAGCGAACGA CCTACACCGA ACTGAGATAC CTACAGCGTG AGCTATGAGA
```

```
10361 AAGCGCCACG CTTCCCGAAG GGAGAAAGGC GGACAGGTAT CCGGTAAGCG GCAGGGTCGG AACAGGAGAG
10431 CGCACGAGGG AGCTTCCAGG GGGAAACGCC TGGTATCTTT ATAGTCCTGT CGGGTTTCGC CACCTCTGAC
10501 TTGAGCGTCG ATTTTTGTGA TGCTCGTCAG GGGGGCGGAG CCTATGGAAA AACGCCAGCA ACGCGGCCTT
10571 TTTACGGTTC CTGGCCTTTT GCTGGCCTTT TGCTCACATG TTCTTTCCTG CGTTATCCCC TGATTCTGTG
10641 GATAACCGTA TTACCGCCTT TGAGTGAGCT GATACCGCTC GCCGCAGCCG AACGACCGAG CGCAGCGAGT
10711 CAGTGAGCGA GGAAGCCCAC CGCGG
```

**Supplementary Table 6:** Overview of the promoter sequences characterised in this chapter. For the backbone sequence of the plasmids see Supplementary Table 3 for *Escherichia coli* plasmids or Supplementary Table 5 for *Sulfolobus acidocaldarius* plasmids. In these sequences the sequence that is substituted with the respective promoter sequences is highlighted in yellow. The non-capitalised base pairs in for $P_{sac7d}$ represent the cloning scar that was present in the original plasmid from Berkner *et al.* [21].

| Promoter | Sequence | Source |
|---|---|---|
| $P_{sac7d}$ | CCCTCACTATAACTAGCTAGTTTAAGGTTTATTAATTTGTTGCTTGATCTTACCAA<br>CATACGTTGATCTAGATATTATGTTGAATTCTTATGTTCTATAGCGTAATTATGAA<br>CAGTTTGTATAACTCCTTTAGAGAATAAATTATATTTCAATATTACTAATTATTGT<br>ACTGGATTCCCCATAAAATTGTATACATTATATAGGAAAAATAATTTGAGGTAGTC<br>TCATAAGTATGACTTAACTTAATACCGTAAGGTTTATTTATGACAATATCGTAAGA<br>TAACTTGACCTAcc | [21] |
| $P_{Saci\_2137core}$ | TATAATACAAATCACTCTCAAAGAGGTTTATAATCTGTTTAAGAAGAATATTACTT | This study |
| $P_{Ec10\text{-}Sa1}$ | TATAATACAAATCACTCTCAAATGGTTTTATAAACTGAGGGGAGAGAATATTACTT | This study |
| $P_{Ec6\text{-}Sa2}$ | TATAATACAAATCACTCTCAAATGTCTTTATAAAACGGGCTTGGAGAATATTACTT | This study |
| $P_{Ec7\text{-}Sa3}$ | TATAATACAAATCACTCTCAAACTCATTTATAAATGCAAGAAGGAGAATATTACTT | This study |
| $P_{Ec8\text{-}Sa4}$ | TATAATACAAATCACTCTCAAAATGCTTTATAAACGAACAGGATAGAATATTACTT | This study |
| $P_{Ec4\text{-}Sa5}$ | TATAATACAAATCACTCTCAAATCCGTTTATAAATTGAGTCAGGAGAATATTACTT | This study |
| $P_{Ec9\text{-}Sa6}$ | TATAATACAAATCACTCTCAAAGACGTTTATAAAAGGATGGAAAAGAATATTACTT | This study |
| $P_{Ec3\text{-}Sa7}$ | TATAATACAAATCACTCTCAAAATAGTTTATAATCATGCTTTTGAGAATATTACTT | This study |
| $P_{Ec1}$ | TATAATACAAATCACTCTCAAAAATTTTTATAAACAGATAGCGGAGAATATTACTT | This study |
| $P_{Ec2}$ | TATAATACAAATCACTCTCAAACTGGTTTATAATGGACATGCGGAGAATATTACTT | This study |
| $P_{Ec5}$ | TATAATACAAATCACTCTCAAAATGTTTTATAACGTGGGGTCGGAGAATATTACTT | This study |

**Supplementary Table 7:** Parameters determined by the model fitting. B_rep: biological replicate, T_rep: technical replicate, $t_{off}$: offset, m: maximum value, d: period around the offset in which the transition takes place (see Equations 2 and 3).

| Plate | Promoter | B_rep | T_rep | Slope and error $\times 10^3$ | $t_{off}$ | m | d |
|---|---|---|---|---|---|---|---|
| 1 | $P_{Ec10\_Sa1}$ | 1 | 1 | 1.733 ± 0.338 | 669 | 4.72 | 640 |
| 1 | $P_{Ec10\_Sa1}$ | 1 | 2 | 1.760 ± 0.157 | 769 | 5.70 | 674 |
| 1 | $P_{Ec10\_Sa1}$ | 1 | 3 | 1.787 ± 0.150 | 805 | 5.78 | 664 |
| 1 | $P_{Ec10\_Sa1}$ | 2 | 1 | 1.622 ± 0.227 | 907 | 5.64 | 707 |
| 1 | $P_{Ec10\_Sa1}$ | 2 | 2 | 1.508 ± 0.184 | 854 | 7.24 | 719 |
| 1 | $P_{Ec10\_Sa1}$ | 2 | 3 | 1.562 ± 0.171 | 875 | 6.97 | 714 |
| 1 | $P_{Ec10\_Sa1}$ | 3 | 1 | 1.849 ± 0.953 | 633 | 247.92 | 0 |
| 1 | $P_{Ec10\_Sa1}$ | 3 | 2 | 1.926 ± 1.025 | 640 | 258.87 | 0 |
| 1 | $P_{Ec10\_Sa1}$ | 3 | 3 | 1.885 ± 0.831 | 635 | 246.90 | 0 |
| 1 | $P_{Ec6\_Sa2}$ | 1 | 1 | 1.224 ± 0.076 | 956 | 4.27 | 800 |
| 1 | $P_{Ec6\_Sa2}$ | 1 | 2 | 1.226 ± 0.081 | 1068 | 5.45 | 798 |
| 1 | $P_{Ec6\_Sa2}$ | 1 | 3 | 1.173 ± 0.059 | 1283 | 4.76 | 875 |
| 1 | $P_{Ec6\_Sa2}$ | 2 | 1 | 0.931 ± 0.045 | 1422 | 5.02 | 1004 |
| 1 | $P_{Ec6\_Sa2}$ | 2 | 2 | 0.959 ± 0.039 | 1474 | 5.09 | 991 |
| 1 | $P_{Ec6\_Sa2}$ | 2 | 3 | 0.975 ± 0.059 | 1468 | 5.10 | 990 |
| 1 | $P_{Ec6\_Sa2}$ | 3 | 1 | 1.172 ± 0.054 | 1370 | 4.36 | 896 |
| 1 | $P_{Ec6\_Sa2}$ | 3 | 2 | 1.180 ± 0.086 | 1354 | 4.39 | 891 |
| 1 | $P_{Ec6\_Sa2}$ | 3 | 3 | 1.191 ± 0.045 | 1368 | 3.87 | 900 |
| 1 | $P_{Ec7\_Sa3}$ | 1 | 1 | 1.312 ± 0.068 | 1321 | 4.35 | 855 |
| 1 | $P_{Ec7\_Sa3}$ | 1 | 2 | 1.230 ± 0.046 | 1305 | 4.68 | 866 |
| 1 | $P_{Ec7\_Sa3}$ | 1 | 3 | 1.377 ± 0.050 | 1305 | 4.27 | 837 |
| 1 | $P_{Ec7\_Sa3}$ | 2 | 1 | 0.736 ± 0.018 | 1431 | 4.17 | 1118 |
| 1 | $P_{Ec7\_Sa3}$ | 2 | 2 | 0.711 ± 0.015 | 1356 | 3.91 | 1145 |
| 1 | $P_{Ec7\_Sa3}$ | 2 | 3 | 0.692 ± 0.021 | 1219 | 3.52 | 1125 |
| 1 | $P_{Ec7\_Sa3}$ | 3 | 1 | 1.121 ± 0.069 | 1075 | 3.93 | 865 |
| 1 | $P_{Ec7\_Sa3}$ | 3 | 2 | 1.160 ± 0.051 | 1203 | 4.84 | 864 |
| 1 | $P_{Ec7\_Sa3}$ | 3 | 3 | 0.771 ± 0.030 | 1533 | 4.88 | 1127 |
| 1 | $P_{Ec8\_Sa4}$ | 1 | 1 | 0.426 ± 0.012 | 1825 | 4.27 | 1606 |
| 1 | $P_{Ec8\_Sa4}$ | 1 | 2 | 0.374 ± 0.010 | 1622 | 4.49 | 1618 |
| 1 | $P_{Ec8\_Sa4}$ | 1 | 3 | 0.353 ± 0.011 | 1456 | 3.75 | 1650 |
| 1 | $P_{Ec8\_Sa4}$ | 2 | 1 | 0.674 ± 0.025 | 1252 | 2.77 | 1124 |
| 1 | $P_{Ec8\_Sa4}$ | 2 | 2 | 0.713 ± 0.024 | 1402 | 3.61 | 1128 |
| 1 | $P_{Ec8\_Sa4}$ | 2 | 3 | 1.284 ± 0.093 | 1161 | 4.95 | 818 |
| 1 | $P_{Ec8\_Sa4}$ | 3 | 1 | 1.610 ± 0.309 | 902 | 7.76 | 717 |
| 1 | $P_{Ec8\_Sa4}$ | 3 | 2 | 1.658 ± 0.034 | 920 | 6.87 | 0 |
| 1 | $P_{Ec8\_Sa4}$ | 3 | 3 | 1.781 ± 0.037 | 969 | 6.23 | 0 |
| 1 | $P_{Ec4\_Sa5}$ | 1 | 1 | 0.966 ± 0.027 | 1447 | 4.51 | 1020 |
| 1 | $P_{Ec4\_Sa5}$ | 1 | 2 | 0.925 ± 0.034 | 1419 | 4.79 | 1002 |
| 1 | $P_{Ec4\_Sa5}$ | 1 | 3 | 0.936 ± 0.037 | 1453 | 4.62 | 1010 |
| 1 | $P_{Ec4\_Sa5}$ | 2 | 1 | 0.620 ± 0.020 | 1655 | 5.38 | 1272 |
| 1 | $P_{Ec4\_Sa5}$ | 2 | 2 | 0.635 ± 0.014 | 1678 | 4.94 | 1276 |
| 1 | $P_{Ec4\_Sa5}$ | 2 | 3 | 0.641 ± 0.014 | 1672 | 4.75 | 1327 |
| 1 | $P_{Ec4\_Sa5}$ | 3 | 1 | 0.708 ± 0.016 | 1472 | 4.05 | 1179 |
| 1 | $P_{Ec4\_Sa5}$ | 3 | 2 | 0.692 ± 0.015 | 1376 | 3.89 | 1148 |
| 1 | $P_{Ec4\_Sa5}$ | 3 | 3 | 0.694 ± 0.016 | 1258 | 3.32 | 1130 |
| 1 | $P_{Ec9\_Sa6}$ | 1 | 1 | 0.665 ± 0.019 | 1857 | 2.61 | 1306 |
| 1 | $P_{Ec9\_Sa6}$ | 1 | 2 | 0.533 ± 0.019 | 1684 | 2.61 | 1414 |
| 1 | $P_{Ec9\_Sa6}$ | 1 | 3 | 0.529 ± 0.018 | 1673 | 2.60 | 1412 |
| 1 | $P_{Ec9\_Sa6}$ | 2 | 1 | 0.886 ± 0.034 | 1511 | 2.75 | 1084 |
| 1 | $P_{Ec9\_Sa6}$ | 2 | 2 | 0.814 ± 0.030 | 1395 | 3.04 | 1065 |
| 1 | $P_{Ec9\_Sa6}$ | 2 | 3 | 0.859 ± 0.024 | 1319 | 2.75 | 1066 |
| 1 | $P_{Ec9\_Sa6}$ | 3 | 1 | 0.477 ± 0.014 | 1300 | 1.90 | 1407 |

| Plate | Promoter | B_rep | T_rep | Slope and error $\times 10^3$ | $t_{off}$ | m | d |
|---|---|---|---|---|---|---|---|
| 1 | $P_{Ec9\_Sa6}$ | 3 | 2 | 0.458 ± 0.013 | 1399 | 2.49 | 1464 |
| 1 | $P_{Ec9\_Sa6}$ | 3 | 3 | 0.559 ± 0.020 | 1635 | 2.29 | 1477 |
| 1 | $P_{Ec3\_Sa7}$ | 1 | 1 | 0.261 ± 0.009 | 2378 | 4.17 | 2521 |
| 1 | $P_{Ec3\_Sa7}$ | 1 | 2 | 0.231 ± 0.011 | 3549 | 3.78 | 3227 |
| 1 | $P_{Ec3\_Sa7}$ | 1 | 3 | 0.252 ± 0.015 | 4080 | 1.91 | 3394 |
| 1 | $P_{Ec3\_Sa7}$ | 2 | 1 | 0.396 ± 0.012 | 2993 | 1.48 | 2303 |
| 1 | $P_{Ec3\_Sa7}$ | 2 | 2 | 0.396 ± 0.013 | 2839 | 2.50 | 2131 |
| 1 | $P_{Ec3\_Sa7}$ | 2 | 3 | 0.391 ± 0.011 | 2943 | 3.04 | 2114 |
| 1 | $P_{Ec3\_Sa7}$ | 3 | 1 | 0.362 ± 0.012 | 3008 | 3.56 | 2230 |
| 1 | $P_{Ec3\_Sa7}$ | 3 | 2 | 0.341 ± 0.011 | 2936 | 3.89 | 2233 |
| 1 | $P_{Ec3\_Sa7}$ | 3 | 3 | 0.389 ± 0.013 | 2859 | 4.04 | 2055 |
| 1 | $P_{malE\_pJL1601}$ | 1 | 1 | 0.437 ± 0.019 | 1765 | 1.20 | 1739 |
| 1 | $P_{malE\_pJL1601}$ | 1 | 2 | 0.369 ± 0.012 | 1563 | 1.47 | 1816 |
| 1 | $P_{malE\_pJL1601}$ | 1 | 3 | 0.330 ± 0.013 | 1364 | 1.56 | 1854 |
| 1 | $P_{malE\_pJL1601}$ | 2 | 1 | 0.283 ± 0.009 | 1313 | 1.12 | 2102 |
| 1 | $P_{malE\_pJL1601}$ | 2 | 2 | 0.285 ± 0.012 | 1359 | 1.23 | 2026 |
| 1 | $P_{malE\_pJL1601}$ | 2 | 3 | 0.288 ± 0.022 | 1389 | 1.25 | 2058 |
| 1 | $P_{malE\_pJL1601}$ | 3 | 1 | 0.363 ± 0.020 | 1719 | 1.18 | 1982 |
| 1 | $P_{malE\_pJL1601}$ | 3 | 2 | 0.385 ± 0.025 | 1823 | 1.16 | 1960 |
| 1 | $P_{malE\_pJL1601}$ | 3 | 3 | 0.399 ± 0.025 | 1797 | 1.16 | 1886 |
| 2 | $P_{Saci\_2137}$ | 1 | 1 | 0.912 ± 0.054 | 856 | 5.24 | 867 |
| 2 | $P_{Saci\_2137}$ | 1 | 2 | 0.964 ± 0.044 | 1002 | 6.06 | 869 |
| 2 | $P_{Saci\_2137}$ | 1 | 3 | 0.658 ± 4.363 | 90 | 289.62 | 1130 |
| 2 | $P_{Saci\_2137}$ | 2 | 1 | 0.717 ± 0.038 | 1263 | 6.70 | 1113 |
| 2 | $P_{Saci\_2137}$ | 2 | 2 | 0.740 ± 0.043 | 1304 | 6.33 | 1127 |
| 2 | $P_{Saci\_2137}$ | 2 | 3 | 0.696 ± 0.032 | 1299 | 5.90 | 1153 |
| 2 | $P_{Saci\_2137}$ | 3 | 1 | 0.575 ± 0.017 | 1352 | 4.99 | 1322 |
| 2 | $P_{Saci\_2137}$ | 3 | 2 | 0.562 ± 0.016 | 1298 | 4.51 | 1319 |
| 2 | $P_{Saci\_2137}$ | 3 | 3 | 0.643 ± 0.022 | 1336 | 4.27 | 1273 |
| 2 | $P_{Saci\_2137core}$ | 1 | 1 | 1.654 ± 0.135 | 871 | 5.81 | 656 |
| 2 | $P_{Saci\_2137core}$ | 1 | 2 | 1.547 ± 0.158 | 799 | 6.55 | 641 |
| 2 | $P_{Saci\_2137core}$ | 1 | 3 | 1.496 ± 0.158 | 699 | 6.21 | 635 |
| 2 | $P_{Saci\_2137core}$ | 2 | 1 | 1.522 ± 0.090 | 803 | 5.45 | 651 |
| 2 | $P_{Saci\_2137core}$ | 2 | 2 | 1.532 ± 0.119 | 923 | 6.44 | 660 |
| 2 | $P_{Saci\_2137core}$ | 2 | 3 | 1.626 ± 0.171 | 1021 | 6.29 | 679 |
| 2 | $P_{Saci\_2137core}$ | 3 | 1 | 1.763 ± 0.266 | 949 | 7.66 | 648 |
| 2 | $P_{Saci\_2137core}$ | 3 | 2 | 1.689 ± 0.249 | 957 | 8.16 | 656 |
| 2 | $P_{Saci\_2137core}$ | 3 | 3 | 1.671 ± 0.200 | 943 | 8.98 | 654 |
| 2 | $P_{malE\_pSVA1450}$ | 1 | 1 | 0.849 ± 0.034 | 1805 | 3.82 | 1179 |
| 2 | $P_{malE\_pSVA1450}$ | 1 | 2 | 0.802 ± 0.025 | 1766 | 3.69 | 1150 |
| 2 | $P_{malE\_pSVA1450}$ | 1 | 3 | 0.795 ± 0.027 | 1734 | 3.71 | 1180 |
| 2 | $P_{malE\_pSVA1450}$ | 2 | 1 | 0.833 ± 0.031 | 1673 | 3.85 | 1189 |
| 2 | $P_{malE\_pSVA1450}$ | 2 | 2 | 0.777 ± 0.024 | 1576 | 3.89 | 1134 |
| 2 | $P_{malE\_pSVA1450}$ | 2 | 3 | 0.687 ± 0.024 | 1418 | 3.18 | 1189 |
| 2 | $P_{malE\_pSVA1450}$ | 3 | 1 | 0.634 ± 0.034 | 1347 | 2.20 | 1297 |
| 2 | $P_{malE\_pSVA1450}$ | 3 | 2 | 0.585 ± 0.026 | 1428 | 2.90 | 1302 |
| 2 | $P_{malE\_pSVA1450}$ | 3 | 3 | 0.655 ± 0.027 | 1632 | 2.92 | 1261 |
| 2 | $P_{sac7d}$ | 1 | 1 | 1.779 ± 0.158 | 1054 | 5.93 | 669 |
| 2 | $P_{sac7d}$ | 1 | 2 | 1.859 ± 0.199 | 1060 | 6.27 | 669 |
| 2 | $P_{sac7d}$ | 1 | 3 | 1.762 ± 0.335 | 1051 | 6.58 | 660 |
| 2 | $P_{sac7d}$ | 2 | 1 | 1.997 ± 0.249 | 959 | 7.98 | 640 |
| 2 | $P_{sac7d}$ | 2 | 2 | 1.920 ± 0.091 | 914 | 10.07 | 555 |
| 2 | $P_{sac7d}$ | 2 | 3 | 2.001 ± 0.199 | 926 | 7.76 | 606 |
| 2 | $P_{sac7d}$ | 3 | 1 | 1.774 ± 0.198 | 929 | 8.40 | 624 |
| 2 | $P_{sac7d}$ | 3 | 2 | 1.881 ± 0.204 | 883 | 7.14 | 603 |
| 2 | $P_{sac7d}$ | 3 | 3 | 1.835 ± 0.405 | 742 | 6.91 | 592 |

| Plate | Promoter | B_rep | T_rep | Slope and error $\times 10^3$ | $t_{off}$ | m | d |
|---|---|---|---|---|---|---|---|
| 2 | $P_{malE\_pJL1601}$ | 1 | 1 | 0.420 ± 0.021 | 1499 | 1.19 | 1681 |
| 2 | $P_{malE\_pJL1601}$ | 1 | 2 | 0.355 ± 0.018 | 1106 | 1.49 | 1661 |
| 2 | $P_{malE\_pJL1601}$ | 1 | 3 | 0.361 ± 0.012 | 1198 | 1.36 | 1696 |
| 2 | $P_{malE\_pJL1601}$ | 2 | 1 | 0.318 ± 0.011 | 1122 | 1.14 | 1873 |
| 2 | $P_{malE\_pJL1601}$ | 2 | 2 | 0.283 ± 0.010 | 1098 | 1.35 | 1910 |
| 2 | $P_{malE\_pJL1601}$ | 2 | 3 | 0.292 ± 0.014 | 1210 | 1.35 | 1914 |
| 2 | $P_{malE\_pJL1601}$ | 3 | 1 | 0.363 ± 0.015 | 1365 | 1.30 | 1745 |
| 2 | $P_{malE\_pJL1601}$ | 3 | 2 | 0.350 ± 0.017 | 1257 | 1.39 | 1755 |
| 2 | $P_{malE\_pJL1601}$ | 3 | 3 | 0.377 ± 0.020 | 1337 | 1.21 | 1716 |

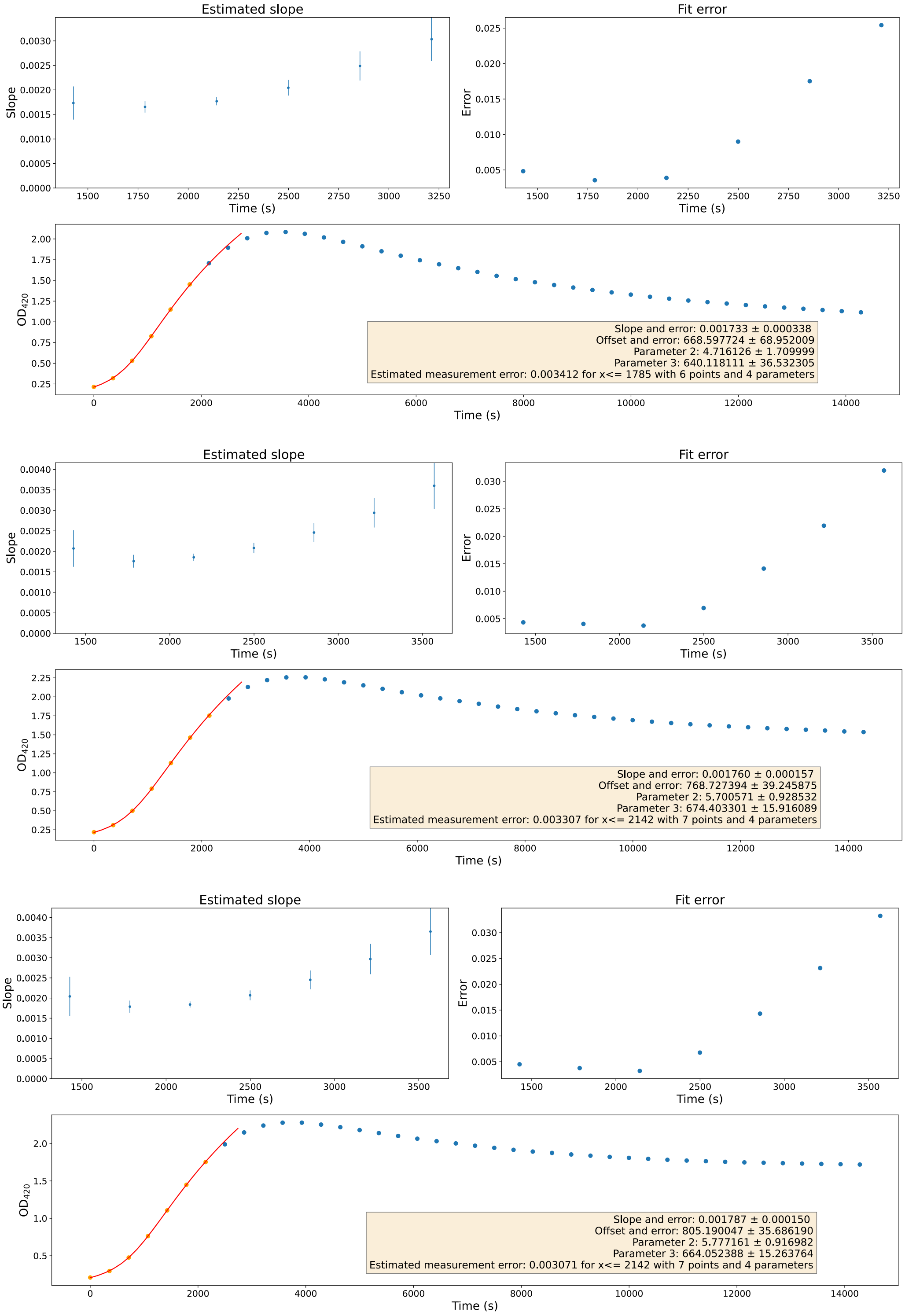


**Supplementary Figure 1: $P_{Ec10\_Sa1}$ Biological Replicate 1** - Estimated slopes (top left), fit errors (top right) and absorbance at 420 nm (bottom), for technical replicates 1 (top), 2 (middle) and 3 (bottom). Parameter 2: m (maximum value) and parameter 3: d (period around the offset in which the transition takes place).

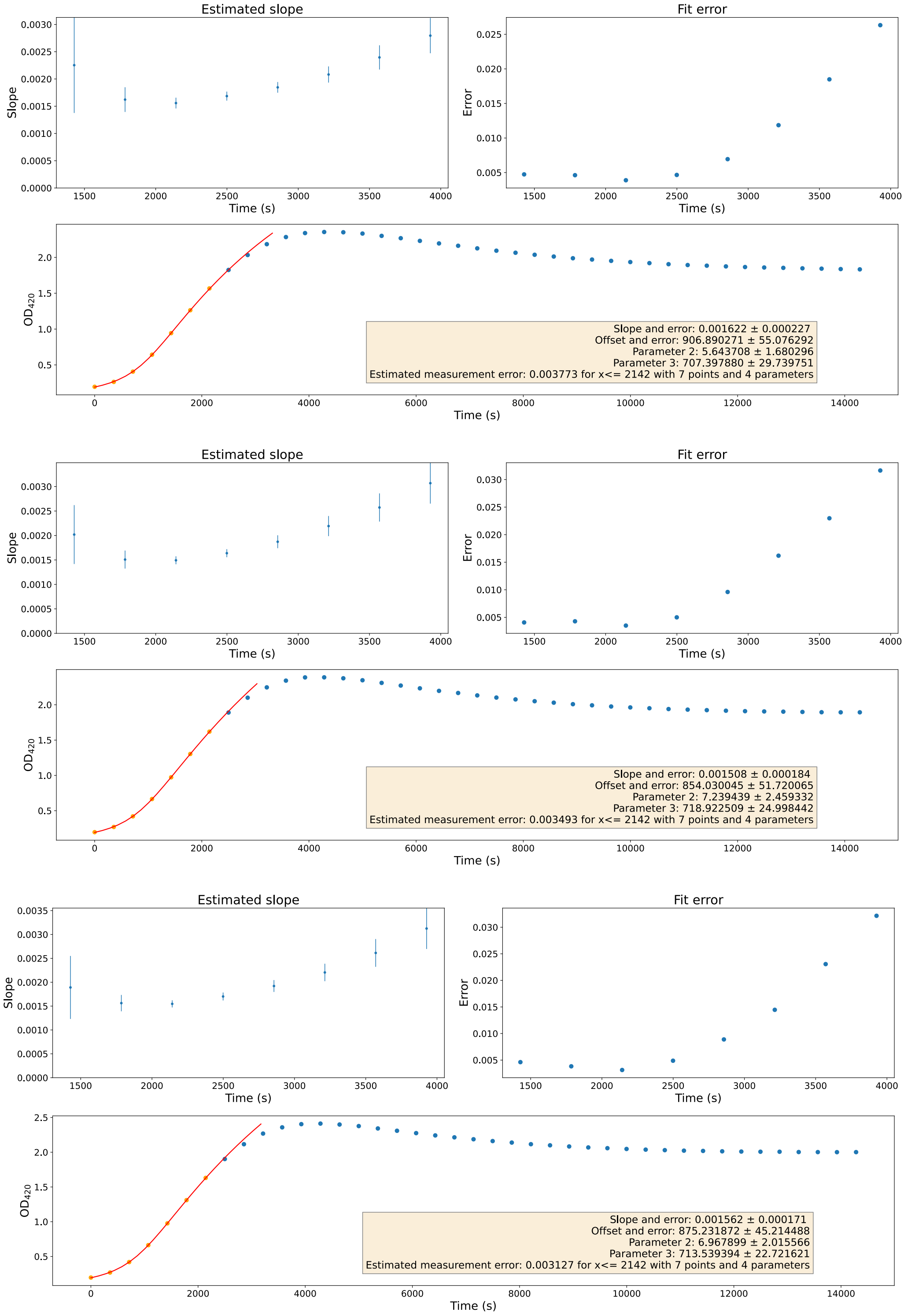


**Supplementary Figure 2: $\mathbf{P_{Ec10\_Sa1}}$ Biological Replicate 2** - Estimated slopes (top left), fit errors (top right) and absorbance at 420 nm (bottom), for technical replicates 1 (top), 2 (middle) and 3 (bottom). Parameter 2: m (maximum value) and parameter 3: d (period around the offset in which the transition takes place).

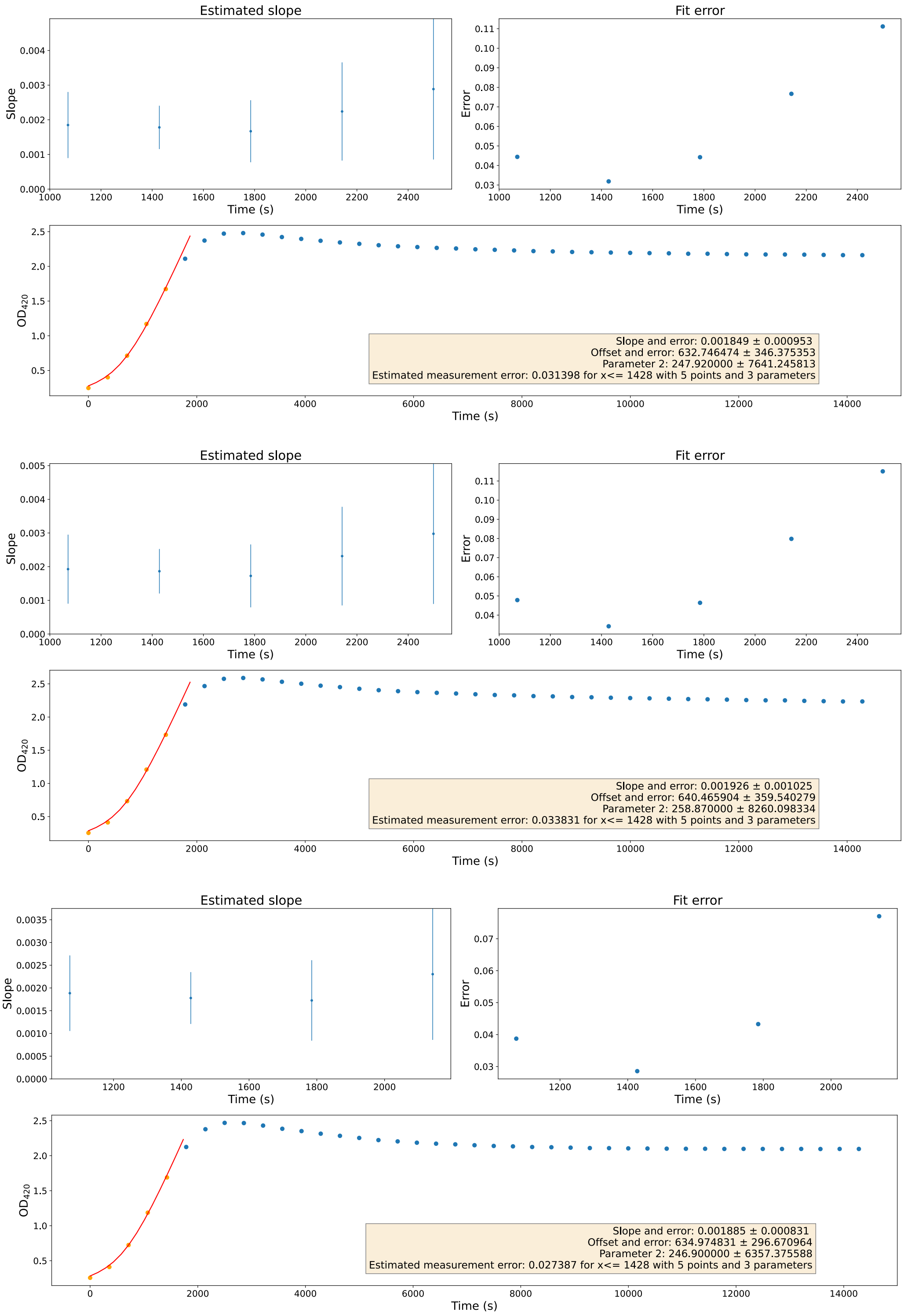


**Supplementary Figure 3: $\mathbf{P_{Ec10\_Sa1}}$ Biological Replicate 3** - Estimated slopes (top left), fit errors (top right) and absorbance at 420 nm (bottom), for technical replicates 1 (top), 2 (middle) and 3 (bottom). Parameter 2: m (maximum value) and parameter 3: d (period around the offset in which the transition takes place).

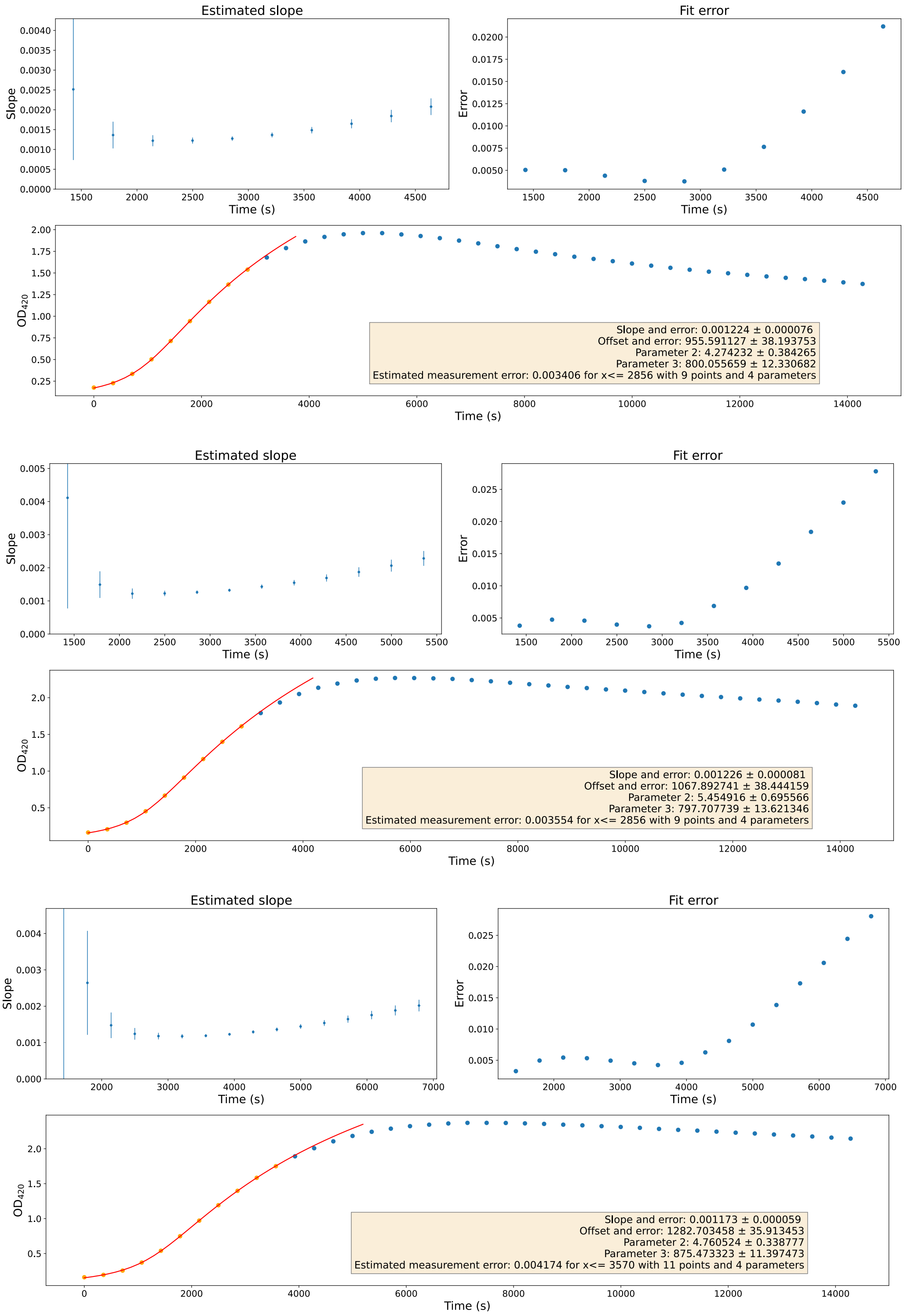


**Supplementary Figure 4: $P_{Ec6\_Sa2}$ Biological Replicate 1** - Estimated slopes (top left), fit errors (top right) and absorbance at 420 nm (bottom), for technical replicates 1 (top), 2 (middle) and 3 (bottom). Parameter 2: m (maximum value) and parameter 3: d (period around the offset in which the transition takes place).

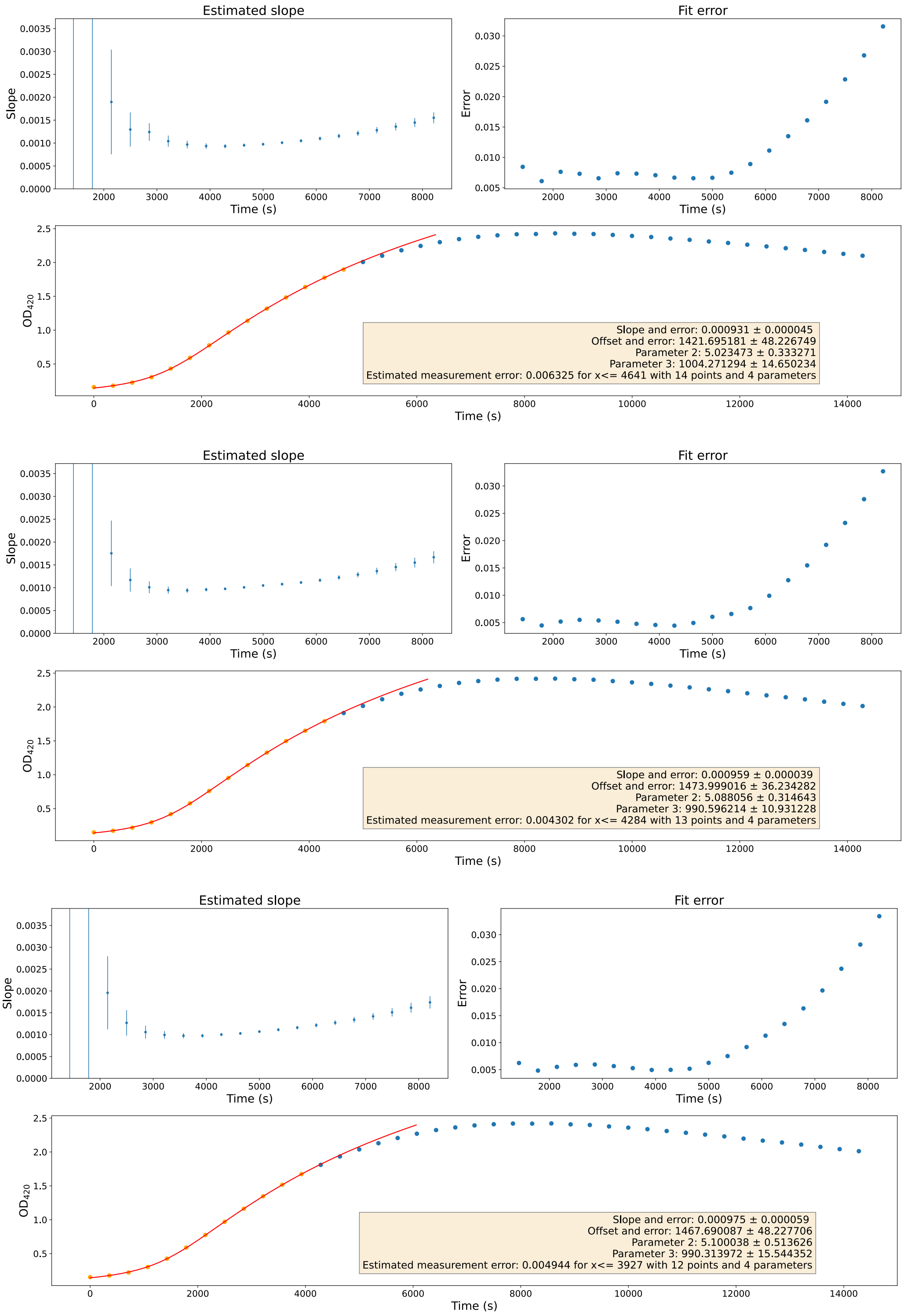


**Supplementary Figure 5: $P_{Ec6\_Sa2}$ Biological Replicate 2** - Estimated slopes (top left), fit errors (top right) and absorbance at 420 nm (bottom), for technical replicates 1 (top), 2 (middle) and 3 (bottom). Parameter 2: m (maximum value) and parameter 3: d (period around the offset in which the transition takes place).

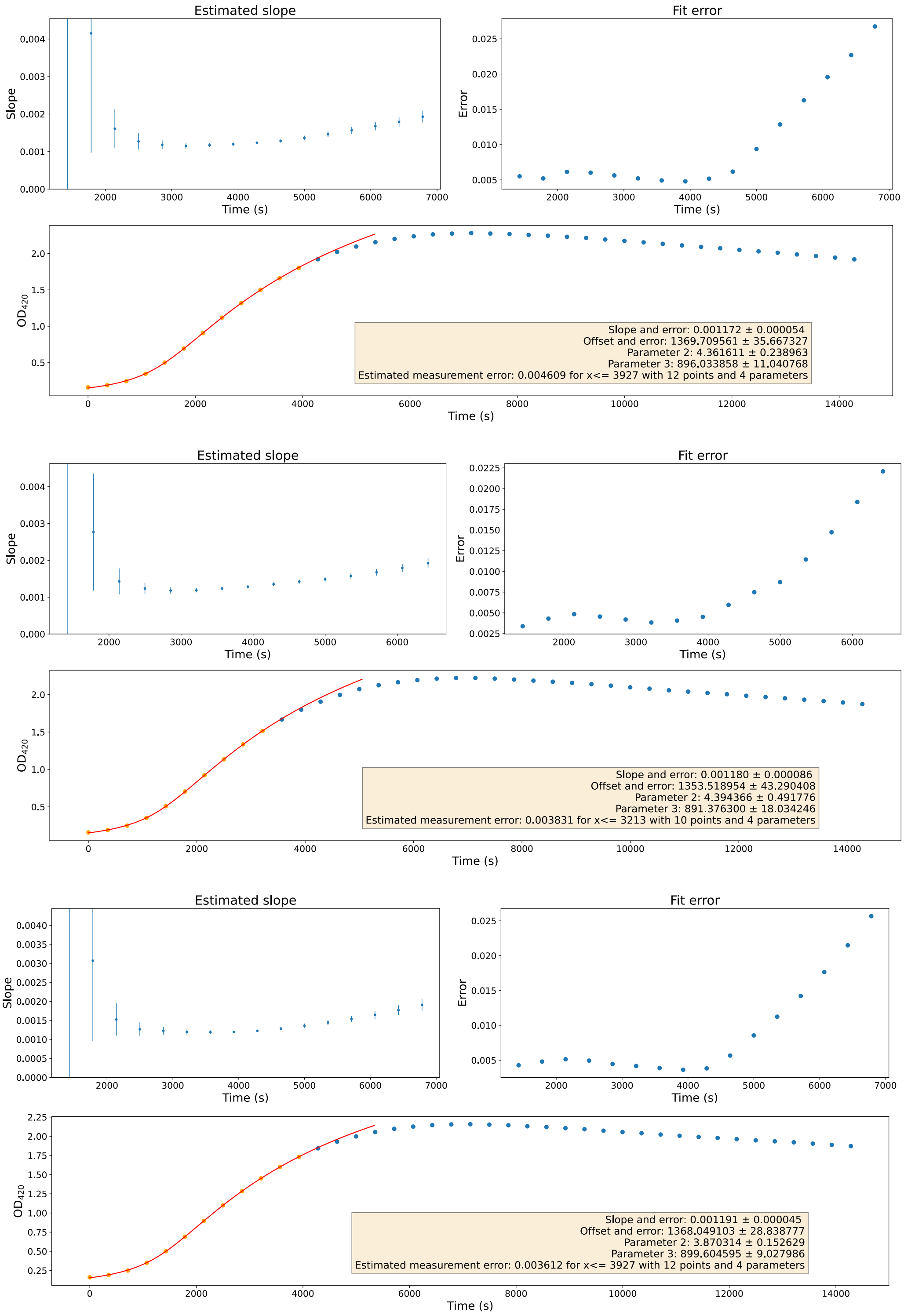


**Supplementary Figure 6: $\mathbf{P_{Ec6\_Sa2}}$ Biological Replicate 3** - Estimated slopes (top left), fit errors (top right) and absorbance at 420 nm (bottom), for technical replicates 1 (top), 2 (middle) and 3 (bottom). Parameter 2: m (maximum value) and parameter 3: d (period around the offset in which the transition takes place).

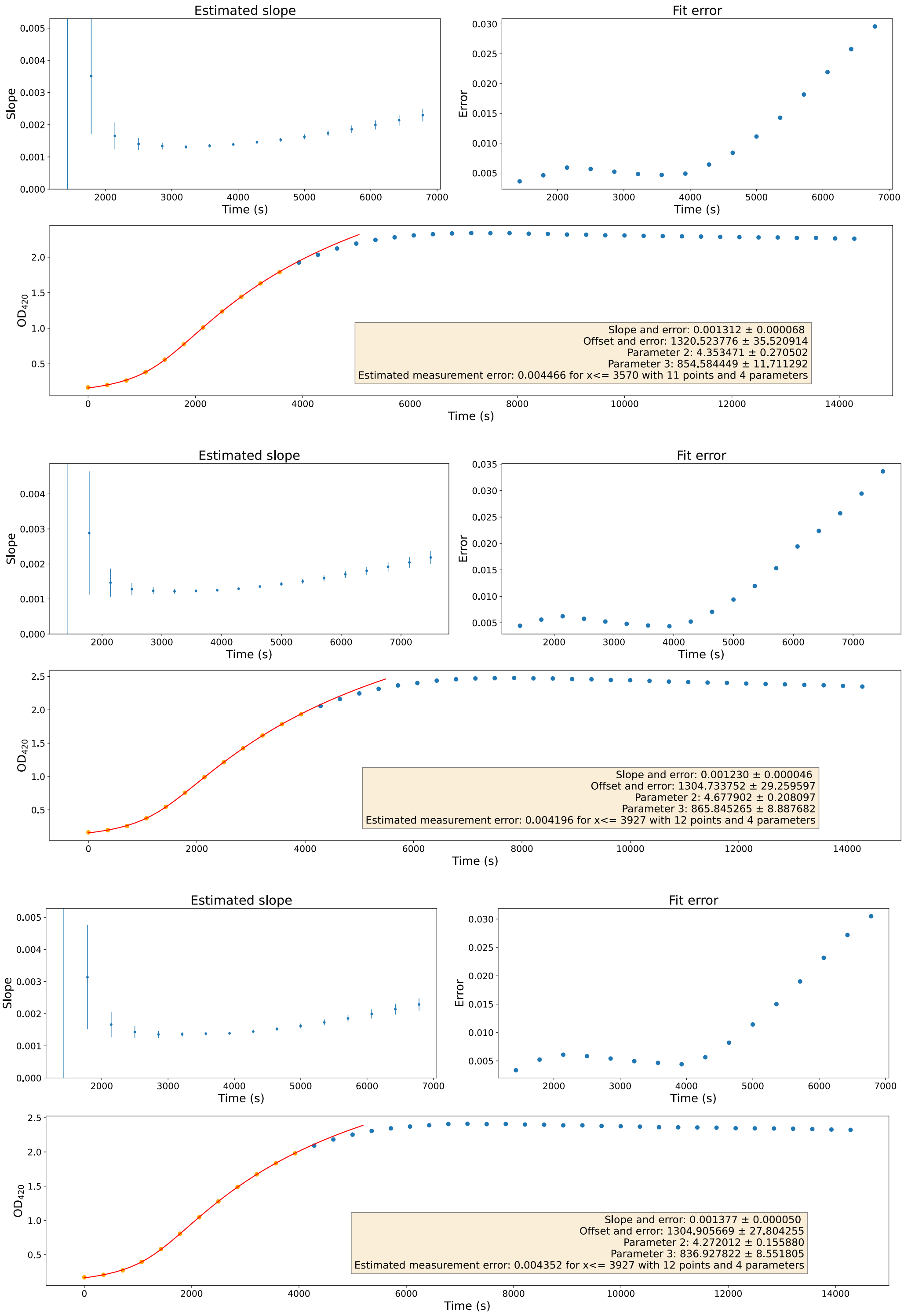


**Supplementary Figure 7: $P_{Ec7\_Sa3}$ Biological Replicate 1** - Estimated slopes (top left), fit errors (top right) and absorbance at 420 nm (bottom), for technical replicates 1 (top), 2 (middle) and 3 (bottom). Parameter 2: m (maximum value) and parameter 3: d (period around the offset in which the transition takes place).

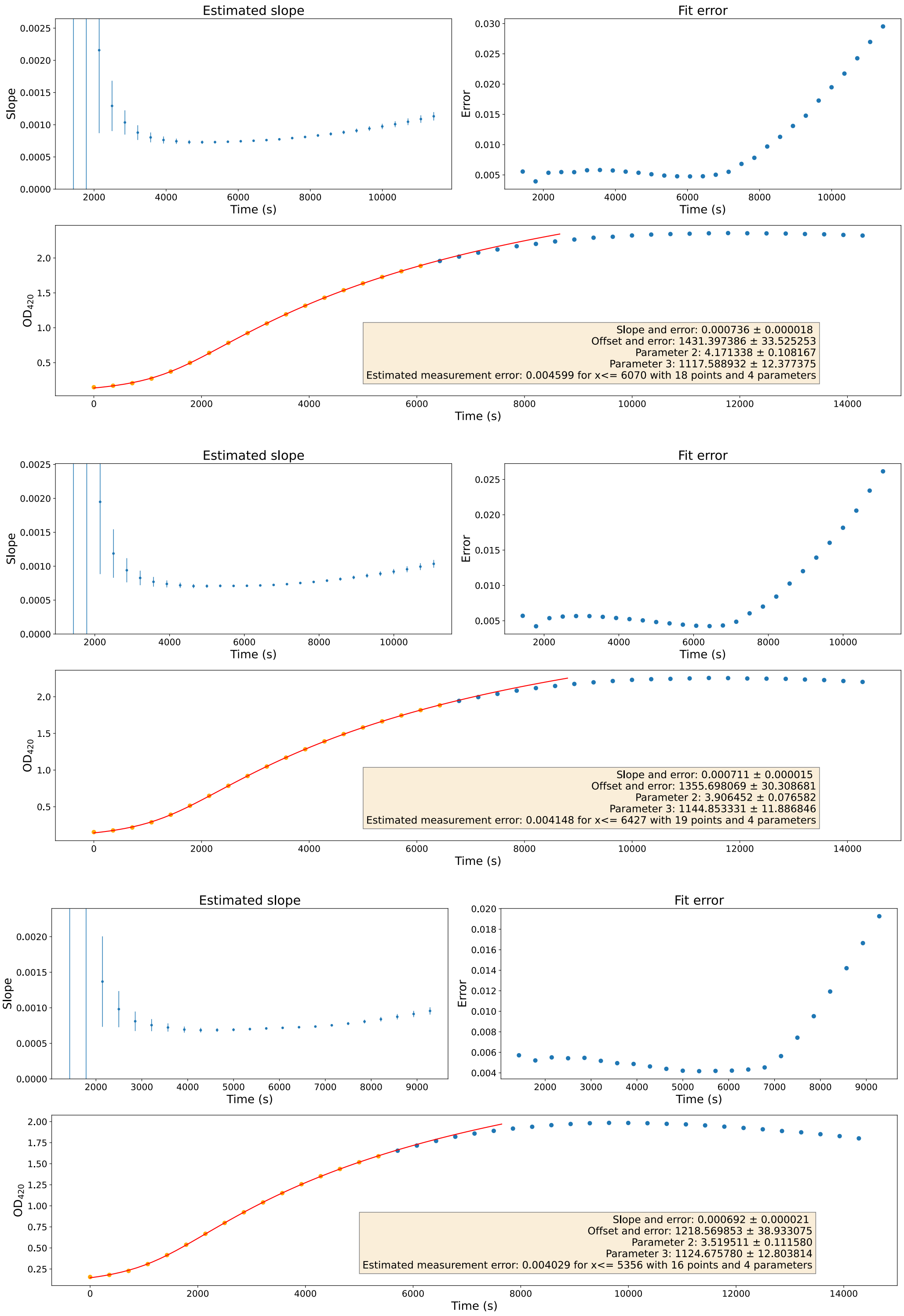


**Supplementary Figure 8: $\mathbf{P_{Ec7\_Sa3}}$ Biological Replicate 2** - Estimated slopes (top left), fit errors (top right) and absorbance at 420 nm (bottom), for technical replicates 1 (top), 2 (middle) and 3 (bottom). Parameter 2: m (maximum value) and parameter 3: d (period around the offset in which the transition takes place).

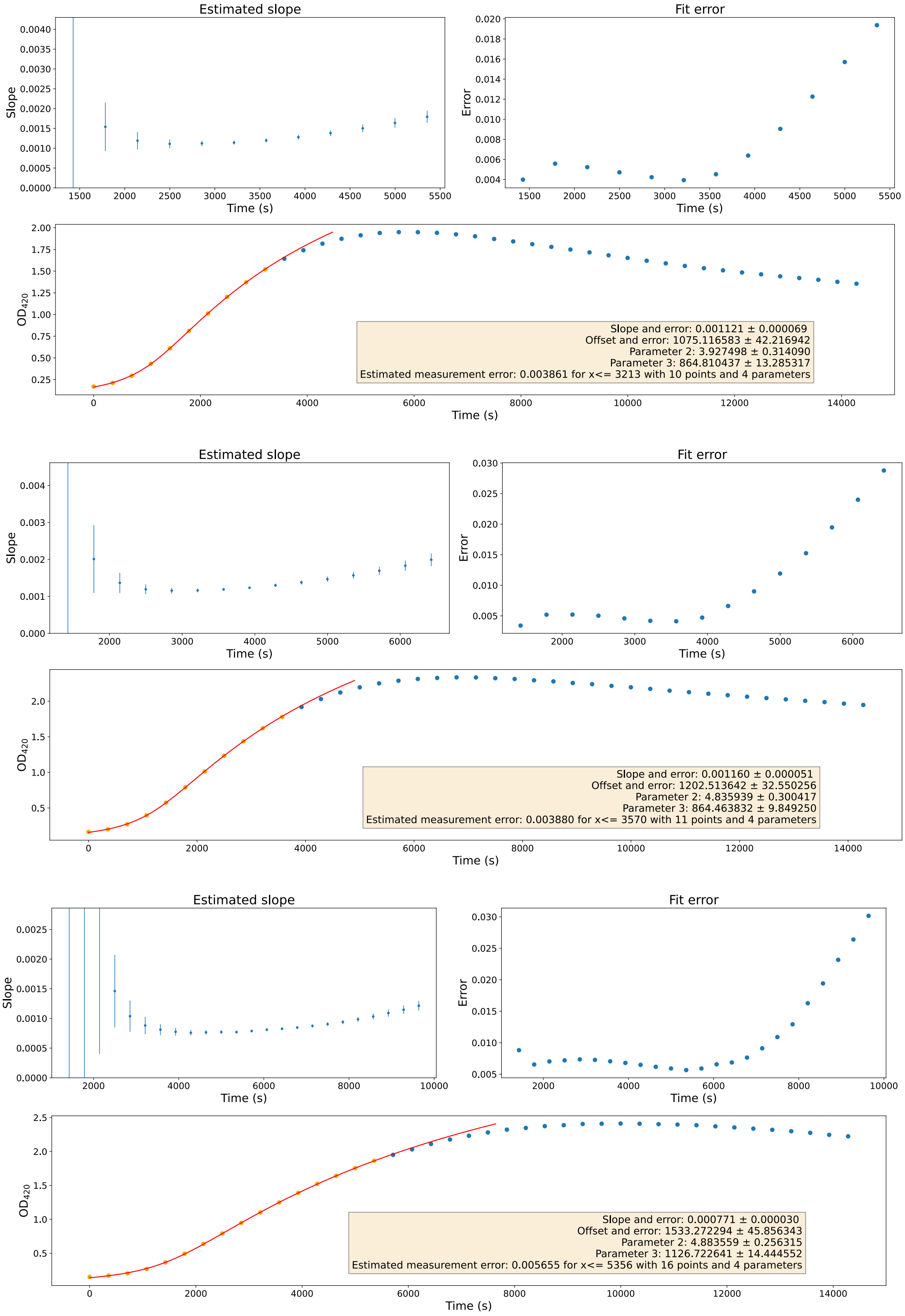


**Supplementary Figure 9: $P_{Ec7\_Sa3}$ Biological Replicate 3** - Estimated slopes (top left), fit errors (top right) and absorbance at 420 nm (bottom), for technical replicates 1 (top), 2 (middle) and 3 (bottom). Parameter 2: m (maximum value) and parameter 3: d (period around the offset in which the transition takes place).

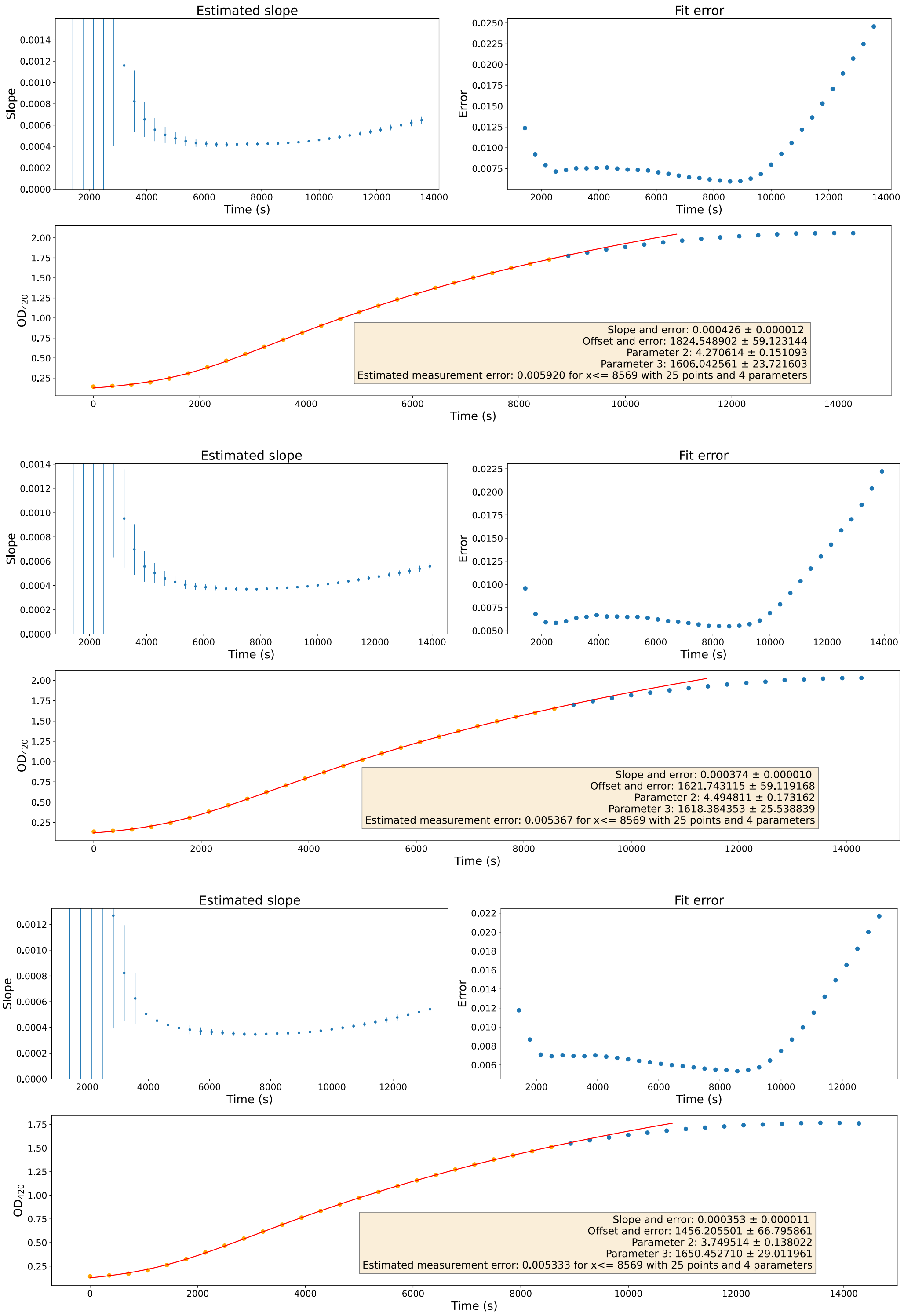


**Supplementary Figure 10: $\mathbf{P_{Ec8\_Sa4}}$ Biological Replicate 1** - Estimated slopes (top left), fit errors (top right) and absorbance at 420 nm (bottom), for technical replicates 1 (top), 2 (middle) and 3 (bottom). Parameter 2: m (maximum value) and parameter 3: d (period around the offset in which the transition takes place).

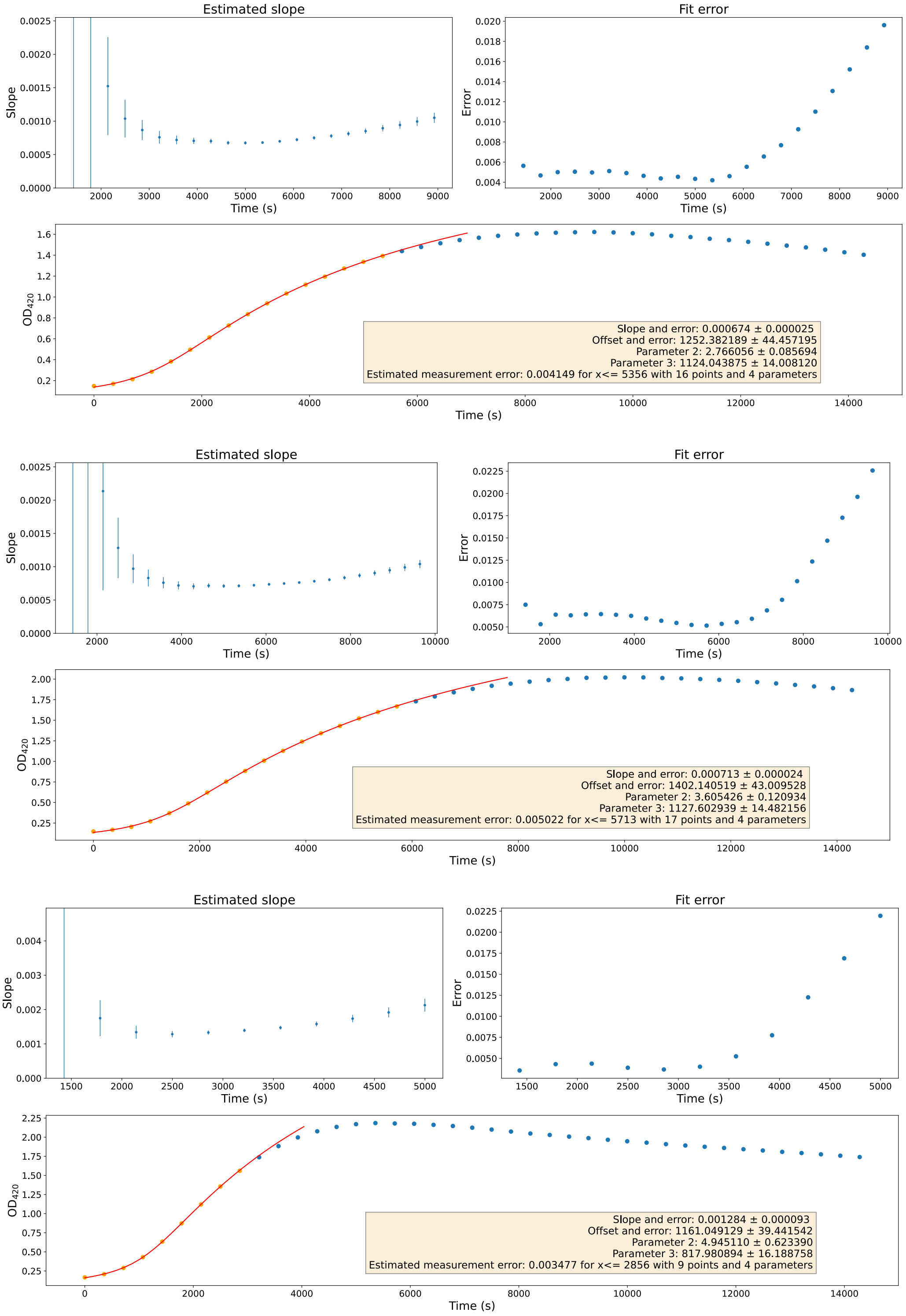


**Supplementary Figure 11: $\mathbf{P_{Ec8\_Sa4}}$ Biological Replicate 2** - Estimated slopes (top left), fit errors (top right) and absorbance at 420 nm (bottom), for technical replicates 1 (top), 2 (middle) and 3 (bottom). Parameter 2: m (maximum value) and parameter 3: d (period around the offset in which the transition takes place).

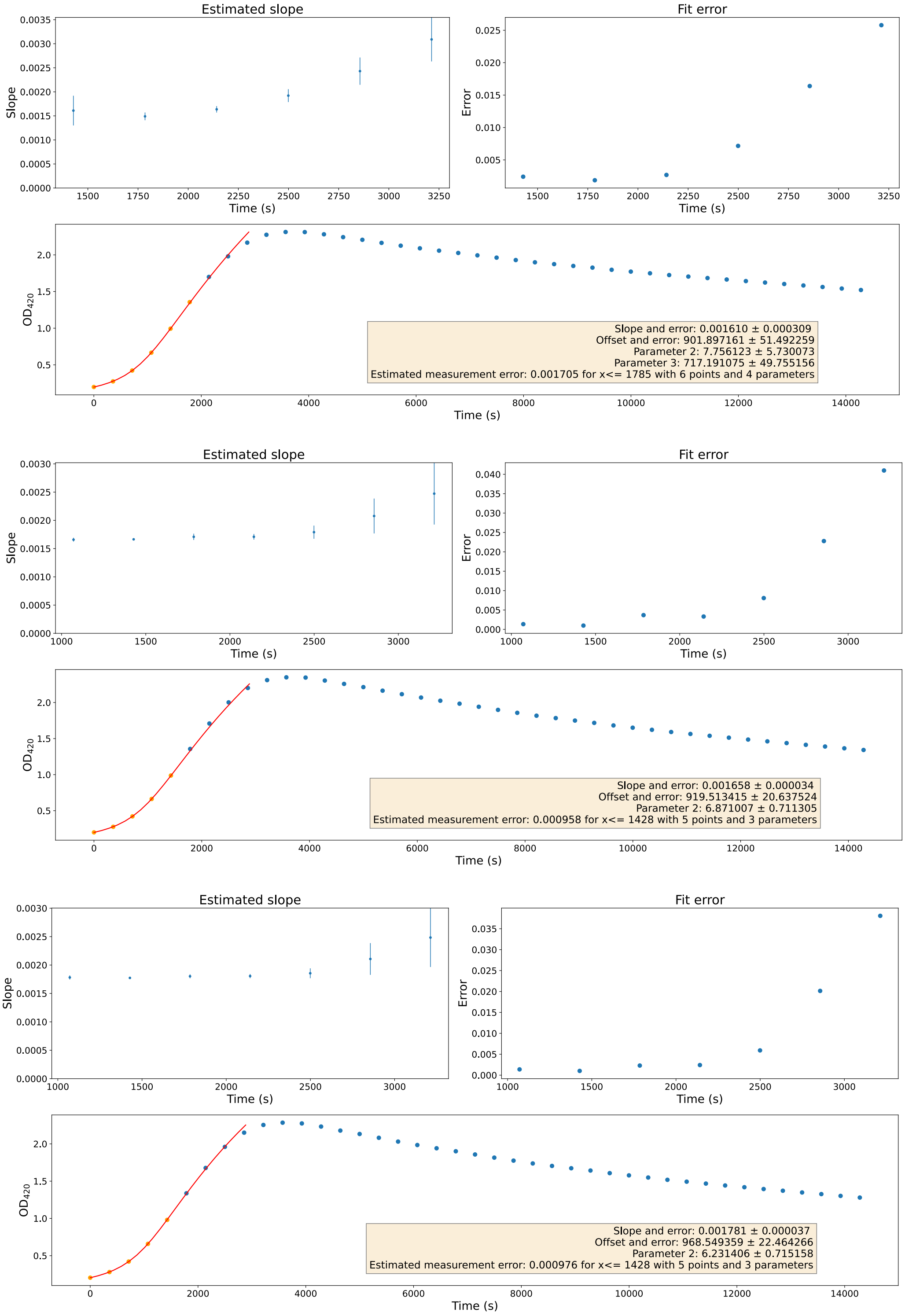


**Supplementary Figure 12: $\mathbf{P_{Ec8\_Sa4}}$ Biological Replicate 3** - Estimated slopes (top left), fit errors (top right) and absorbance at 420 nm (bottom), for technical replicates 1 (top), 2 (middle) and 3 (bottom). Parameter 2: m (maximum value) and parameter 3: d (period around the offset in which the transition takes place).

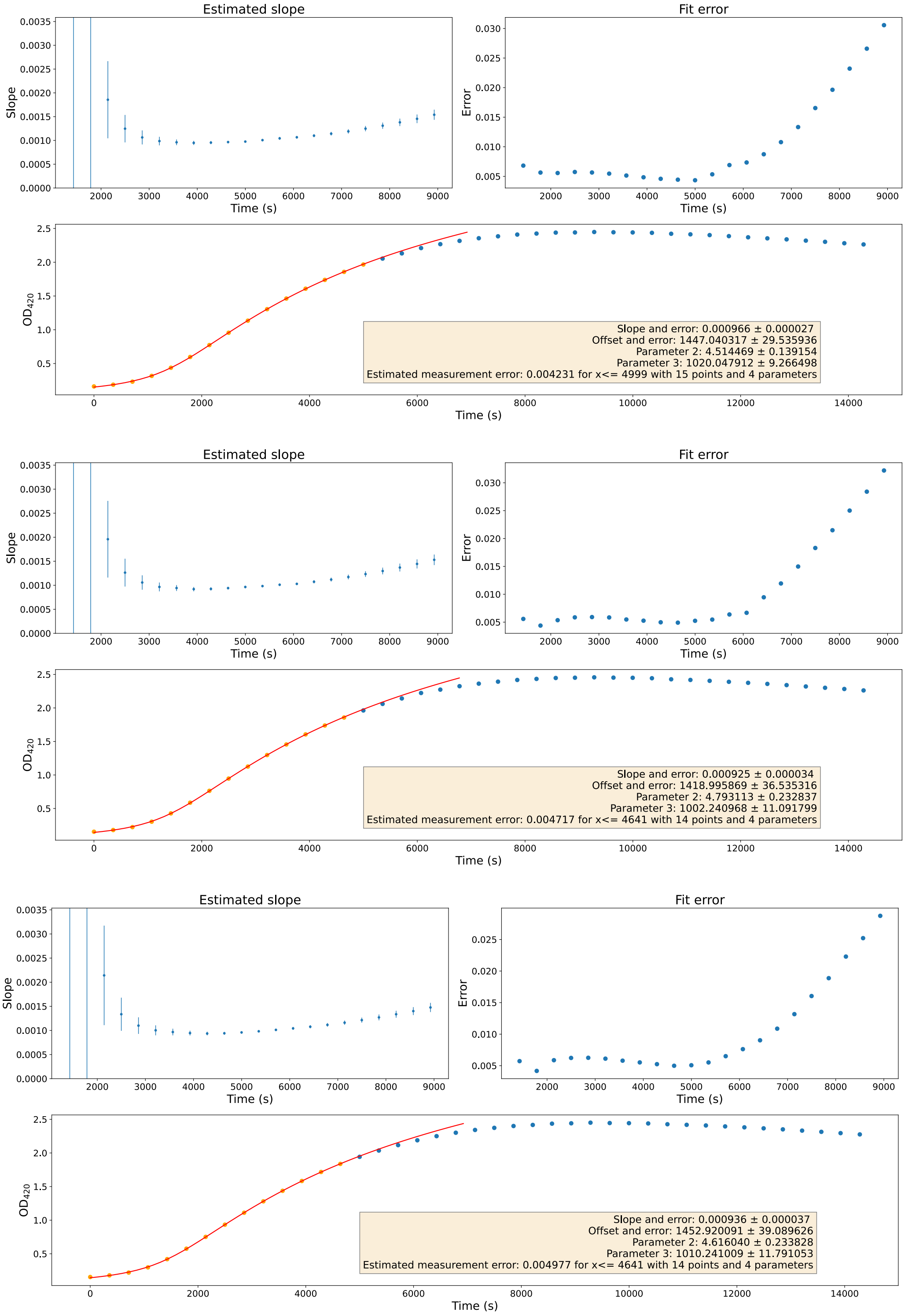


**Supplementary Figure 13: $P_{Ec4\_Sa5}$ Biological Replicate 1** - Estimated slopes (top left), fit errors (top right) and absorbance at 420 nm (bottom), for technical replicates 1 (top), 2 (middle) and 3 (bottom). Parameter 2: m (maximum value) and parameter 3: d (period around the offset in which the transition takes place).

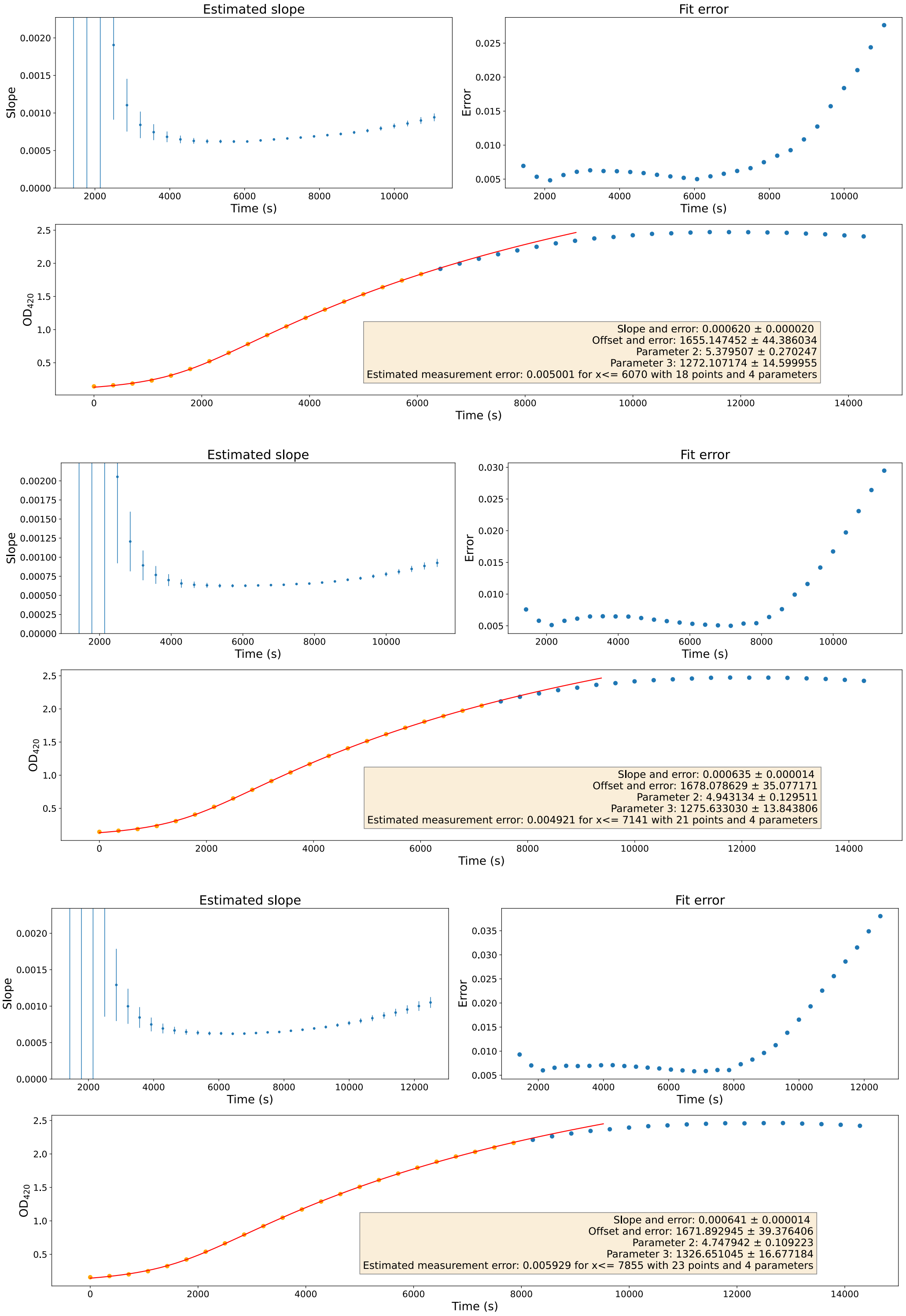


**Supplementary Figure 14: $P_{Ec4\_Sa5}$ Biological Replicate 2** - Estimated slopes (top left), fit errors (top right) and absorbance at 420 nm (bottom), for technical replicates 1 (top), 2 (middle) and 3 (bottom). Parameter 2: m (maximum value) and parameter 3: d (period around the offset in which the transition takes place).

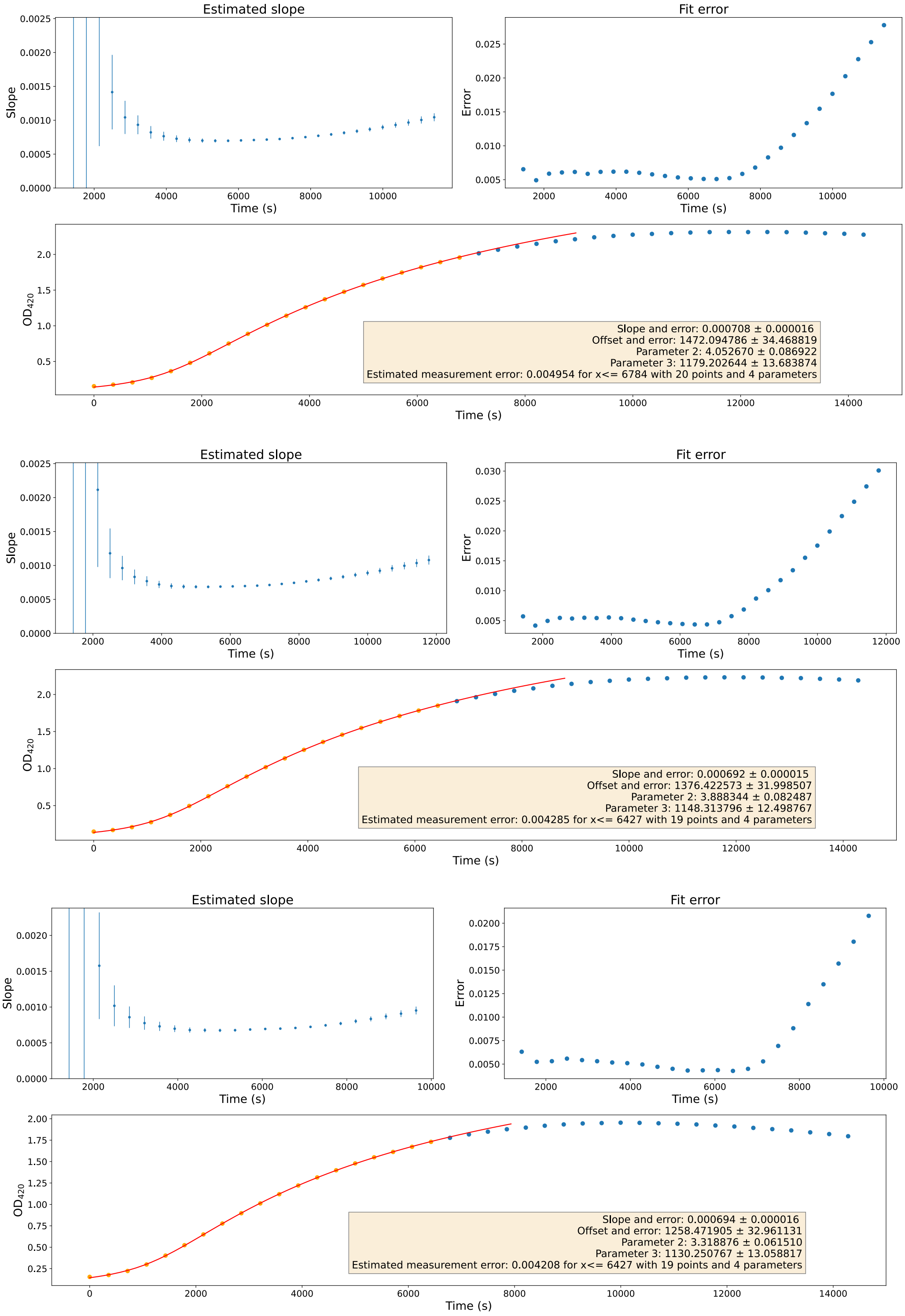


**Supplementary Figure 15: $P_{Ec4\_Sa5}$ Biological Replicate 3** - Estimated slopes (top left), fit errors (top right) and absorbance at 420 nm (bottom), for technical replicates 1 (top), 2 (middle) and 3 (bottom). Parameter 2: m (maximum value) and parameter 3: d (period around the offset in which the transition takes place).

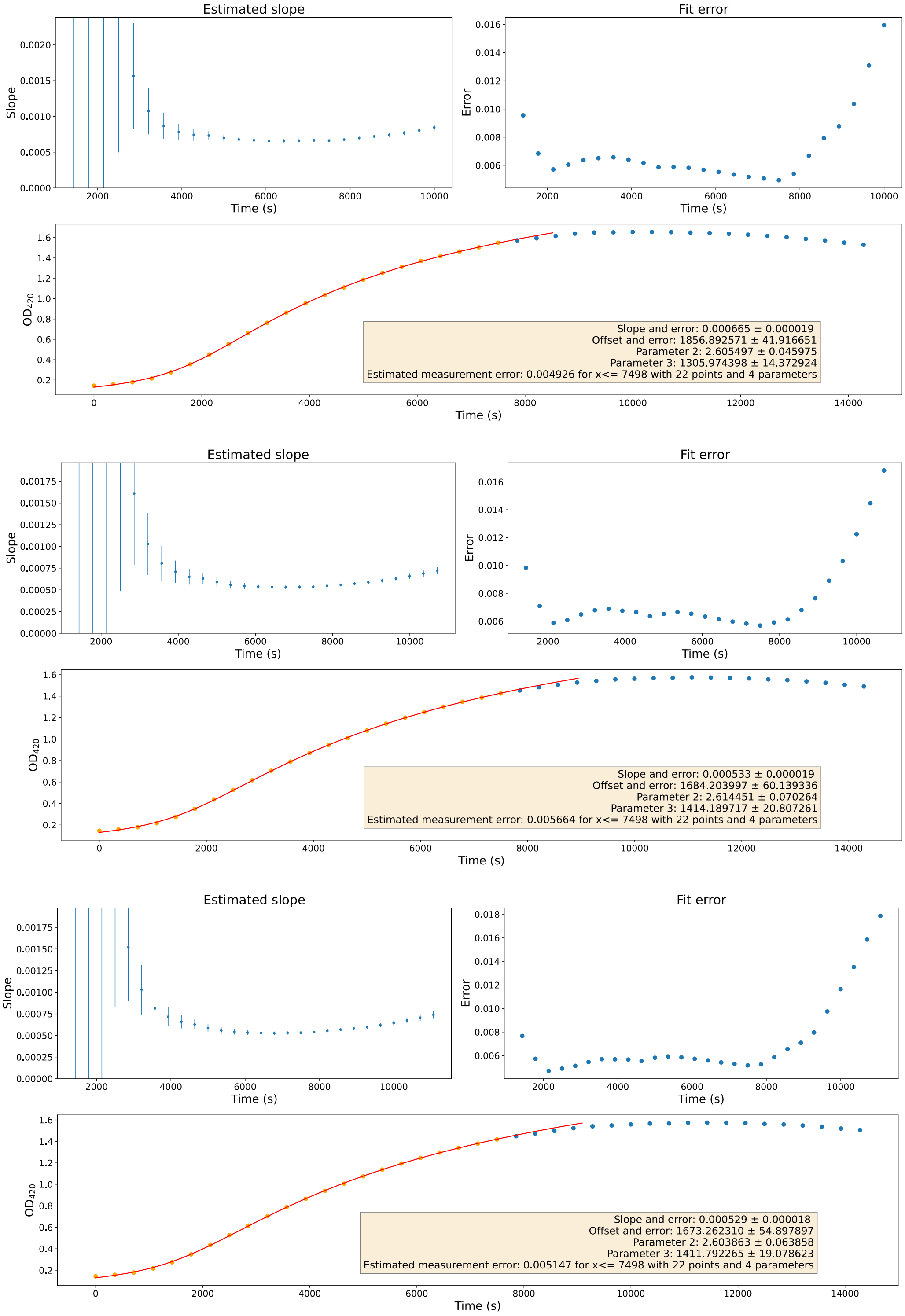


**Supplementary Figure 16: $P_{Ec9\_Sa6}$ Biological Replicate 1** - Estimated slopes (top left), fit errors (top right) and absorbance at 420 nm (bottom), for technical replicates 1 (top), 2 (middle) and 3 (bottom). Parameter 2: m (maximum value) and parameter 3: d (period around the offset in which the transition takes place).

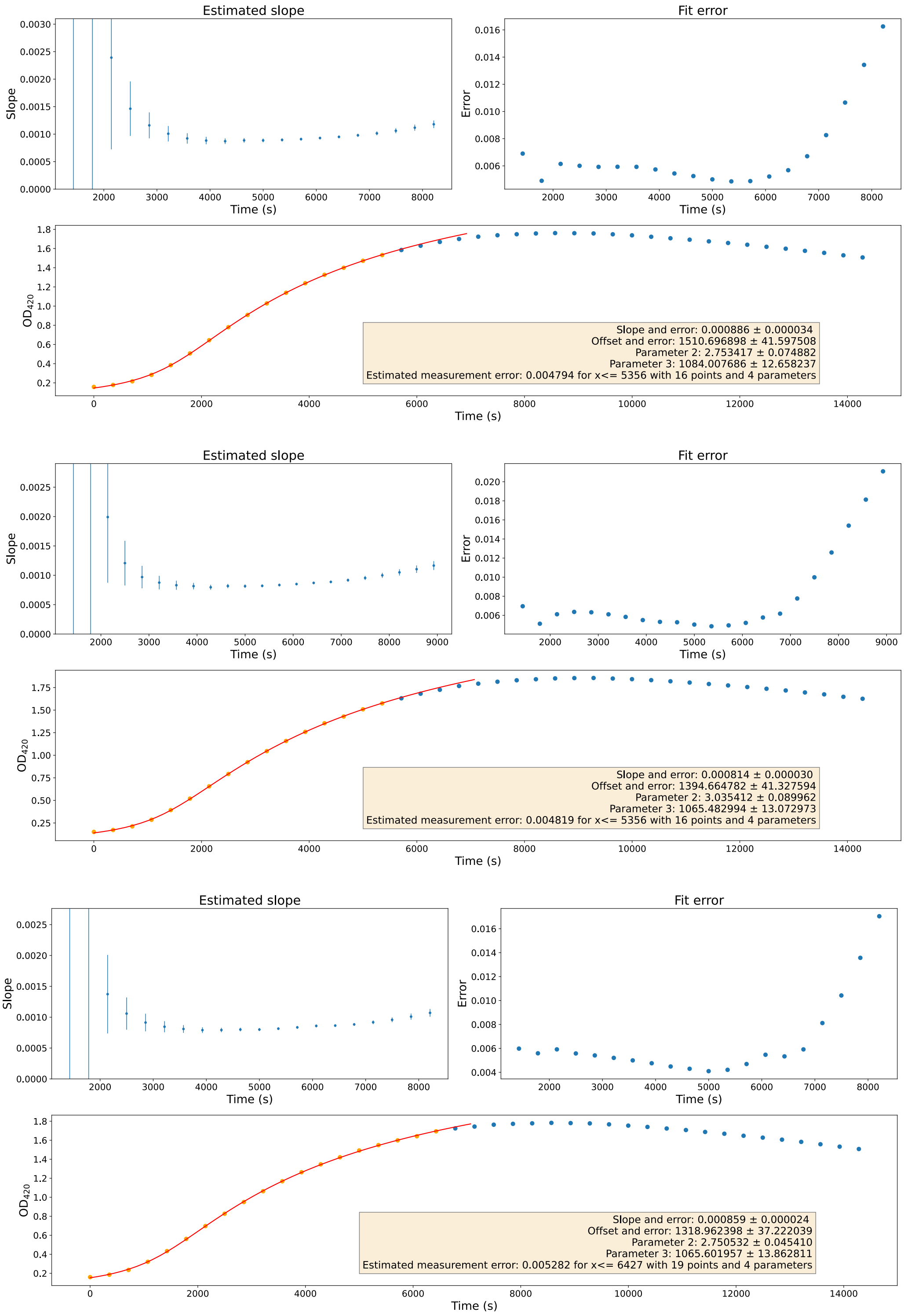


**Supplementary Figure 17: $P_{Ec9\_Sa6}$ Biological Replicate 2** - Estimated slopes (top left), fit errors (top right) and absorbance at 420 nm (bottom), for technical replicates 1 (top), 2 (middle) and 3 (bottom). Parameter 2: m (maximum value) and parameter 3: d (period around the offset in which the transition takes place).

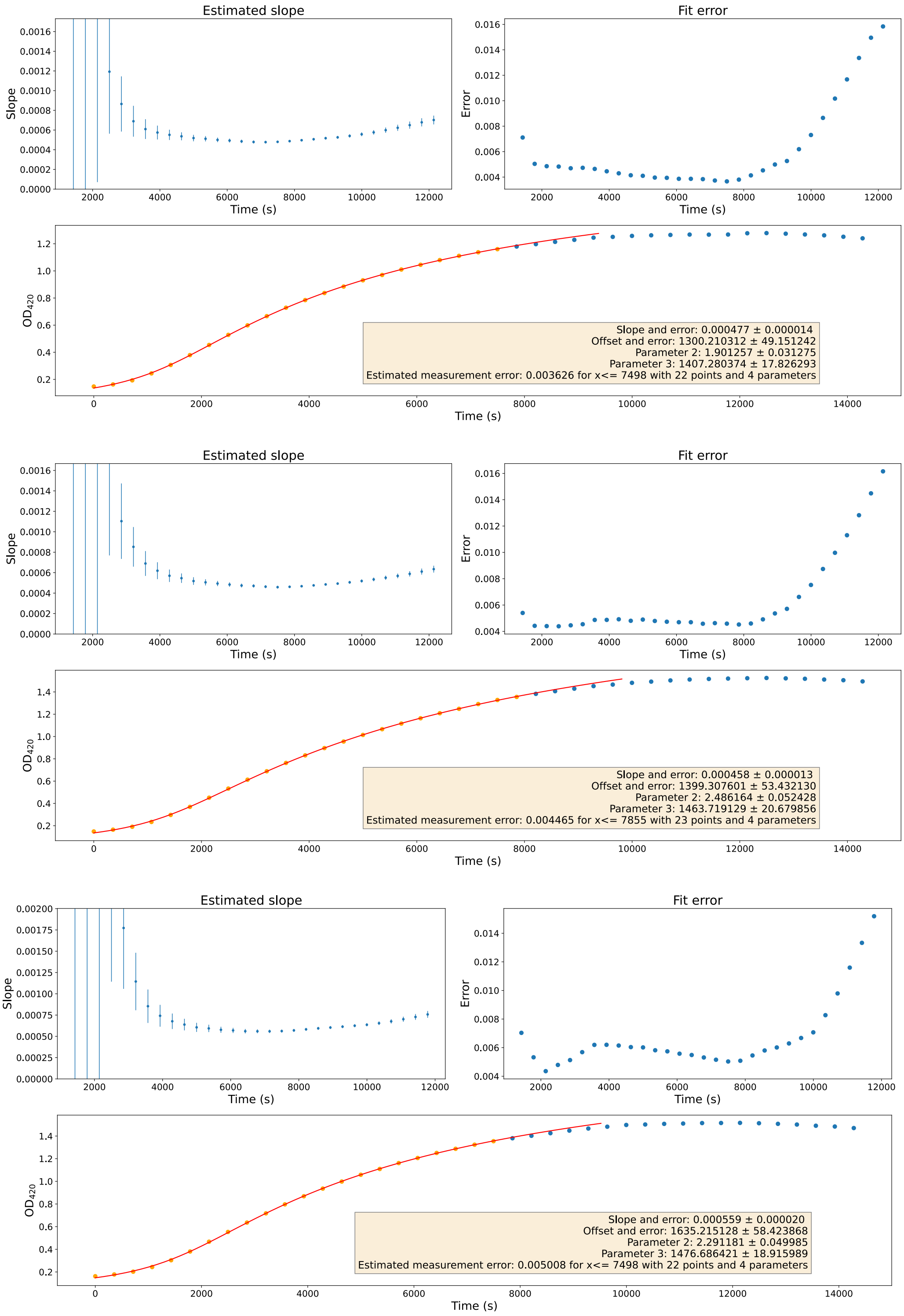


**Supplementary Figure 18: $P_{Ec9\_Sa6}$ Biological Replicate 3** - Estimated slopes (top left), fit errors (top right) and absorbance at 420 nm (bottom), for technical replicates 1 (top), 2 (middle) and 3 (bottom). Parameter 2: m (maximum value) and parameter 3: d (period around the offset in which the transition takes place).

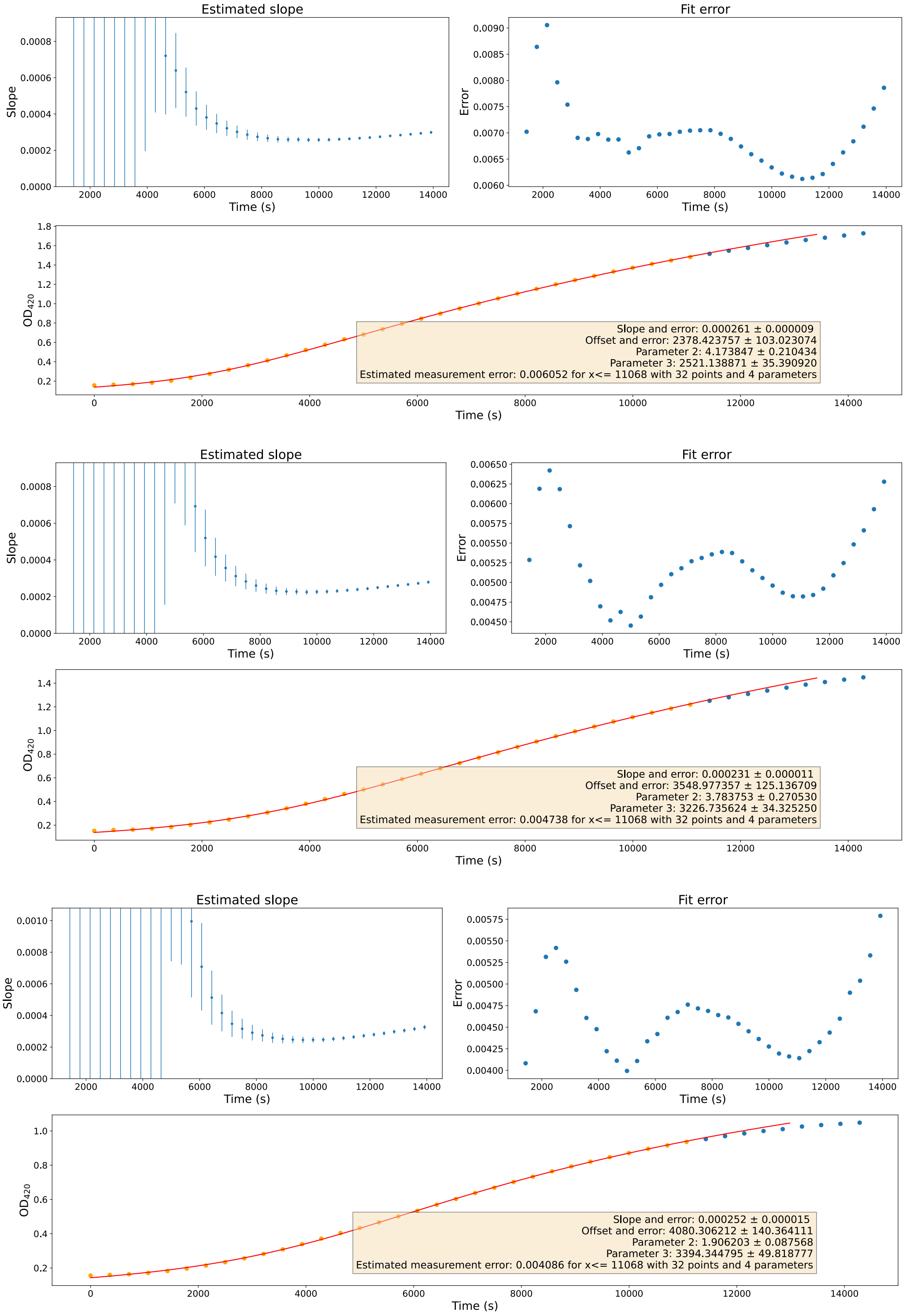


**Supplementary Figure 19: $\mathbf{P_{Ec3\_Sa7}}$ Biological Replicate 1** - Estimated slopes (top left), fit errors (top right) and absorbance at 420 nm (bottom), for technical replicates 1 (top), 2 (middle) and 3 (bottom). Parameter 2: m (maximum value) and parameter 3: d (period around the offset in which the transition takes place).

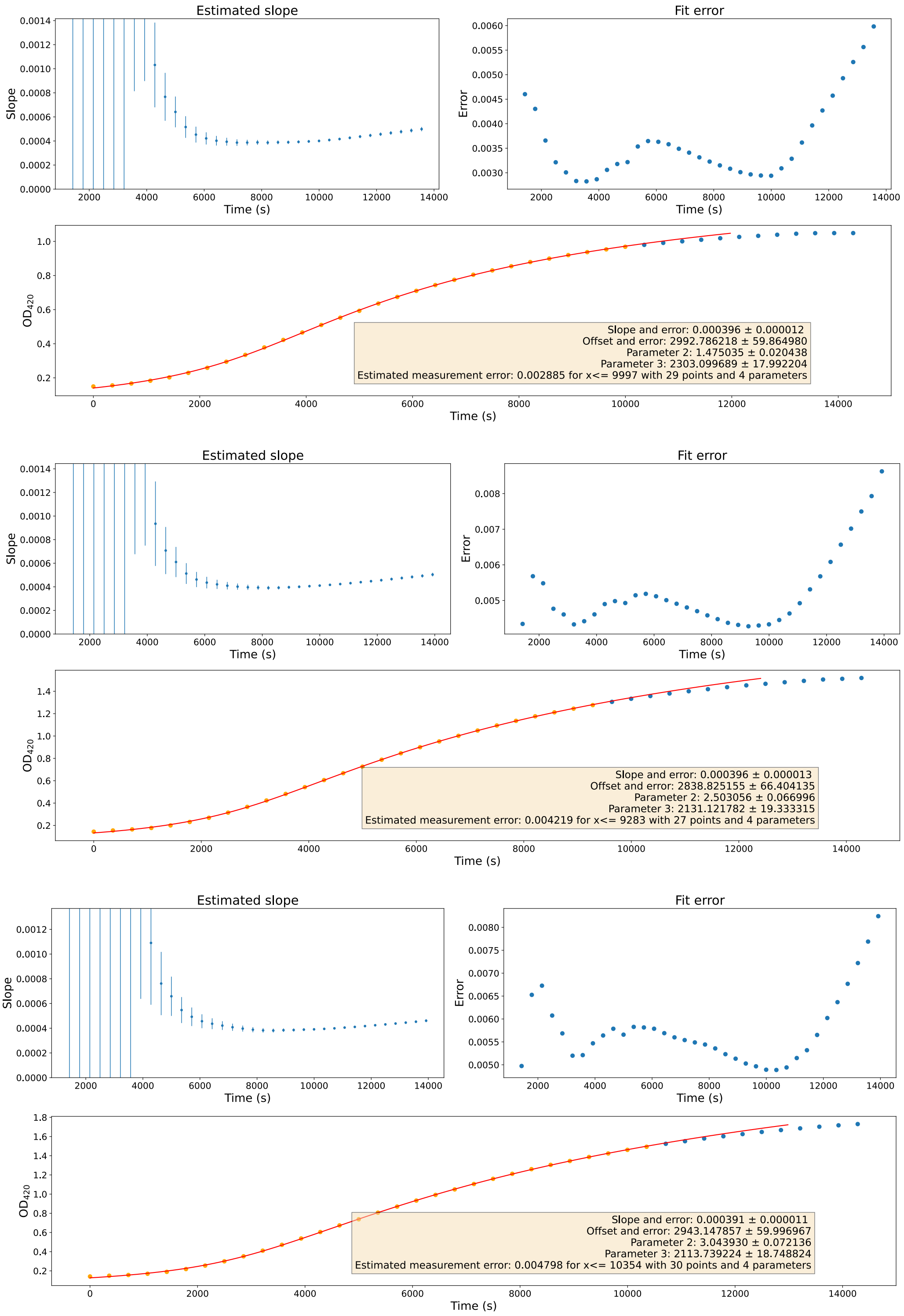


**Supplementary Figure 20: $\mathbf{P_{Ec3\_Sa7}}$ Biological Replicate 2** - Estimated slopes (top left), fit errors (top right) and absorbance at 420 nm (bottom), for technical replicates 1 (top), 2 (middle) and 3 (bottom). Parameter 2: m (maximum value) and parameter 3: d (period around the offset in which the transition takes place).

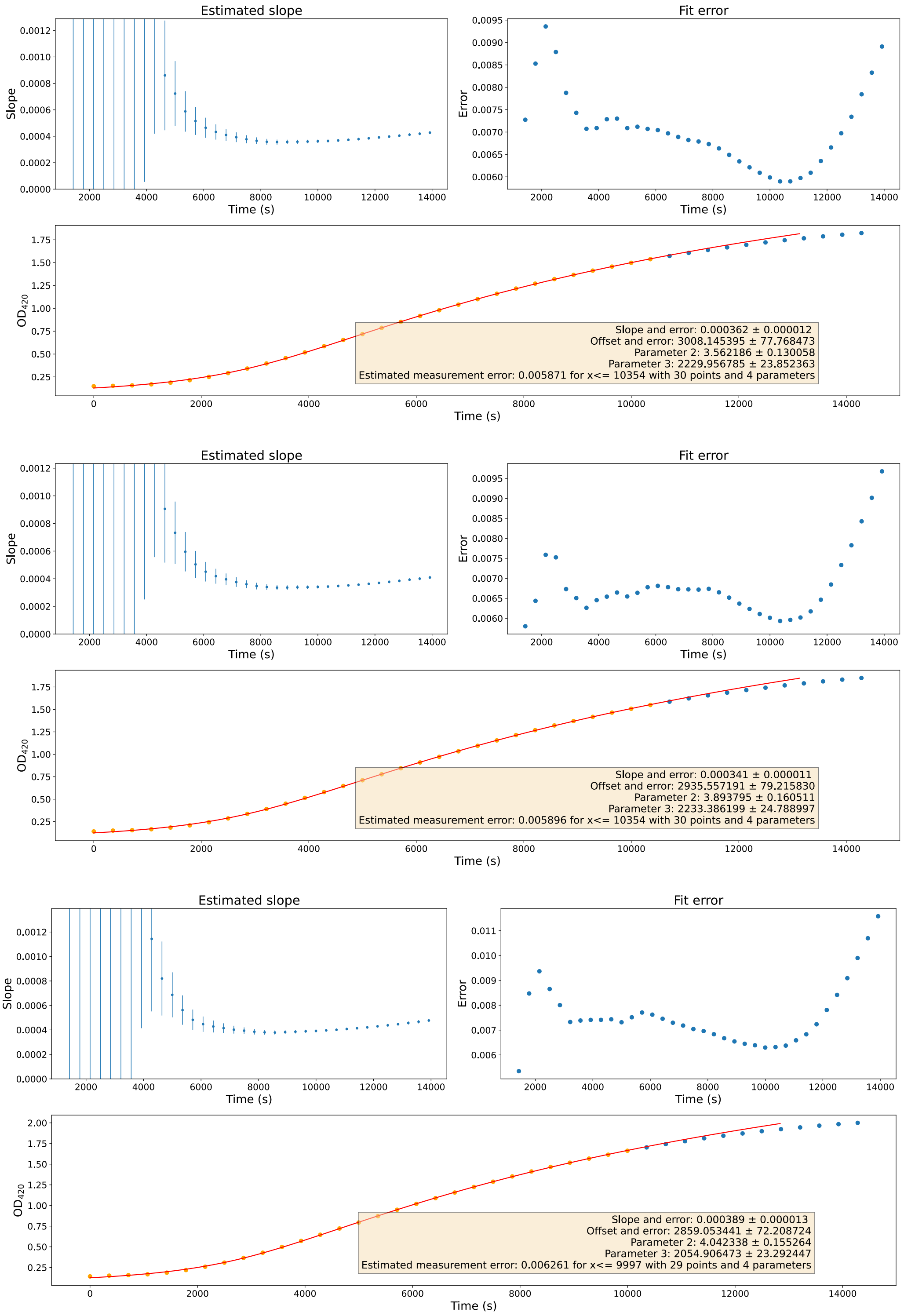


**Supplementary Figure 21: $\mathbf{P_{Ec3\_Sa7}}$ Biological Replicate 3** - Estimated slopes (top left), fit errors (top right) and absorbance at 420 nm (bottom), for technical replicates 1 (top), 2 (middle) and 3 (bottom). Parameter 2: m (maximum value) and parameter 3: d (period around the offset in which the transition takes place).

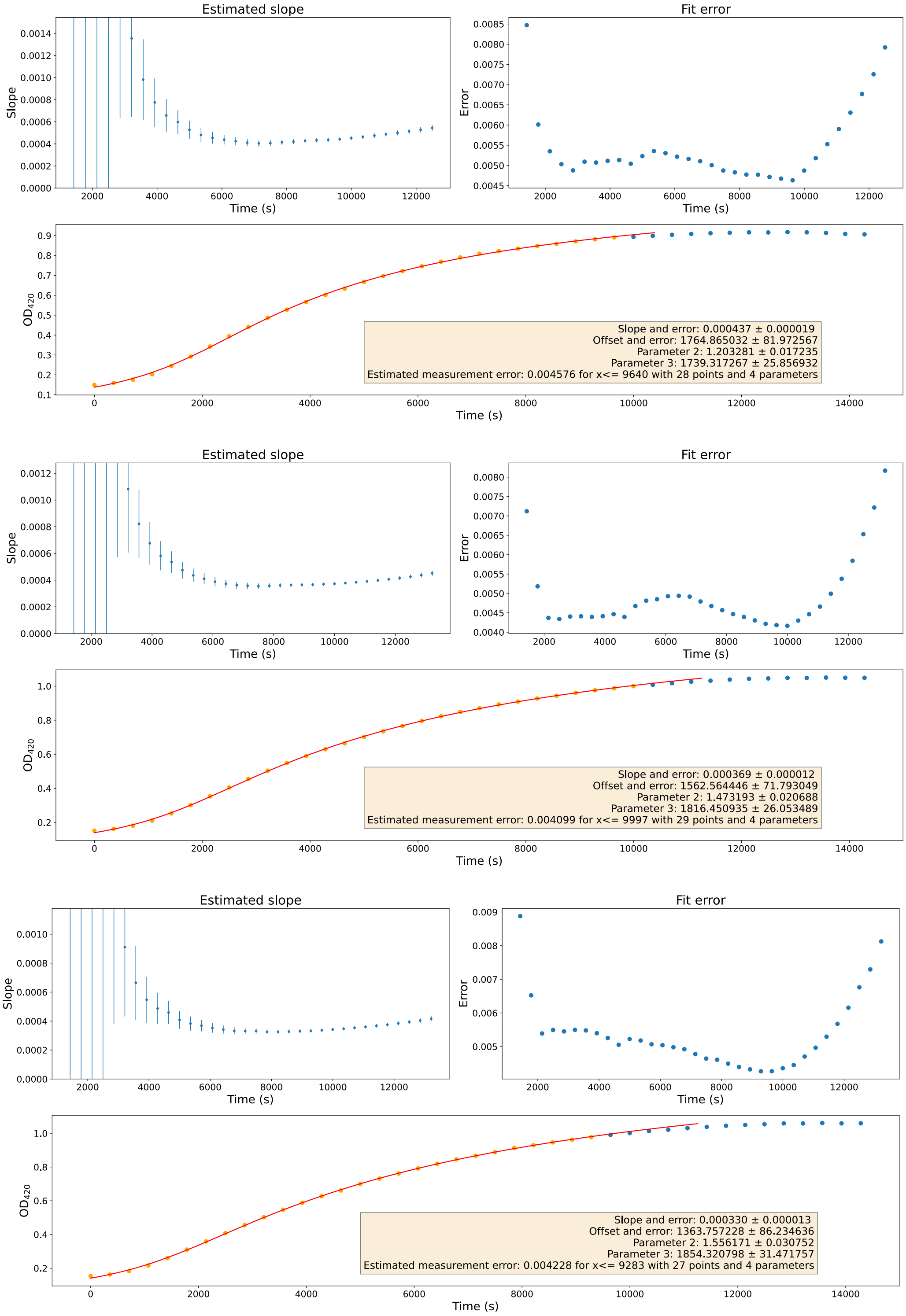


**Supplementary Figure 22: P$_{\mathbf{malE_pJL1601}}$ (Plate 1) Biological Replicate 1** - Estimated slopes (top left), fit errors (top right) and absorbance at 420 nm (bottom), for technical replicates 1 (top), 2 (middle) and 3 (bottom). Parameter 2: m (maximum value) and parameter 3: d (period around the offset in which the transition takes place).

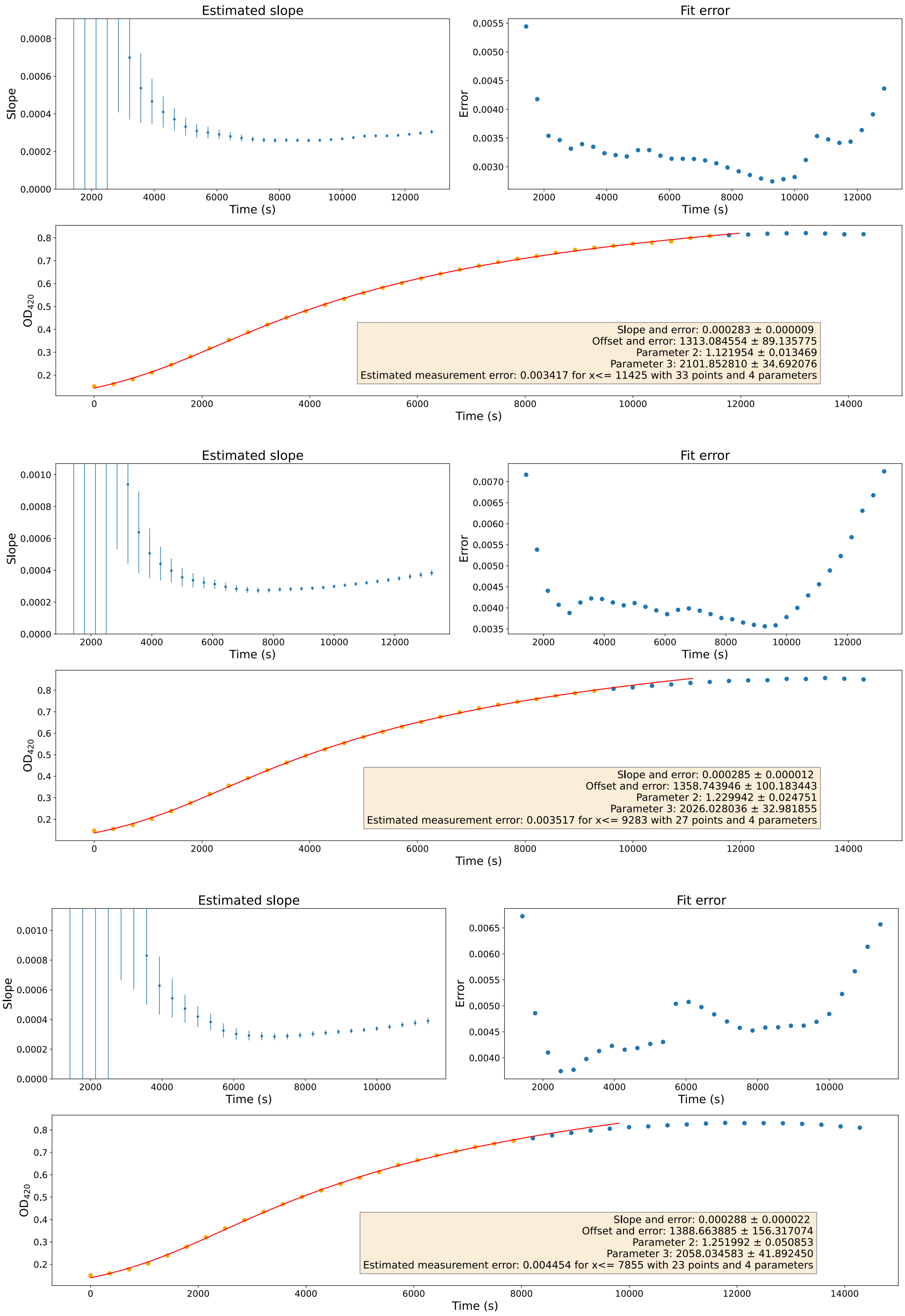


**Supplementary Figure 23: $P_{malE\_pJL1601}$ (Plate 1) Biological Replicate 2** - Estimated slopes (top left), fit errors (top right) and absorbance at 420 nm (bottom), for technical replicates 1 (top), 2 (middle) and 3 (bottom). Parameter 2: m (maximum value) and parameter 3: d (period around the offset in which the transition takes place).

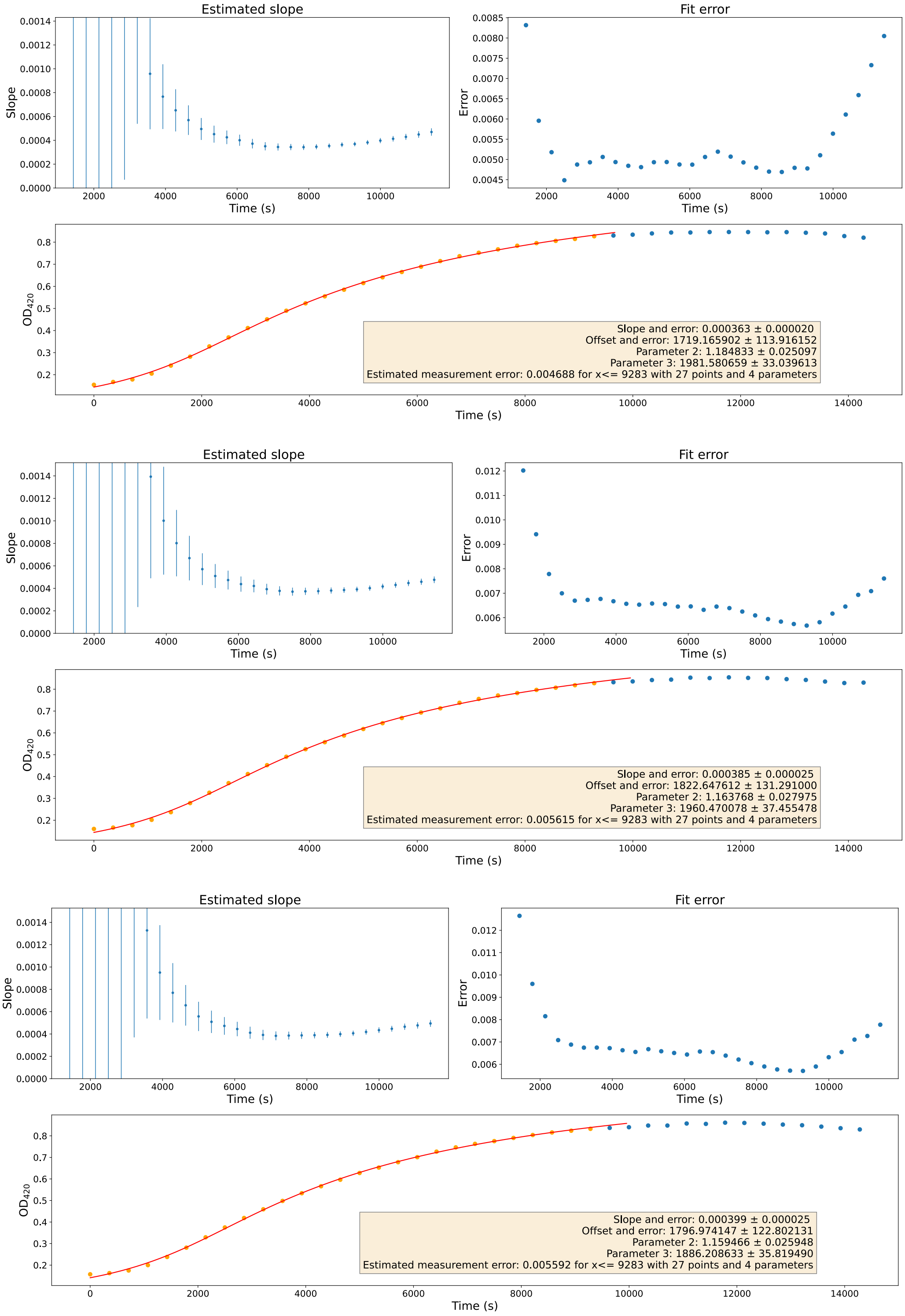


**Supplementary Figure 24: $P_{malE\_pJL1601}$ (Plate 1) Biological Replicate 3** - Estimated slopes (top left), fit errors (top right) and absorbance at 420 nm (bottom), for technical replicates 1 (top), 2 (middle) and 3 (bottom). Parameter 2: m (maximum value) and parameter 3: d (period around the offset in which the transition takes place).

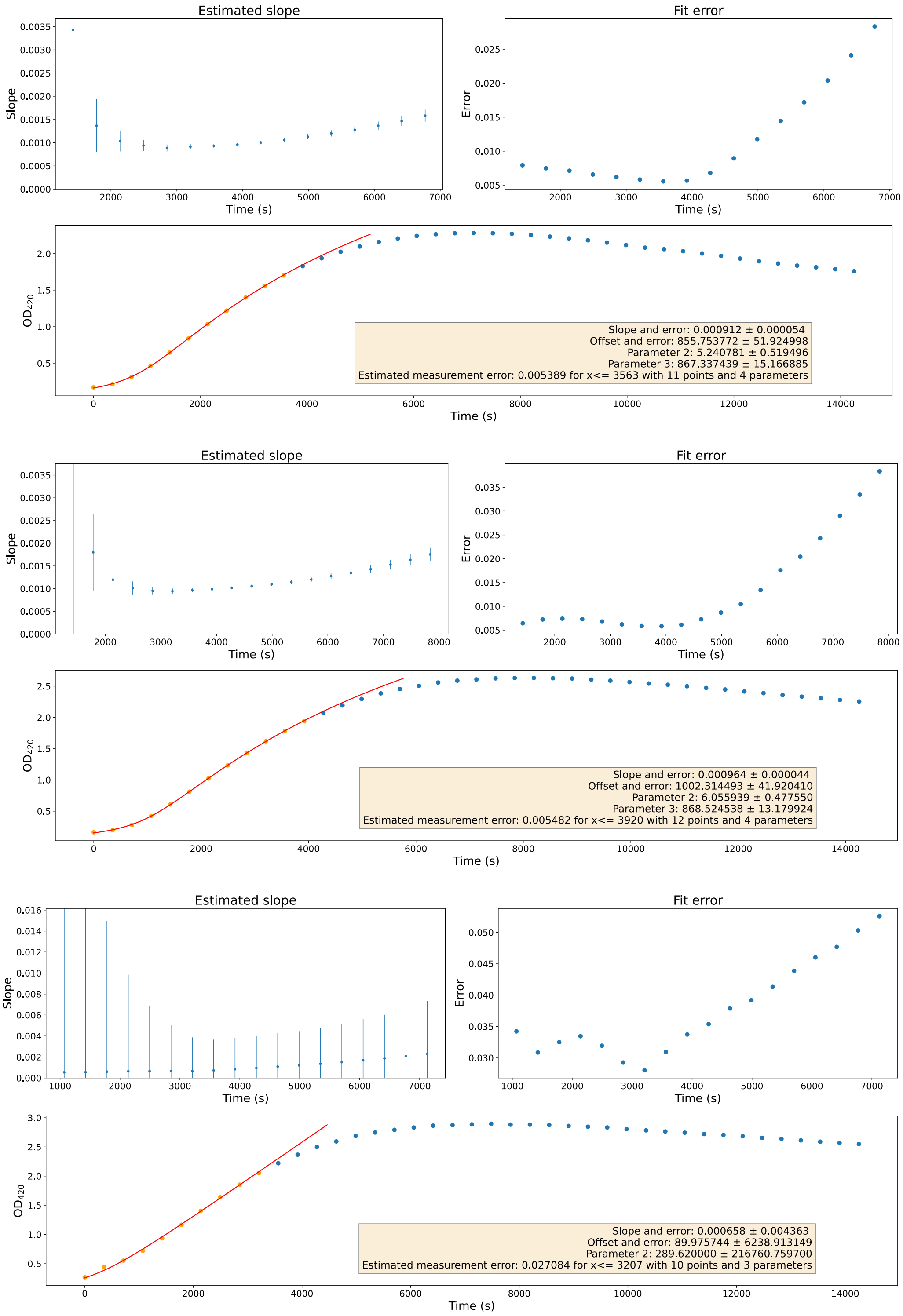


**Supplementary Figure 25: $\mathbf{P_{Saci\_2137}}$ Biological Replicate 1** - Estimated slopes (top left), fit errors (top right) and absorbance at 420 nm (bottom), for technical replicates 1 (top), 2 (middle) and 3 (bottom). Parameter 2: m (maximum value) and parameter 3: d (period around the offset in which the transition takes place).

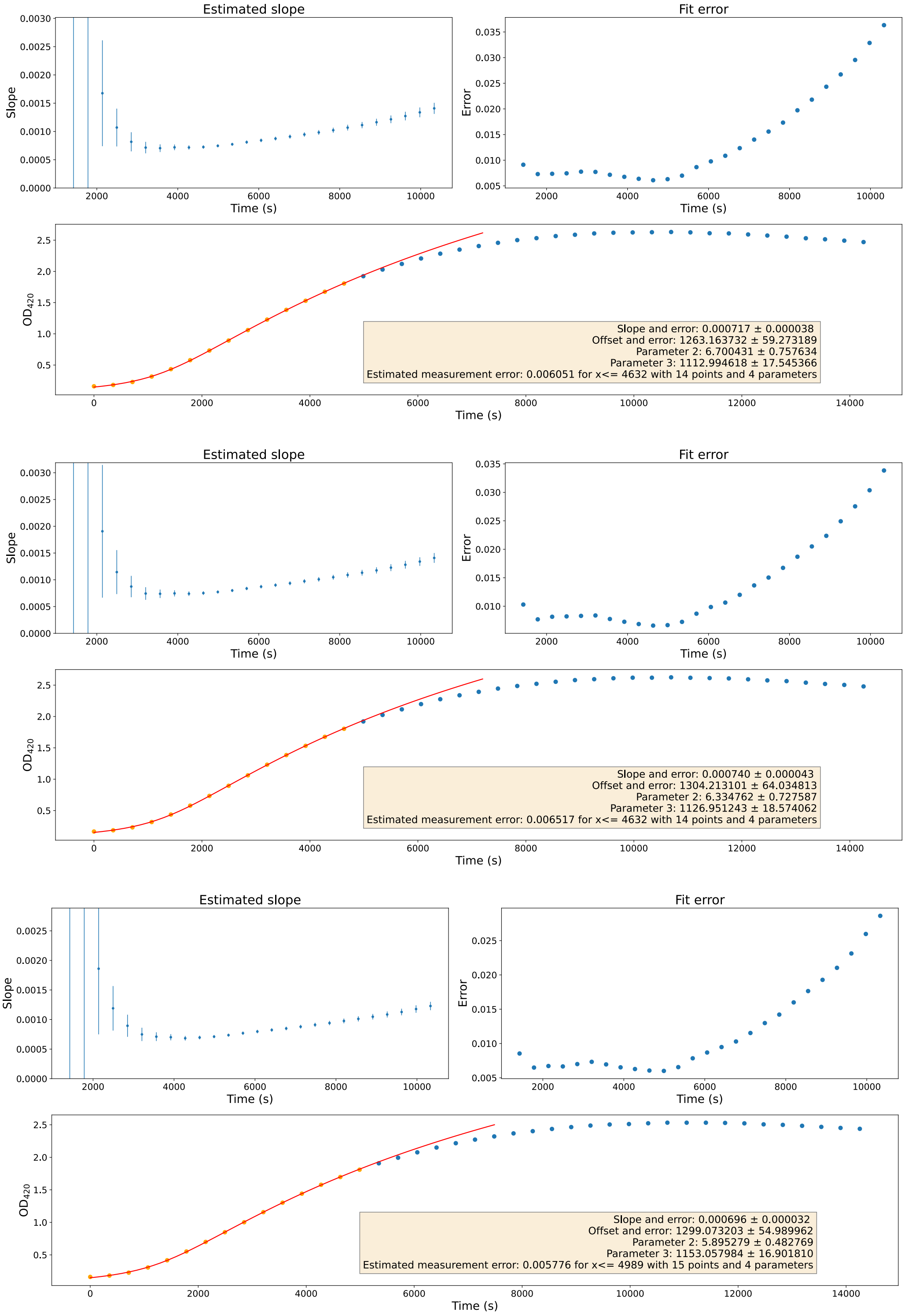


**Supplementary Figure 26: $P_{Saci\_2137}$ Biological Replicate 2** - Estimated slopes (top left), fit errors (top right) and absorbance at 420 nm (bottom), for technical replicates 1 (top), 2 (middle) and 3 (bottom). Parameter 2: m (maximum value) and parameter 3: d (period around the offset in which the transition takes place).

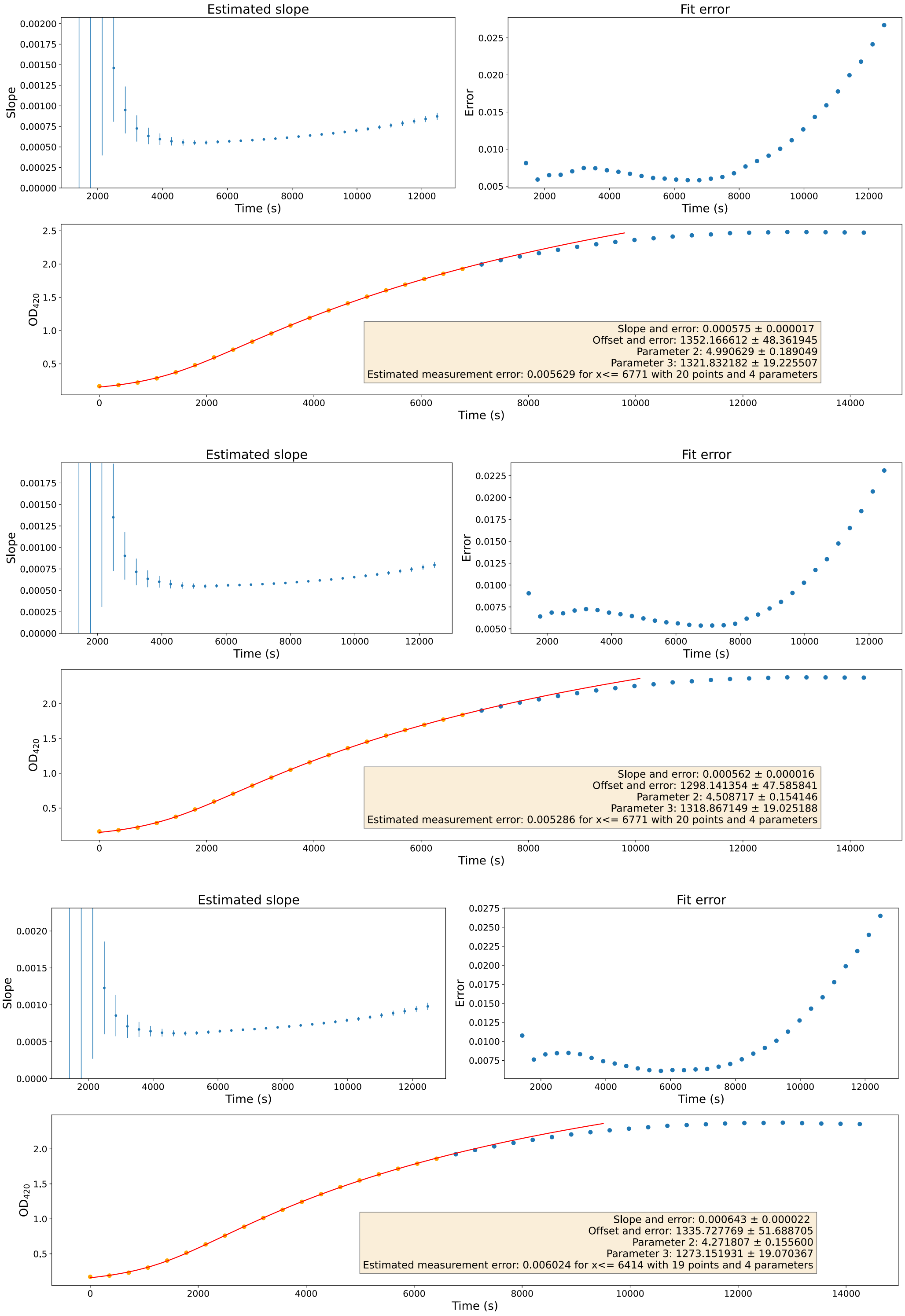


**Supplementary Figure 27: $\mathbf{P_{Saci\_2137}}$ Biological Replicate 3** - Estimated slopes (top left), fit errors (top right) and absorbance at 420 nm (bottom), for technical replicates 1 (top), 2 (middle) and 3 (bottom). Parameter 2: m (maximum value) and parameter 3: d (period around the offset in which the transition takes place).

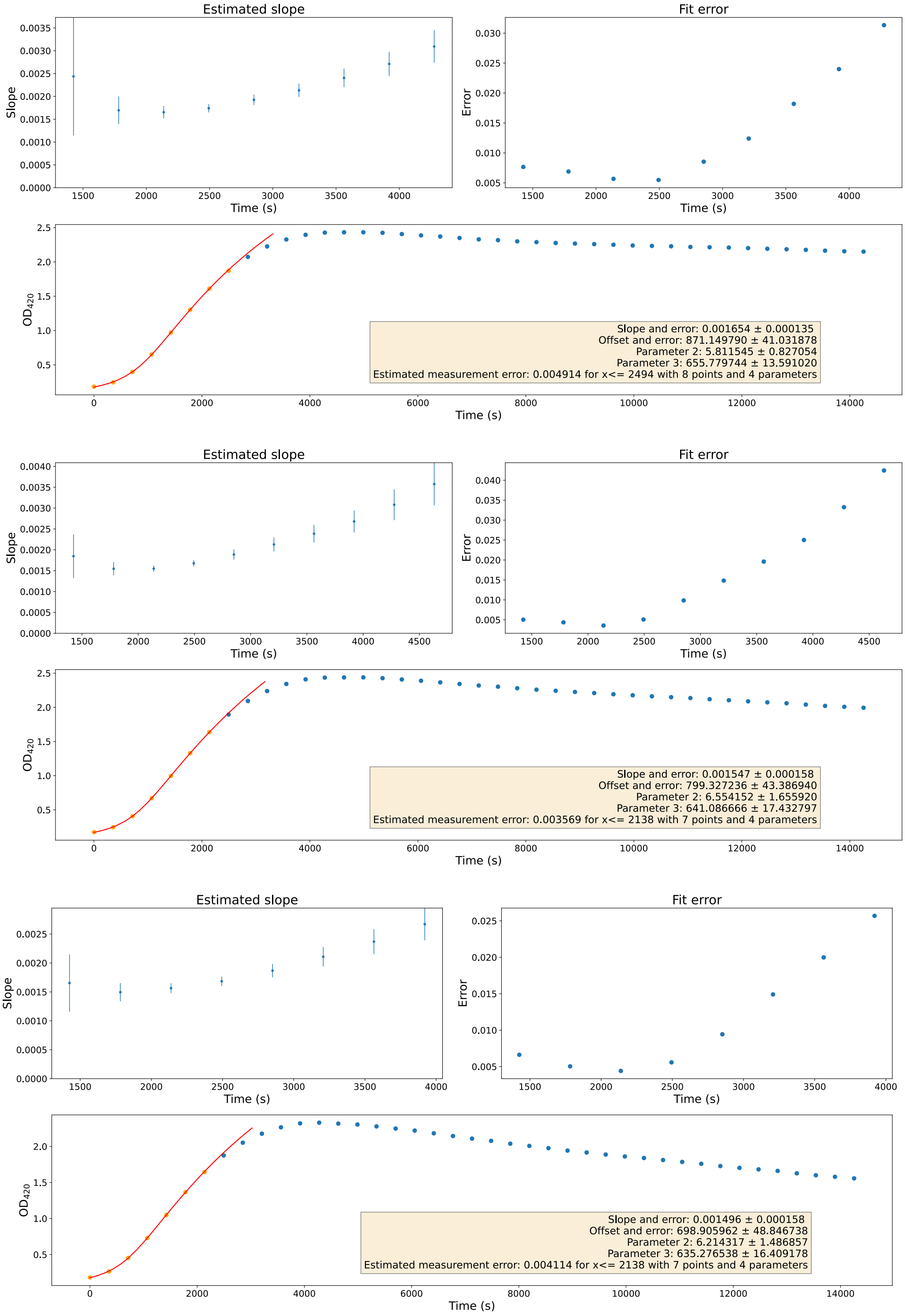


**Supplementary Figure 28: $P_{Saci\_2137core}$ Biological Replicate 1** - Estimated slopes (top left), fit errors (top right) and absorbance at 420 nm (bottom), for technical replicates 1 (top), 2 (middle) and 3 (bottom). Parameter 2: m (maximum value) and parameter 3: d (period around the offset in which the transition takes place).

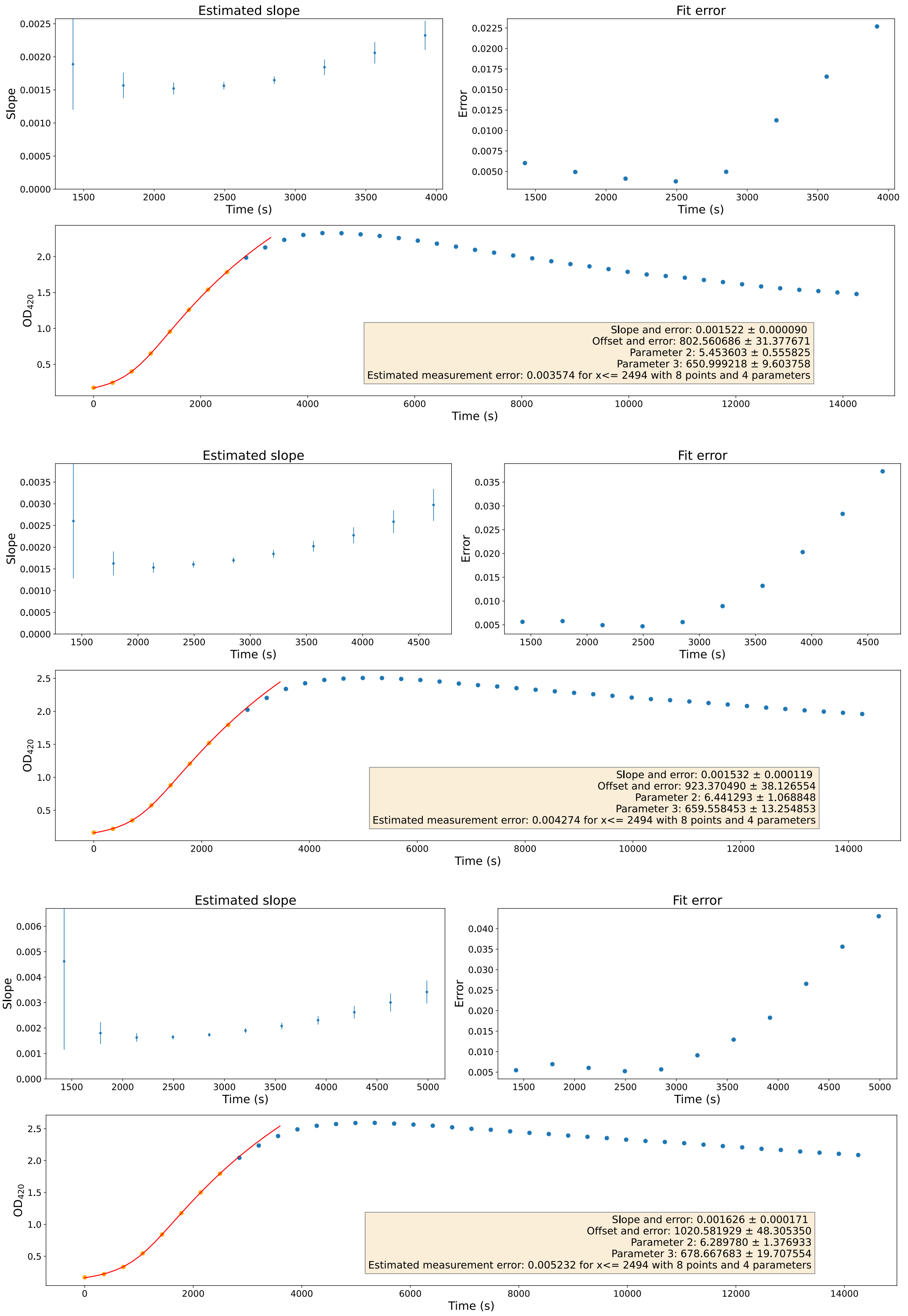


**Supplementary Figure 29: $P_{Saci\_2137core}$ Biological Replicate 2** - Estimated slopes (top left), fit errors (top right) and absorbance at 420 nm (bottom), for technical replicates 1 (top), 2 (middle) and 3 (bottom). Parameter 2: m (maximum value) and parameter 3: d (period around the offset in which the transition takes place).

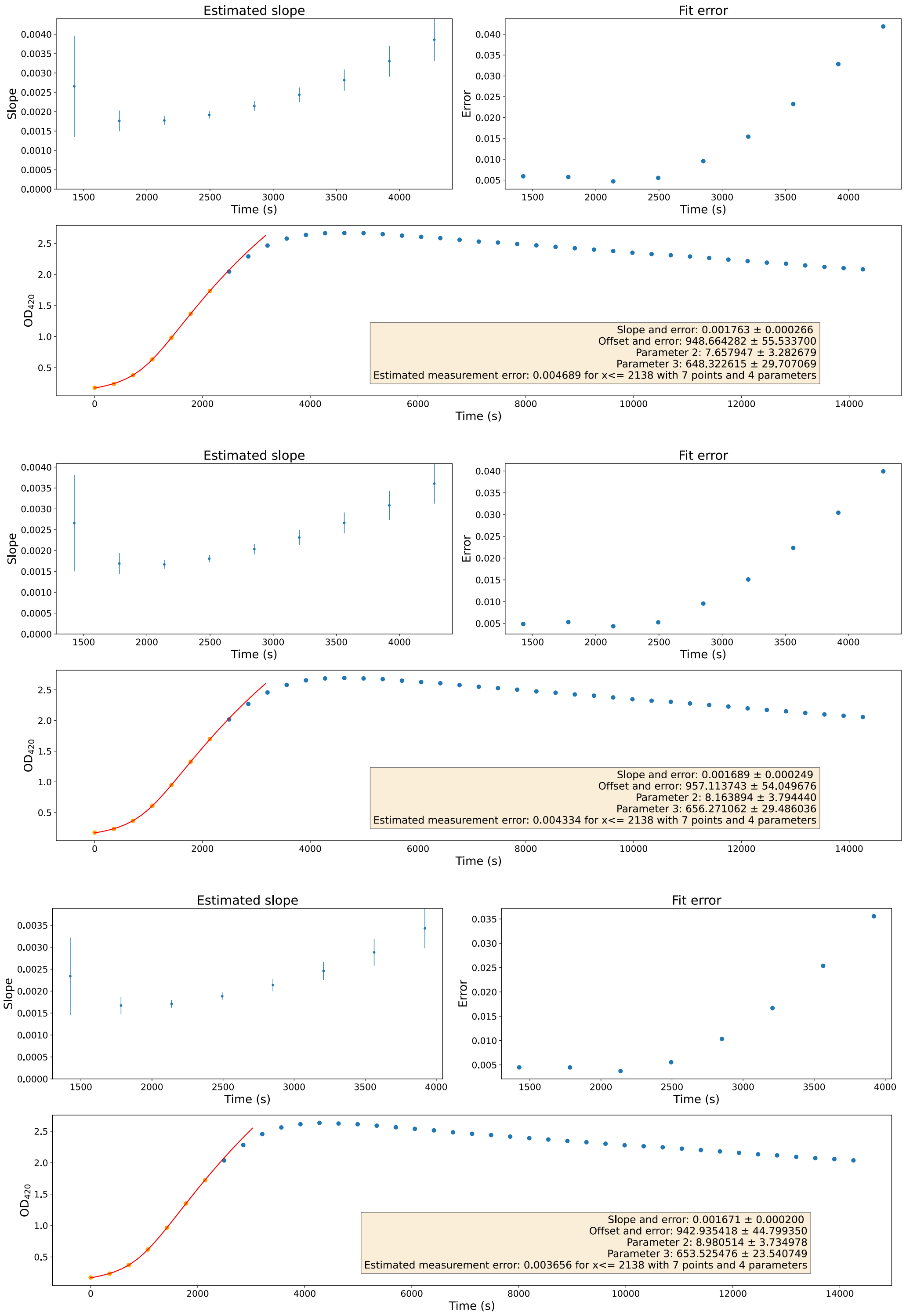


**Supplementary Figure 30: $P_{Saci\_2137core}$ Biological Replicate 3** - Estimated slopes (top left), fit errors (top right) and absorbance at 420 nm (bottom), for technical replicates 1 (top), 2 (middle) and 3 (bottom). Parameter 2: m (maximum value) and parameter 3: d (period around the offset in which the transition takes place).

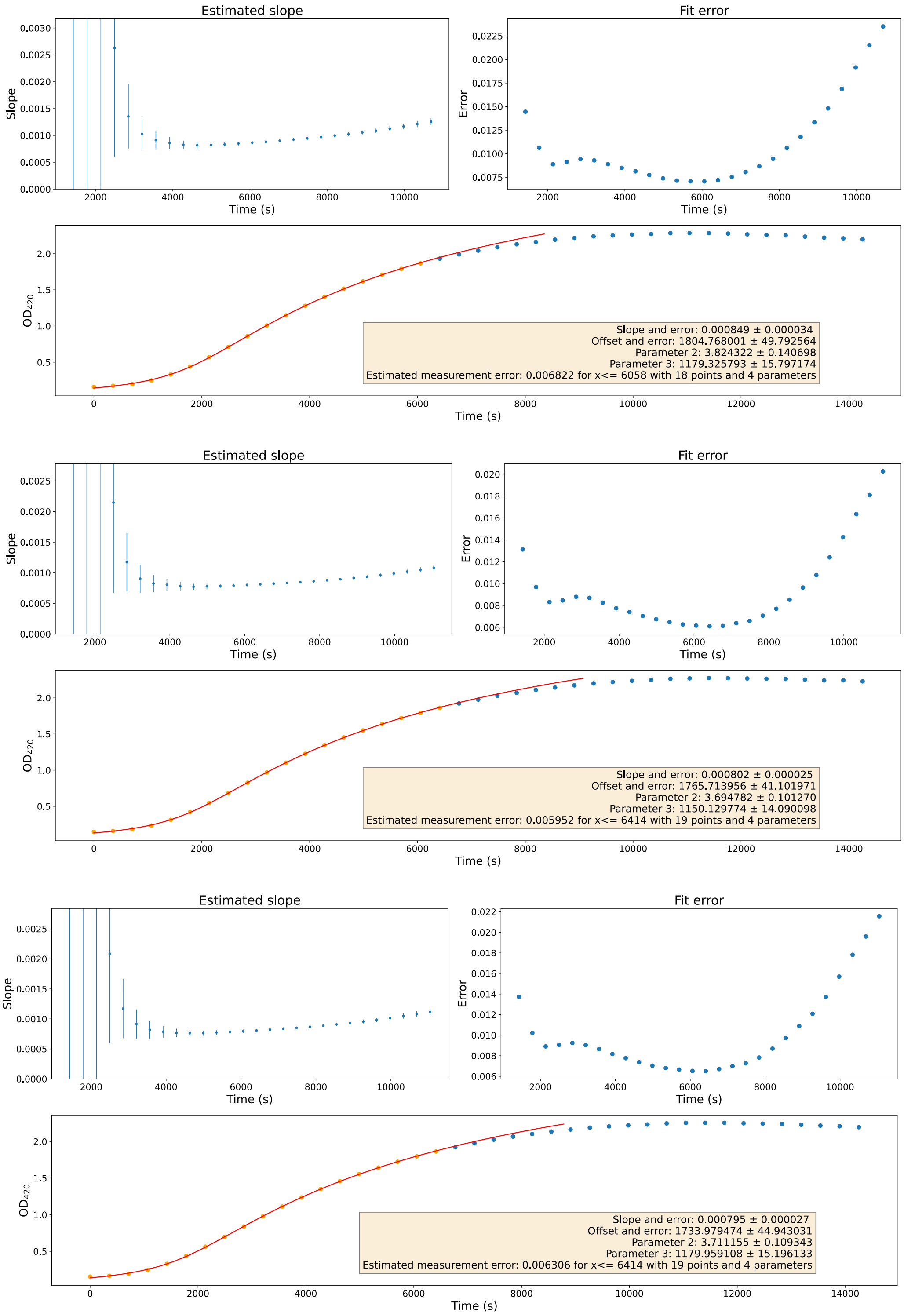


**Supplementary Figure 31: $\mathbf{P_{malE\_pSVA1450}}$ Biological Replicate 1** - Estimated slopes (top left), fit errors (top right) and absorbance at 420 nm (bottom), for technical replicates 1 (top), 2 (middle) and 3 (bottom). Parameter 2: m (maximum value) and parameter 3: d (period around the offset in which the transition takes place).

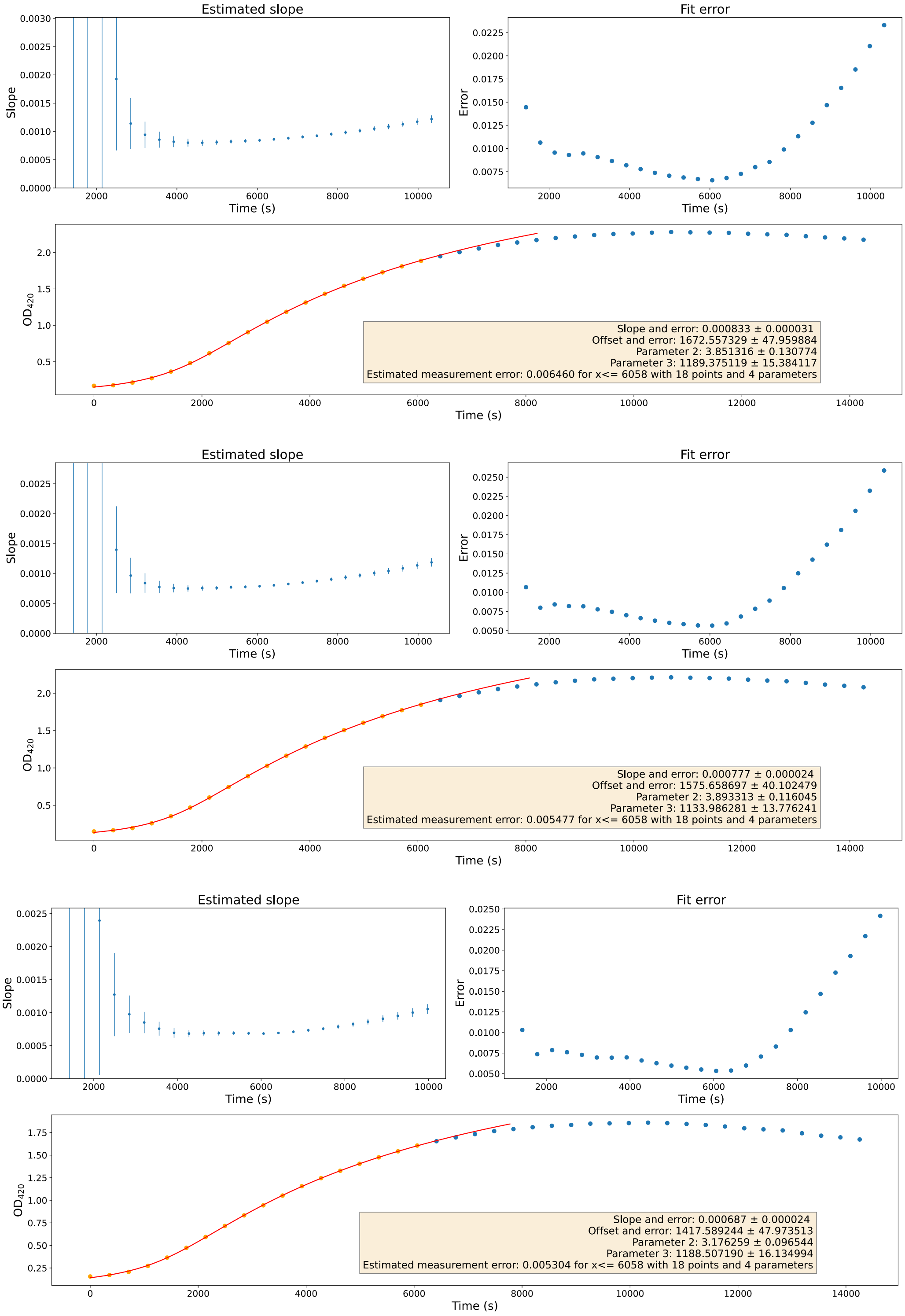


**Supplementary Figure 32: $\mathbf{P_{malE\_pSVA1450}}$ Biological Replicate 2** - Estimated slopes (top left), fit errors (top right) and absorbance at 420 nm (bottom), for technical replicates 1 (top), 2 (middle) and 3 (bottom). Parameter 2: m (maximum value) and parameter 3: d (period around the offset in which the transition takes place).

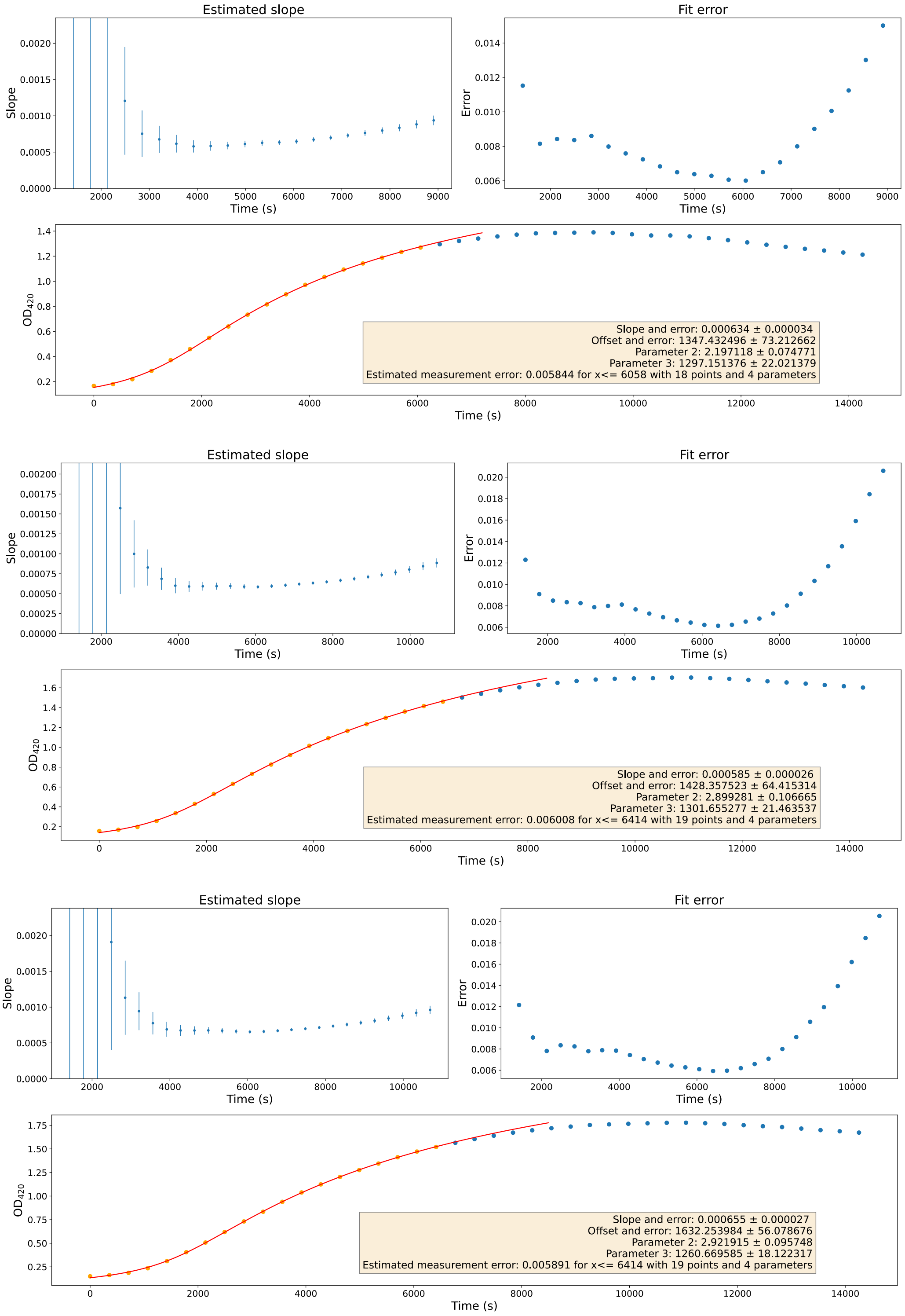


**Supplementary Figure 33: $\mathbf{P_{malE\_pSVA1450}}$ Biological Replicate 3** - Estimated slopes (top left), fit errors (top right) and absorbance at 420 nm (bottom), for technical replicates 1 (top), 2 (middle) and 3 (bottom). Parameter 2: m (maximum value) and parameter 3: d (period around the offset in which the transition takes place).

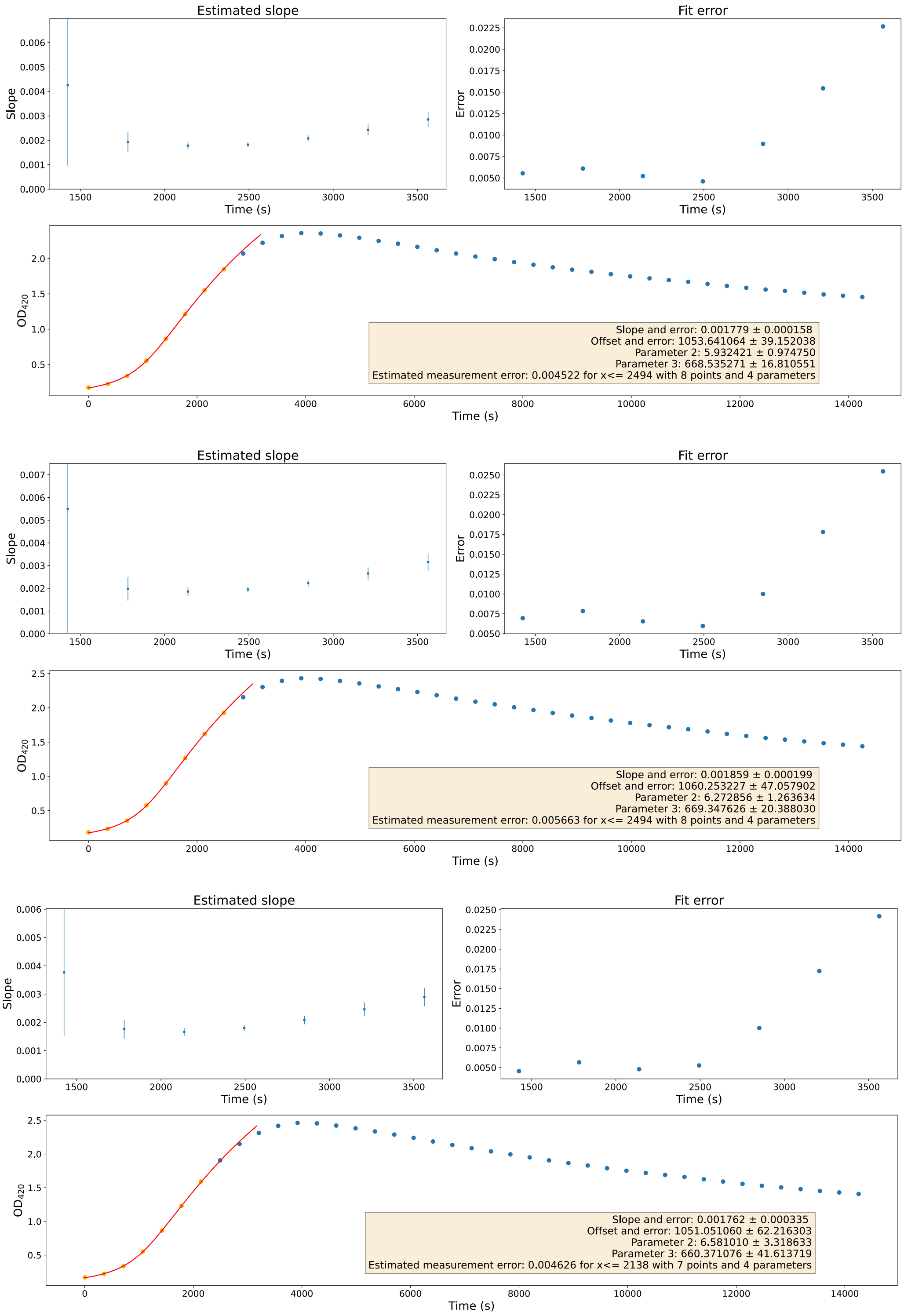


**Supplementary Figure 34: $P_{sac7d}$ Biological Replicate 1** - Estimated slopes (top left), fit errors (top right) and absorbance at 420 nm (bottom), for technical replicates 1 (top), 2 (middle) and 3 (bottom). Parameter 2: m (maximum value) and parameter 3: d (period around the offset in which the transition takes place).

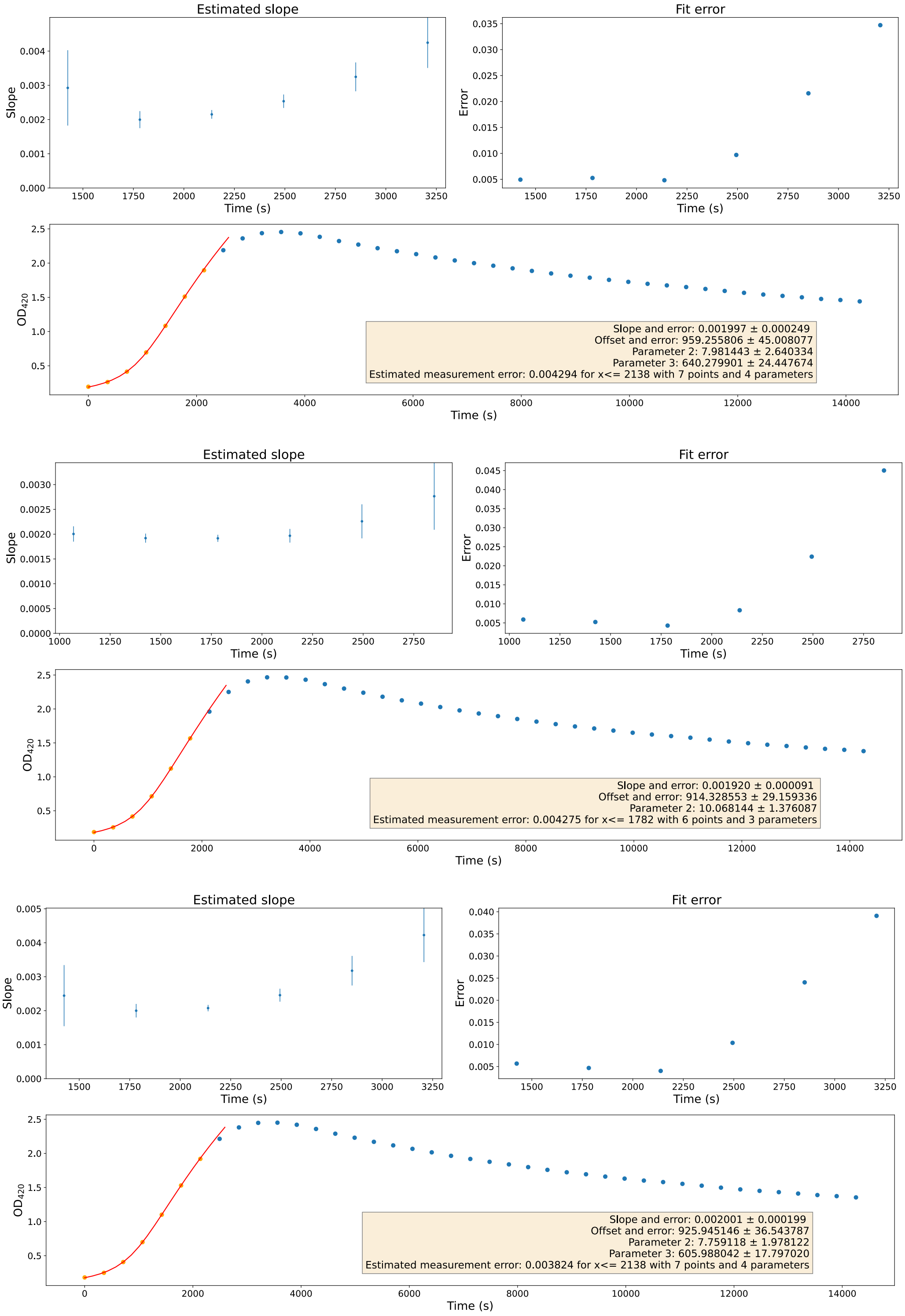


**Supplementary Figure 35: $P_{sac7d}$ Biological Replicate 2** - Estimated slopes (top left), fit errors (top right) and absorbance at 420 nm (bottom), for technical replicates 1 (top), 2 (middle) and 3 (bottom). Parameter 2: m (maximum value) and parameter 3: d (period around the offset in which the transition takes place).

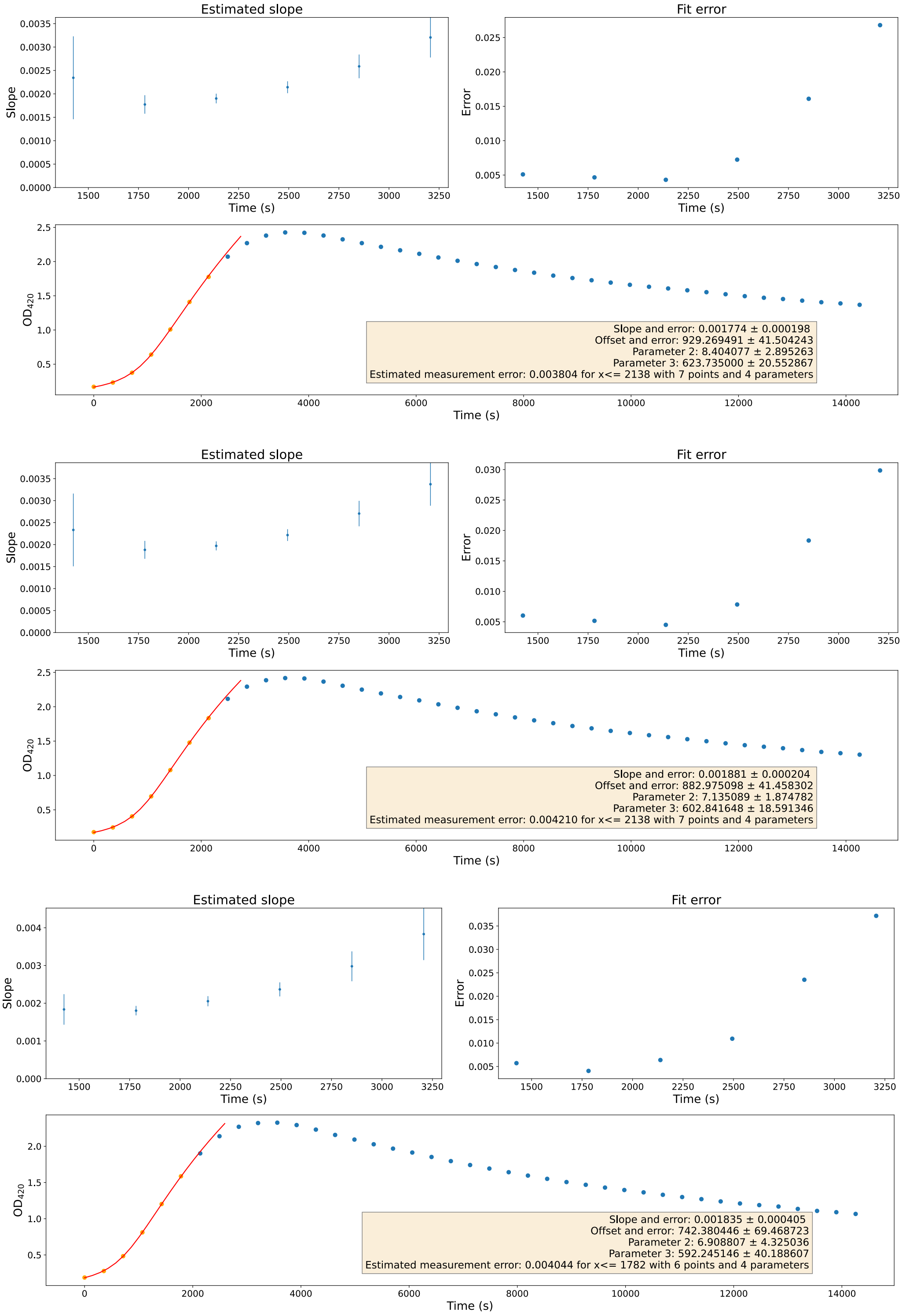


**Supplementary Figure 36: $P_{sac7d}$ Biological Replicate 3** - Estimated slopes (top left), fit errors (top right) and absorbance at 420 nm (bottom), for technical replicates 1 (top), 2 (middle) and 3 (bottom). Parameter 2: m (maximum value) and parameter 3: d (period around the offset in which the transition takes place).

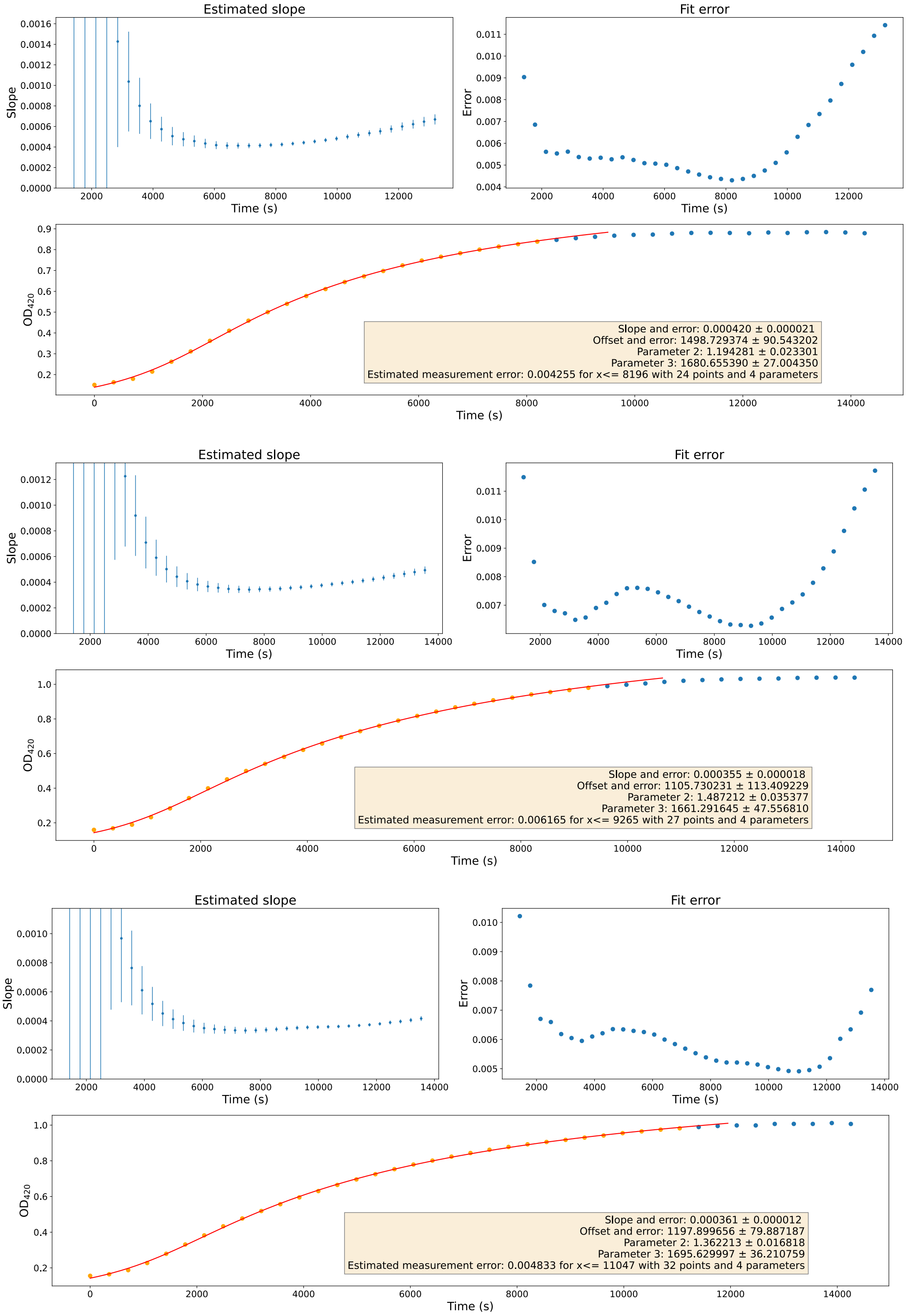


**Supplementary Figure 37: $P_{malE\_pJL1601}$ (Plate 2) Biological Replicate 1** - Estimated slopes (top left), fit errors (top right) and absorbance at 420 nm (bottom), for technical replicates 1 (top), 2 (middle) and 3 (bottom). Parameter 2: m (maximum value) and parameter 3: d (period around the offset in which the transition takes place).

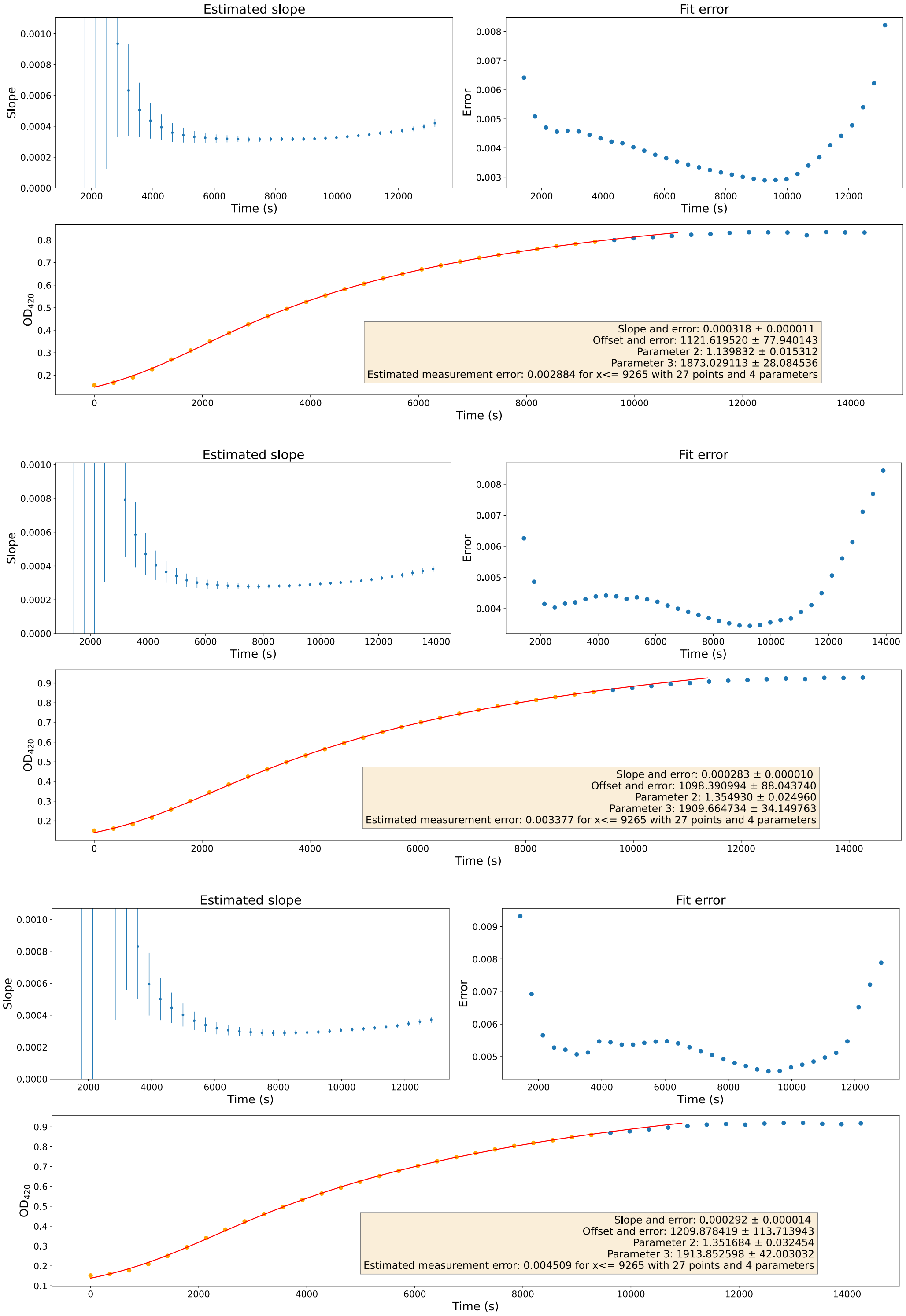


**Supplementary Figure 38: $P_{malE\_pJL1601}$ (Plate 2) Biological Replicate 2** - Estimated slopes (top left), fit errors (top right) and absorbance at 420 nm (bottom), for technical replicates 1 (top), 2 (middle) and 3 (bottom). Parameter 2: m (maximum value) and parameter 3: d (period around the offset in which the transition takes place).

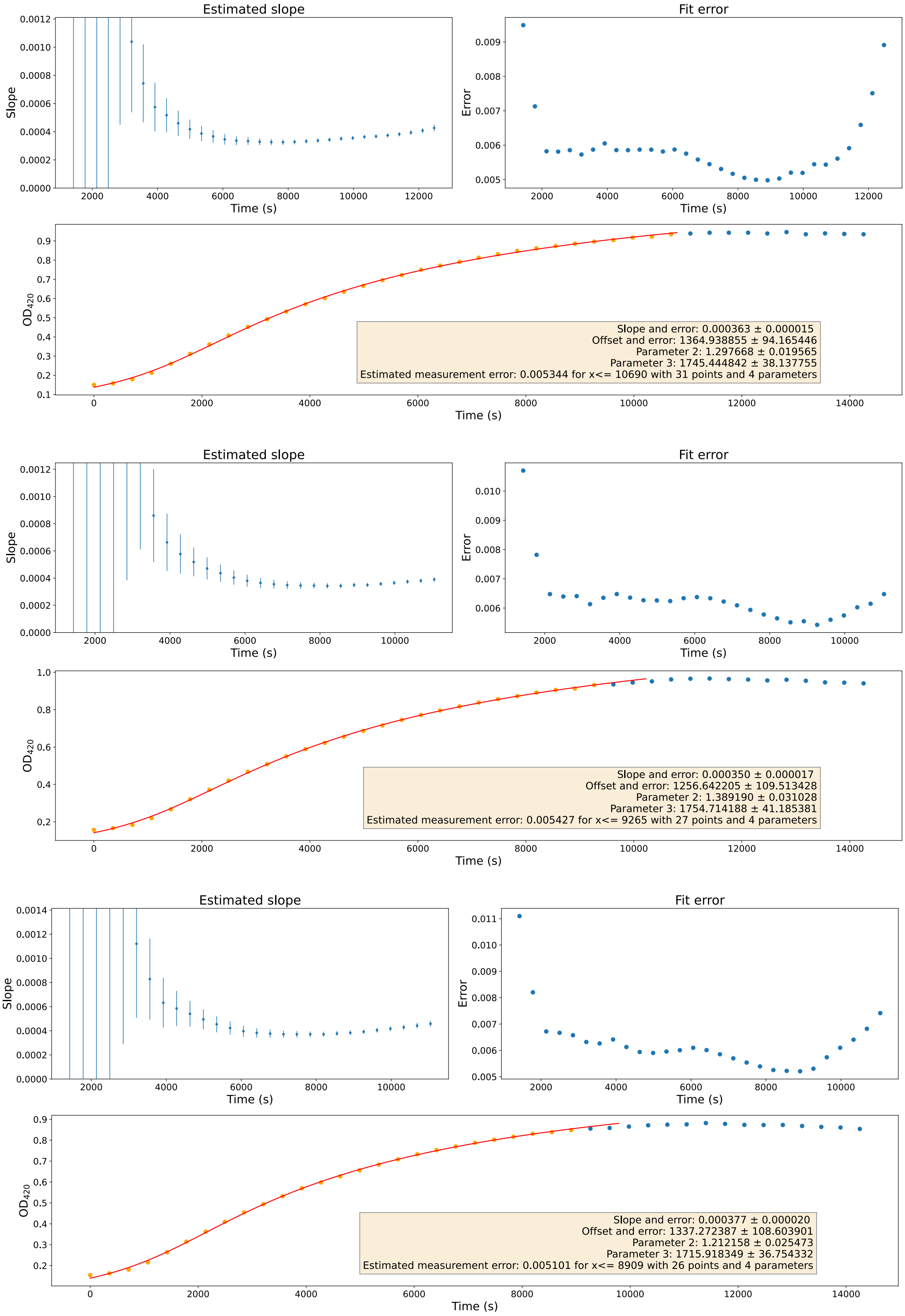


**Supplementary Figure 39: $P_{malE\_pJL1601}$ (Plate 2) Biological Replicate 3** - Estimated slopes (top left), fit errors (top right) and absorbance at 420 nm (bottom), for technical replicates 1 (top), 2 (middle) and 3 (bottom). Parameter 2: m (maximum value) and parameter 3: d (period around the offset in which the transition takes place).
